# Patterning and growth are coordinated early in the cell cycle

**DOI:** 10.1101/2022.09.22.508753

**Authors:** Cara M. Winter, Pablo Szekely, Heather Belcher, Raina Carter, Matthew Jones, Scott E. Fraser, Thai V. Truong, Philip N. Benfey

## Abstract

Precise control of cell division is essential for proper patterning and growth during the development of multicellular organisms. Coordination of formative (asymmetric) divisions that generate new tissue patterns with proliferative (symmetric) divisions that promote growth is poorly understood. Here, we employed quantitative 4D light sheet and confocal microscopy to probe in vivo the dynamics of two transcription factors, SHORTROOT (SHR) and SCARECROW (SCR), which are required for asymmetric division in the stem cell niche of Arabidopsis roots [1,2]. Long-term (up to 48 hours), frequent (every 15 minutes) imaging of the two regulators in tandem in single cells, in conjunction with a SHR induction system, enabled us to challenge an existing bistable model[3] of the SHR/SCR gene regulatory network. By directly controlling SHR and SCR expression dynamics, we were able to identify key features that are essential for rescue of asymmetric division in *shr* mutants. We show that instead of high stable levels of nuclear SHR and SCR, only low transient levels of expression are required. Nuclear SHR kinetics do not follow predictions of the bistable model, and the regulatory relationship between SHR and SCR can be modeled by monostable alternatives. Furthermore, expression of these two regulators early in the cell cycle determines the orientation of the division plane, resulting in either formative or proliferative cell division. Our findings provide evidence for an uncharacterized mechanism by which developmental regulators directly coordinate patterning and growth.

## Introduction

Control of cell division is essential for the development and survival of multicellular organisms. The final size, shape, and function of tissues hinge upon the coordination of formative divisions that generate new cell types with proliferative divisions that increase cell number and contribute to growth [4]. Due to intrinsic and extrinsic cell polarity, a 90-degree rotation of the division plane determines whether a cell will divide asymmetrically (producing daughter cells with different fates) or symmetrically (producing daughter cells with similar fates) [5,6]. The wrong choice can lead to over proliferation of cells, resulting in aberrant morphogenesis or tumorigenesis [7–9]. Developmental regulators that specify cell fate and directly interface with the cell cycle machinery [10–12] are likely arbiters of this decision, but challenges in long-term, multicolor imaging in multicellular organisms have limited knowledge about how these regulators dynamically control cell division in situ. Here, we use long-term imaging, mathematical modeling, and quantitative analysis to show how two developmental regulators coordinate the choice between asymmetric and symmetric division in the stem cell niche of the Arabidopsis root.

The transcription factors SHORTROOT (SHR) and SCARECROW (SCR) control the formative asymmetric division in the Arabidopsis root that gives rise to the endodermis and cortex cell types (ground tissue). This division occurs in the immediate progeny of a stem cell, the cortex/endodermal initial cell daughter (CEID) cell (Fig. S5a). SHR, a mobile intercellular signaling molecule, moves from the central tissues of the root into the endodermis and CEID where it activates SCR [12,13–17]. SHR and SCR together activate the cell cycle regulator CYCLIND6 (CYCD6) only in the CEID, triggering asymmetric division [18]. In *shr* and *scr* mutants, asymmetric division of the CEID cell does not occur, resulting in a single ground tissue layer, rather than distinct files of endodermis and cortex cells (Fig. S5b).

Cruz-Ramirez et al.[3] proposed a bistable model to explain both how and where SHR and SCR trigger the decision to divide. According to the model, high stable steady states of SCR and nuclear SHR are necessary to trigger asymmetric division. These high steady states are reached specifically in the CEID due to a switch-like response to a SHR gradient on the radial axis and an auxin gradient on the longitudinal axis that reaches a maximum in the stem cell niche. In this model, high auxin levels in the CEID promote SHR/SCR-dependent expression of CYCD6, which phosphorylates RETINOBLASTOMA RELATED1 (RBR) and releases SCR from RBR sequestration. This allows a SCR autoregulatory loop to engage. The increasing levels of SCR amplify levels of nuclear SHR by retaining it in the nucleus, leading to high levels of both SCR and nuclear SHR, and the CEID is robustly committed to division through this bistable switch.

The concept of bistability was first introduced more than 60 years ago by Monod and Jacob [19], who proposed that the structure of a gene regulatory network can translate reversible molecular interactions into an irreversible, binary decision. Specifically, the presence of positive feedback loops can resuit in a bistable system that can switch between on and off states [20]. Bistability is at the heart of mathematical models of the dynamics of decision-making in many systems [21,22]. Bistable switches enable directional, abrupt and stable transitions between cellular states or can translate a graded signal into a binary output [23–27]. For example, in Drosophila embryos, the sharp boundary of hunchback (hb) expression has been modeled as a bistable switch involving a gradient of Bicoid (Bcd) and a hb auto-regulatory positive feedback loop [28]. A mathematical model of the cell cycle from measurements in cultured mammalian cells suggests that irreversible commitment to cell division occurs through a bistable switch involving retinoblastoma (RB) and E2F [29].

Mathematical models are often based on protein-protein and protein-DNA interactions derived from “snapshot” data taken at single points in time from whole tissues. However, the specific molecular interactions of a gene regulatory network and behavior of particular proteins may vary from cell to cell in a multicellular organism depending on the presence or absence of binding partners and chromatin state [30,31]. Furthermore, positive feedback does not always lead to bistability [32], and other decision-making mechanisms exist in addition to bistability. For example, the simple presence of a factor at the right place and right time can be sufficient to alter the cell cycle program and commit a cell to an alternate fate [12]. Thus, there is a need for experimental inquiry into gene regulatory network dynamics in an in vivo context for an accurate understanding of the molecular mechanisms underlying the specification of asymmetric division [31].

Quantitative time-lapse imaging of transcription factor dynamics has provided key insights into gene regulatory network function in single cell organisms and mammalian cell lines [33–36]. For example, through observation of PU.1 and GATA1 dynamics in mouse cell lines, Hoppe et al. [37] challenged an existing model that early myeloid lineage choice is determined by random fluctuations of these mutually antagonistic transcription factors. Studies using MS2 and similar systems allow for the imaging of transcription factor dynamics at the level of single molecules [38,39]. However, phototoxicity and photobleaching restrict all these studies using confocal microscopy to short timescales or infrequent sampling, limit the number of fluorophores that can be imaged simultaneously, and have made studies of network dynamics in vivo elusive.

To optimally probe the dynamics of a transcription factor network and examine the regulatory relationships between constituents, at least two transcription factors need to be observed in tandem on a long timescale [31]. If a marker is included for normalization purposes, then at least three fluorophores need to be imaged simultaneously. Due to its lower phototoxicity, light sheet microscopy provides the means for longer-term multi-color imaging of transcription factor dynamics. Its potential for the examination of gene expression dynamics in vivo has been extolled for nearly two decades, but the technology has primarily been used for observation of cellular dynamics and morphology changes during development [40–42].

In this work, we employ quantitative 4D imaging of living roots to gain insight into the dynamics of the SHR/SCR gene regulatory network that controls asymmetric division. We built a light sheet microscope customized for root tracking and imaging, enabling simultaneous 3D observation of multiple regulators over long timescales. Utilizing an inducible SHR line to rescue asymmetric divisions in the *shr* mutant, and various induction regimens, we generated and quantified a variety of SHR and SCR expression profiles in over 1,000 cells for up to 48 hours after induction. We used these profiles to identify key features of SHR and SCR dynamics required to specify asymmetric division and to investigate bistability as the underlying mechanism. We model the regulatory relationship between SHR and SCR using a few simple monostable ODE models, indicating that bistability is not needed to explain the dynamics that we observe. We also show that, in contrast to a key prediction of high “locked” nuclear SHR and SCR levels, transient expression of both regulators is capable of triggering asymmetric division at a fraction of the concentration at which they will ultimately be expressed.

Finally, our in vivo measurements revealed a key aspect that was missing from the existing model: namely that SHR and SCR levels are interpreted within the context of the cell cycle. We propose that low threshold levels of SHR and SCR act early in the cell cycle to trigger an alternate cell cycle program, changing the orientation of the division plane. This model successfully predicts asymmetric cell division up to 94% of the time.

## Results

### Long-term *in vivo* confocal imaging links inducible SHR dynamics to asymmetric division

To observe and quantify the dynamics of SHR in vivo, we generated a fluorescently-labeled inducible SHR construct, SHR:GAL4-GR UAS:SHR-GFP, capable of rescuing the asymmetric divisions absent in *shr2* mutants [1,2]. Compared with the small number and infrequent divisions of CEID cells in wild-type roots, the numerous divisions induced in single roots with this SHR induction system made it feasible to quantify SHR expression in sufficient numbers of cells for analysis. Into this background we introduced a construct, 35S:H2B-RFP, to uniformly label nuclei in the root. We induced expression of *SHR-GFP* mRNA in its endogenous expression domain by treating SHR:GAL4-GR UAS:SHR-GFP 35S:H2B-RFP *shr2* plants with diminishing concentrations of dexamethasone (dex) starting from maximal dex activity [43] (10, 1, 0.05, 0.03, 0.02 and 0.01 μM), and imaged the roots in 3D every 15 minutes for up to 24 hours using confocal microscopy (Fig. 1a,b; Extended Data Videos 1-4). We quantified the relative levels of SHR protein (see Methods) present in ground tissue cell lineages (N = 935 from 29 roots) from the time of induction up to asymmetric division or the end of the experiment if no division occurred (Fig. 1c,d; Supplementary Data Table 1).

**Fig. 1.**
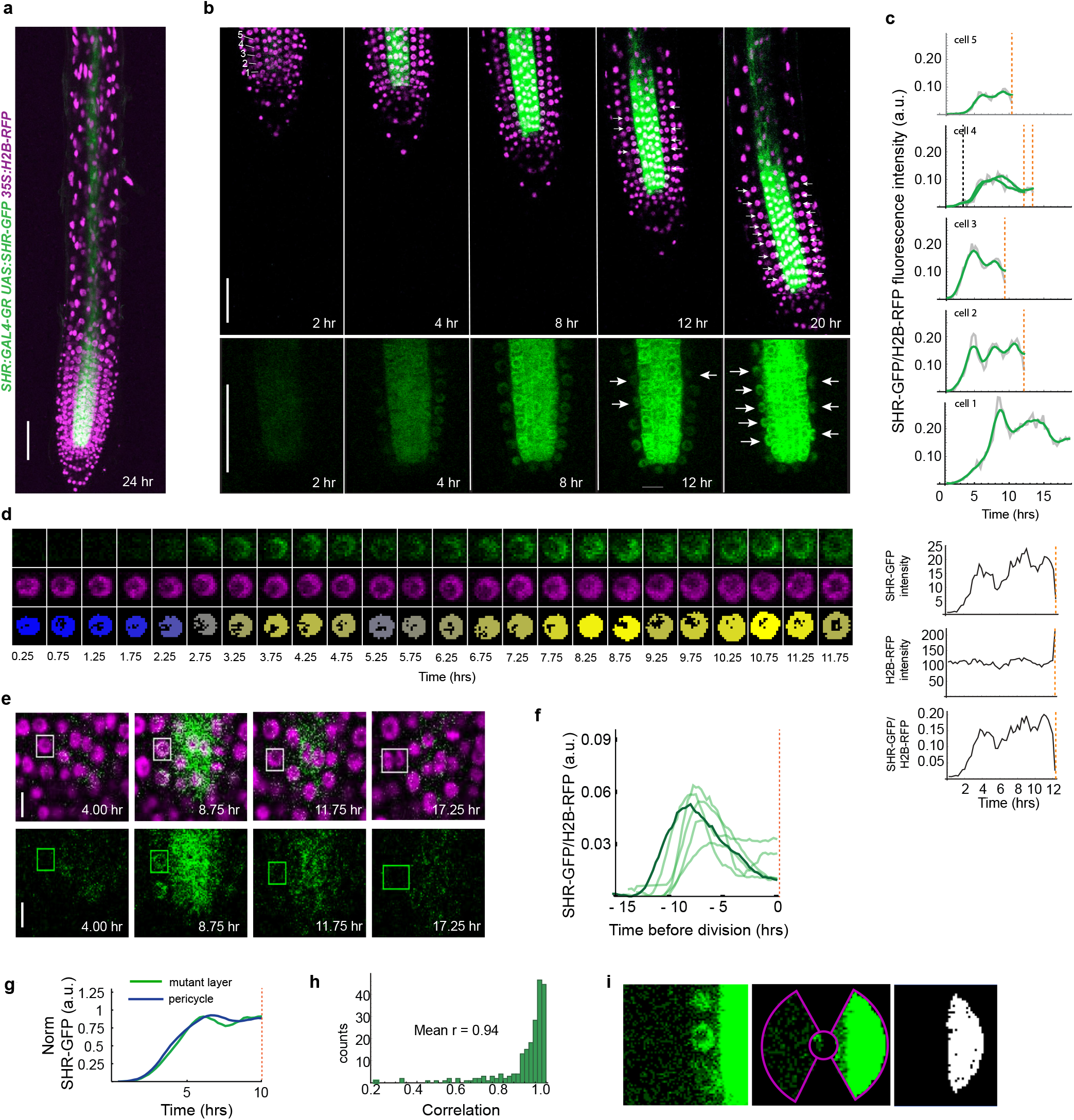
Long-term 4D confocal imaging of SHR reveals dynamics are inconsistent with bistability. Maximum projection (a) and median longitudinal sections (scale bar - 50 μm) (b) at various timepoints after induction with dex. Green, SHR-GFP; Magenta, H2B-RFP; Top, SHR-GFP and H2B-RFP; Bottom, SHR-GFP only Gamma is set to 0.75 to show signal in the mutant layer. Numbers at top left, first five cell positions in mutant ground tissue (scale bar - 50 μm). c) Raw (grey) and smoothed (green) relative fluorescence intensity of SHR-GFP over time in the first five cells of a single cell file after full induction (10 μM dex). d) Raw images of SHR-GFP (green), H2B-RFP (magenta), and Otsu-mask colored by SHR-GFP/H2B-RFP signal of a representative cell over time. Right, corresponding quantified trajectories. e) Median longitudinal sections through a root tip treated with low dex (scale bar – 10 μm). f) SHR trajectories for asymmetrically dividing cells treated with low dex that show a low early peak of expression. Dark green, SHR trajectory corresponding to images in e. g) SHR trajectories over time for a representative mutant layer cell (green) and its nearest pericycle region (blue). h) Histogram of correlations between each cell’s SHR trajectory in the ground tissue and its nearest pericycle region. r, Pearson correlation coefficient; n = 206 cells from 28 roots. i) Images illustrate the method used to extract pericycle data. Left: image of a ground tissue nucleus and its nearest pericycle region expressing SHR. Middle: result of applying localization mask (see Methods) to image. Right: final binary mask used to extract pericycle pixel intensities. Black dashed line, symmetric division; Orange dashed line, asymmetric division.

Rescuing asymmetric divisions in the *shr2* mutant layer with an inducible SHR has been used previously to identify a key regulator of CEID division [18]. These divisions can be induced in young plants (5 days post-germination), are dependent on SCR (Fig. S5c,d), and arise from the *shr* mutant layer, which exhibits an expression profile with few characteristics of endodermal identity [44]. Thus, they are likely to be controlled by the SHR/SCR pathway active in the stem cell niche and not the SCR-independent pathway controlling asymmetric divisions of the endodermis that give rise to additional cortex layers later in development [45–48].

SHR expression was induced quantitatively in response to different levels of dex (Fig. S6a-c). At the highest concentration (10 μM), we observed maximal expression of SHR-GFP protein in the stele and pericycle, followed by movement into the *shr2* mutant ground tissue layer and asymmetric divisions along the length of the meristem [18] (Fig. 1a,b; Extended Data Videos 1-4). Divisions occurred only in the meristem, and occasionally, a cell divided anticlinally (symmetrically) before the periclinal asymmetric division (see cell 4, Fig. 1c). Cells directly adjacent to the quiescent center (QC) (Fig. S5a,b) in position 1 divided later and at a lower frequency than more shootward cells (Fig. S6d,e). Since QC cells can signal to adjacent cells to maintain them in a non-dividing, undifferentiated state [49], we removed these cells from subsequent analyses. Decreasing levels of dex resulted in lower induction of SHR-GFP and fewer dividing cells (Fig. S6a-c,f,g). After 24 hours of induction, we observed near complete rescue of meristematic asymmetric divisions at 1 μM and 10 μM dex (96 and 99 percent, respectively; Fig. S6g) and no asymmetric divisions at a concentration of 0.01 μM dex.

The bistable model postulates that SHR triggers asymmetric division when nuclear SHR levels in the ground tissue “flip” to a high steady state (Fig. S6h). This rapid increase in nuclear SHR, relative to a steady influx and increase in cytosolic SHR, is due to sequestration of SHR by increasing levels of SCR in the ground tissue nuclei [3]. Consistent with this model, in many cases we observed a rapid increase in SHR levels followed by a period during which higher SHR levels were relatively constant prior to division (Fig. 1c). However, in other cases, a transient low peak of SHR expression was able to trigger division many hours later (Fig. 1e,f, Extended Data Videos 5 and 6).

In addition, we found that the kinetic profile of nuclear SHR in the ground tissue nuclei was closely correlated with the SHR expression profile in the nearest pericycle cells (from which SHR moves into the ground tissue cytosol [13,50,51])(r = 0.94, Fig. 1g-i). This suggests that movement of SHR and accumulation in the nucleus is driven primarily by concentration differences between the two cell files. Thus, we could find no evidence that levels of nuclear SHR are regulated as predicted by the model.

Taken together, the nuclear SHR kinetics we measured are inconsistent with a bistable model in which a flip to high steady state levels of nuclear SHR and SCR is necessary to trigger division. To explore alternative models and better understand the mechanism by which these regulators control asymmetric division, we sought next to directly examine SHR regulation of SCR expression.

### Long-term in vivo 4D light sheet imaging of SHR and SCR dynamics in tandem reveals bistability is not needed

We measured the kinetics of SHR and SCR expression directly and simultaneously in single nuclei, using a light sheet microscope that we built and customized to image and track growing root tips (see Methods, Fig. S7a-c). This microscope is capable of imaging multiple fluorophores in 3D over a long period of time with reduced phototoxicity and photobleaching under near-physiological conditions [52] (Extended Data Video 7). We first introduced SCR:SCR-mKATE2 and UBQ10:H2B-CFP constructs into the SHR:GAL4-GR UAS:SHR-GFP *shr2* background. Next, we induced transcription of SHR-GFP and subsequent activation of SCR-mKATE2 by treating SHR:GAL4-GR UAS:SHR-GFP *shr2* SCR:SCR-mKATE2 UBQ10:H2B-CFP plants with dex (see Methods). We used different concentrations of dex (40 μM, 20 μM and 0.4 μM) as either a continuous treatment or as pulses (see Methods) to obtain a range of SHR and SCR expression profiles. We imaged and quantified the levels of SHR and SCR in the ground tissue every 15 min for up to 48 hours, from the time of induction up to the time of asymmetric division or the end of the experiment (Fig. 2a-e; Extended Data Videos 8 and 9, Supplementary Data Table 2). In roots exposed to constant levels of 40 μM dex (full induction), cells began dividing asymmetrically around 13 hours after SHR induction. Excluding the cell closest to the QC, 89% of the meristematic cells had divided after 45 hours.

**Fig. 2.**
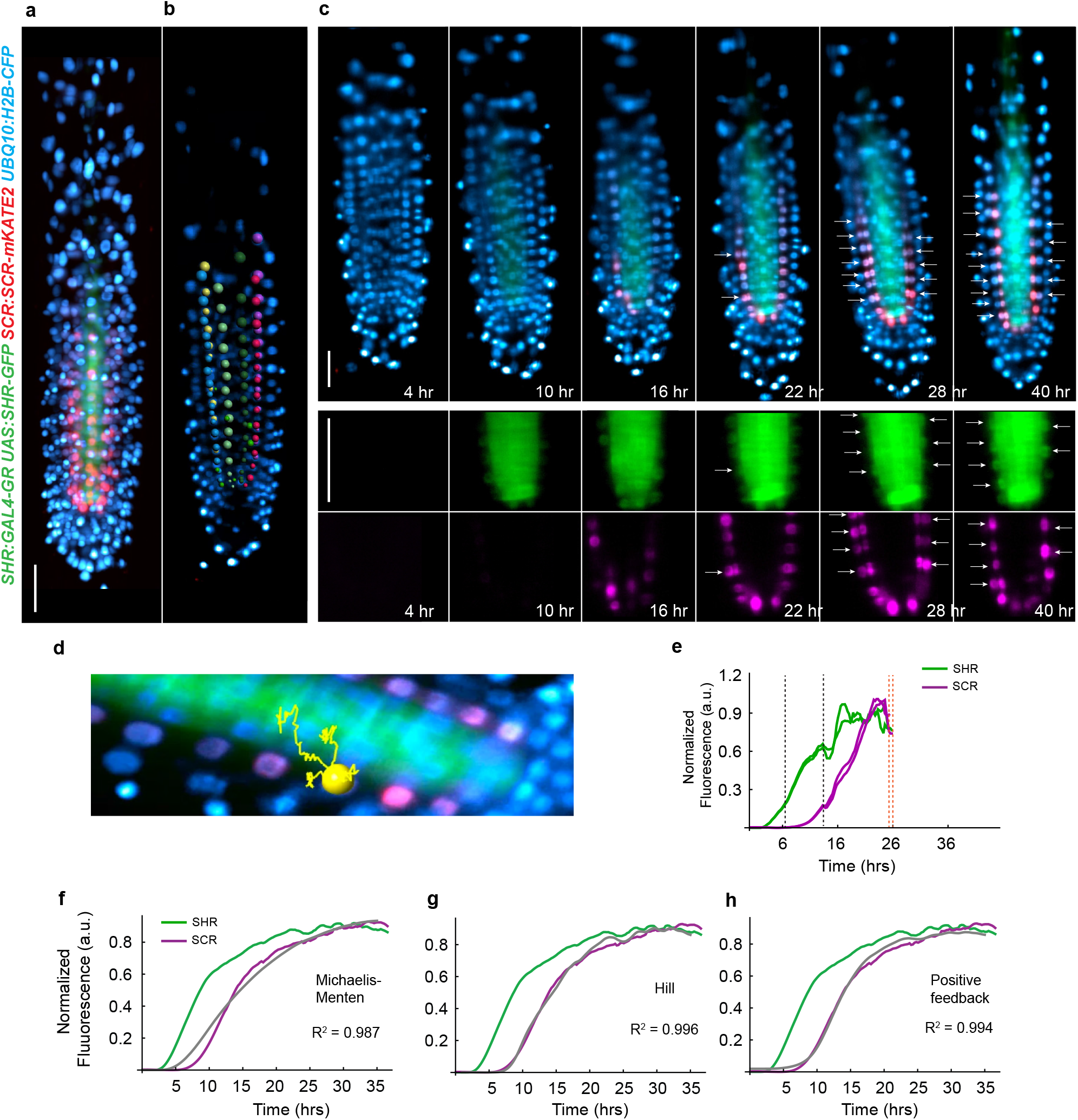
Long-term 4D light sheet imaging of SHR and SCR dynamics in tandem reveals bistability is not needed to model their regulatory relationship. a) 3D reconstruction of z-stack showing induced SHR and SCR expression. Gamma is set to 0.05 to show signal in the mutant layer (scale bar - 50 μm). b) Endodermal nuclei detected by Imaris are selected for quantification. (Colors specify different cell files (scale bar - 50 μm). c) Median longitudinal sections showing timecourse of fully induced (40 μM dex) root at various timepoints after treatment. White arrows, asymmetric divisions; SHR-GFP, SCR-mKATE2, and H2B-CFP signal (top), SHR-GFP and SCR-mKATE2 alone (bottom) (scale bar - 50 μm). d) Nucleus selected for quantification in Imaris visualized on a median section of the root. Yellow line, track detected by Imaris showing movement of the nucleus over time within the registered images. e) Quantification of SHR-GFP and SCR-mKATE2 fluorescence after full induction for the cell shown in (d). f-h) Mean normalized expression of SHR (green) and SCR (magenta)(see Methods) and predictions of SCR expression from the Michaelis-Menten (f), Hill (g), and Positive Feedback (h) models (grey lines). R-squared, adjusted R-squared; n = 274 cells from 9 roots. Black dashed line, symmetric division; Orange dashed line, asymmetric division.

In fully induced roots, SHR and SCR expression appeared to follow simple dynamics (Fig. 2e). To further investigate the regulation of SCR by SHR, we fit the data to three basic ODE models (see Methods). We first reduced the noise in the system by taking the average of the SHR and SCR dynamics for all cells from fully induced roots (see Methods). The resulting curves closely follow a sigmoid pattern both for SHR and SCR, with SCR having a slightly steeper rise (Fig 2f-h).

For each model, we next used the average SHR dynamics as input to predict SCR expression. We fit the model parameters such that the predicted SCR expression trajectory best matched the measured average SCR dynamics (see Methods). All the models have a linear degradation term and differ only in the production term. The simplest model has a Michaelis-Menten production term (Fig. 2f, adjusted R2 = 0.987), which predicts a linear relationship between SHR and SCR up to a saturation point. In the next model, we used a generalized Hill function (Fig. 2g, adjusted R2 = 0.996), finding a best-fit Hill coefficient larger than 1 (see Methods). A high Hill coefficient suggests that the system shows ultrasensitivity, which is an amplified response to a given input [53,54]. Ultrasensitivity could exist for several reasons, including positive feedback [53,54].

In the third model, we explicitly incorporated positive feedback of SCR [55,56] into a Michaelis-Menten production term (Fig. 2h, adjusted R2 = 0.994).The fits of all three models were comparable. However, qualitatively, the Hill model showing ultrasensitivity visually appeared to capture the rise of SCR better than the Michaelis-Menten model. The third model explicitly incorporating positive feedback also appears to capture the rise of SCR, suggesting that SCR autoregulation could explain the ultrasensitivity we observe (Figure 2f-h). Taken together these monostable alternatives to Cruz-Ramirez et al. [3] indicate bistability is not needed to explain the regulatory relationship of SHR and SCR.

### A low threshold of SHR and SCR specifies asymmetric division early in the cell cycle

SCR levels at the time of asymmetric division varied from cell to cell. Even for cells that divided very close in time and that were derived from the same parent cell through a symmetric division, the expression of SCR at the time of division could vary by as much as 2-fold (Fig. 3a-c), suggesting that the levels of SCR at the time of division are not the critical factor in determining when or whether to divide.

**Fig. 3.**
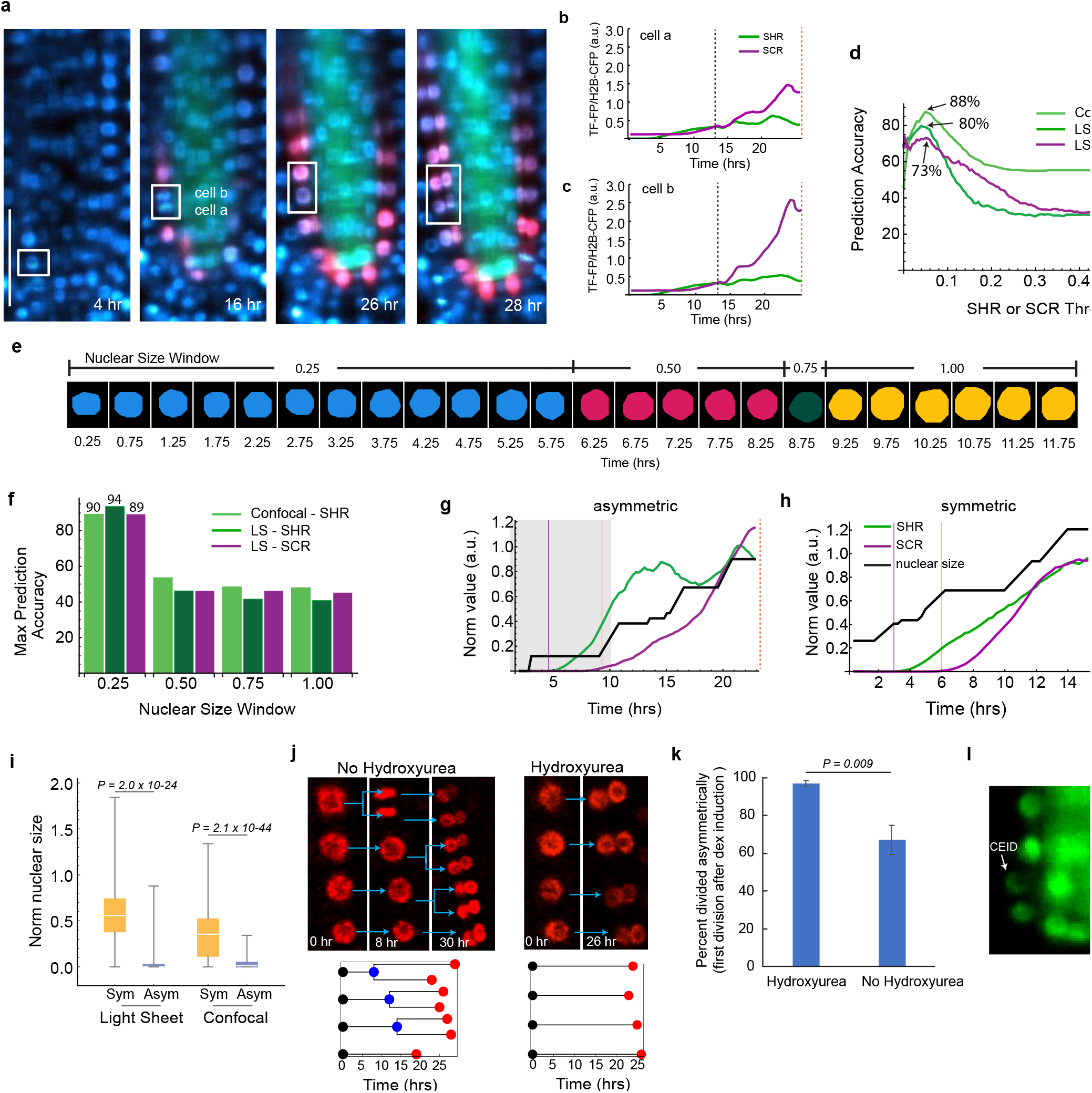
Low threshold levels of SHR and SCR present during an early cell cycle window specify asymmetric division. a) Median light sheet longitudinal sections showing a cell that divided symmetrically into cells a and b (scale bar - 50 μm). b,c) Quantification of transcription factor fluorescent protein (TF-FP) trajectories for the cells in a. SHR-GFP (green) and SCR-mKATE2 (magenta). d) Prediction accuracy of trajectory classification into asymmetrically dividing and non-dividing cells for a range of SHR and SCR thresholds. LS, light sheet; LS, n = 449 cells from 14 roots; Confocal, n = 743 cells from 29 roots; e) Masks used to calculate the nuclear size trajectory for the cell shown in Figure 1d. Every other timepoint is shown. f) Maximum prediction accuracy of trajectory classification into symmetrically and asymmetrically dividing cells across a range of thresholds for a given nuclear size window (see Methods). LS, 500 cells from 14 roots; Confocal, n = 633 cells from 29 roots. g,h) Examples of cells accurately predicted to divide asymmetrically (g) or symmetrically (h). Shaded region corresponds to timeframe within the normalized nuclear size window 0 - 0.25. The normalized nuclear size of the cell in (h) is > 0.25 at the start of the timecourse. Vertical lines indicate times at which SHR (purple) and SCR (orange) thresholds are met. Green, SHR-GFP intensity; Magenta, SCR-mKATE2 intensity; Black, nuclear size. i) Normalized nuclear size at the beginning of the time course for symmetrically and asymmetrically dividing cells from the confocal and light sheet datasets. Two-sided Mann-Whitney test P is shown. LS, symmetric, n = 159 cells from 14 roots; LS asymmetric, n = 59 cells from 11 roots; Confocal, asymmetric, n = 181 cells from 24 roots; Confocal, symmetric, n = 231 cells from 29 roots. j) Median longitudinal confocal images of a cell file after dex induction from roots pre-treated with (right) or without (left) hydroxyurea. Below: Graphical representation showing the timing of symmetric (blue dots) and asymmetric (red dots) divisions for each cell. k) Percent of the first divisions after dex induction that were asymmetric for roots treated with (n = 94 cells from 3 roots) or without hydroxyurea (n = 94 cells from 3 roots). Unpaired one-sided Student’s T-test P is shown. l) Median section light sheet image of SHR-GFP in CEI, CEID, and endodermal cells. Black dashed line, symmetric division; Orange dashed line, asymmetric division. Center line of box plots, median; box limits, 0.25 and 0.75 quartiles; box plot whiskers, full range of data.

We hypothesized that a low threshold level of both SHR and SCR triggers the decision to divide at some timepoint prior to division. To test this hypothesis, we used a discrimination model to determine the accuracy of predicting asymmetric division across a range of SHR and SCR thresholds (Fig. 3d). Optimal thresholds of both SHR and SCR were low relative to the range of SHR and SCR expression levels in the timecourse and were able to accurately predict asymmetric division 80% and 73% of the time, respectively. A similar analysis of the SHR confocal data found a maximum prediction accuracy of 88% (Fig. 3d). The lower prediction accuracy obtained using SCR levels suggests that its role in asymmetric division is secondary to that of SHR.

Despite the high level of accuracy using a SHR threshold, 20% and 12% of the light sheet and confocal trajectories, respectively, were not predicted correctly using a SHR or SCR threshold alone. We considered the possibility that position in the cell cycle or other dynamic features of the SHR and SCR trajectories may contribute to the decision to divide asymmetrically. To test this hypothesis, we extracted from the data the size of the nucleus at each timepoint for all cells and used this as a proxy for position in the cell cycle [12] (see Methods and Figure 3e). We next defined a set of features for the confocal (n = 45) and light sheet (n = 63) data, to describe various aspects of the SHR, SCR and nuclear size trajectory dynamics (e.g., max rate, mean SHR, area under the curve, max and min nuclear size, etc. (Supplementary Data Table 3). We assessed the ability of each feature to discriminate between asymmetrically dividing and non-dividing cells (see Methods).

Among the most predictive features were those associated with levels of SHR (mean SHR, max SHR, quantile 0.5 SHR). Notably, features describing SCR levels were less predictive than SHR levels, in support of a primary role for SHR in driving the decision to divide asymmetrically. Features relating to nuclear size were also significant predictors of asymmetric division such as ‘mean nuclear size’ and ‘norm nuclear size at SHR threshold’ (Extended Data Tables 1 and 2). This suggested the possibility that threshold levels of SHR might be required during a specific window of the cell cycle for asymmetric division to occur. In this hypothesis, if SHR is absent during a critical window, the cell commits to a symmetric division, and subsequent exposure to above-threshold levels of SHR does not alter this program.

To test this hypothesis, we separated each trajectory into separate cells if one or more symmetric divisions occurred. Next, we used a discrimination analysis (see Methods) to determine the accuracy of predicting asymmetric vs. symmetric division for each cell based on whether a threshold of SHR or SCR was reached during one of four quarters of the cell cycle (approximated by nuclear size [12]; Fig. 3e). We performed this analysis across a range of thresholds and found that requiring the threshold for both SHR and SCR to be met in the first quarter of the cell cycle resulted in higher accuracies (90%, 94% and 89% for the confocal SHR, light sheet SHR, and light sheet SCR, respectively) than the predictions using threshold alone (Fig. 3d) and higher accuracy than requiring the threshold to be met in any of the other three quarters of the cell cycle (Fig. 3f-h).

In support of the hypothesis that threshold levels of SHR must be met early during the cell cycle, we found that cells that divided symmetrically had significantly larger nuclei at the beginning of the timecourse compared to asymmetrically dividing cells (Fig. 3i), suggesting that these cells were already past a critical window of the cell cycle when SHR first reached threshold levels.

We next recalculated the dynamic features for SHR and SCR trajectories occurring over the course of a single cell cycle and determined the ability of each of these features to predict asymmetrically and symmetrically dividing cells. For both the light sheet and confocal datasets, the most predictive feature was ‘Maximum SHR level during nuclear size window of 0 – 0.25’, accurately predicting 94% and 89% of the two respective datasets (Extended Data Tables 3 and 4). Finally, using a support vector machine algorithm applied to all the features together, we can accurately predict division 94% of the time for both the light sheet and confocal data.

To further examine whether position in the cell cycle determines sensitivity to SHR, we synchronized the cell cycle throughout the root meristem by treating SHR:GAL4-GR UAS:SHR-GFP *shr2* roots with 2 mM hydroxyurea for 17 hours just prior to induction with dex. This treatment causes cell cycle arrest at the G1/S transition for most cells [57]. We anticipated that treating synchronized roots with dex would result in larger numbers of cells exposed to SHR during the critical early cell cycle window, leading to greater numbers of asymmetrically dividing cells.

Consistent with our hypothesis, 97% +/- 0.02% of the first divisions after dex induction (n = 3 roots) in hydroxyurea-treated roots were asymmetric compared to only 67% +/-0.08% of cells (n = 3 roots) treated with dex alone (Fig. 3j and k, Fig. S8, p-value = 0.009; Extended Data Videos 10 and 11).

To understand how our findings inform division of the CEID in wild-type plants, we investigated SHR expression in SHR:SHR-GFP SCR:SCR-mKATE2 UBQ10:H2B-CFP *shr2* plants, which have a wild-type phenotype. We were able to capture one timecourse that included both a CEI division and subsequent division of the CEID (Extended Data Videos 12 and 13). Levels of SHR in both the CEI and the CEID were lower than in the endodermis. We also found that levels of SHR fluctuated but never appeared entirely absent. SHR expression returned to pre-division levels quickly after division of the CEI (Fig. 3l; Extended Data Videos 12 and 13). Thus, it is likely that in wild-type plants, SHR is always present early in the cell cycle of the CEID, providing the conditions necessary for asymmetric division there.

## Discussion

How developmental regulators control cell division is a central question in developmental biology with potentially broad applications in understanding basic cell cycle control. It was previously suggested that SHR and SCR control the decision to divide asymmetrically through a bistable switch [3]. However, from direct observation of SHR and SCR dynamics, we did not detect the expression patterns that would be expected in a bistable system. Although we cannot exclude the possibility of bistability without a definitive test for hysteresis [29,58], which would be nearly impossible to perform in our system, our data suggest that SHR and SCR are unlikely to regulate asymmetric cell division through a bistable mechanism.

We provide evidence for a model in which low threshold levels of SHR and SCR present at a critical early window of the cell cycle trigger an alternate cell cycle program resulting in mitosis with an altered division plane orientation (Fig. 4a) and a shorter duration (Fig. S6i). Notably, however, SHR induction cannot initiate asymmetric division outside of the meristem, indicating that SHR is not sufficient to trigger cell division itself, and must act upon cells that are in a proliferative phase of development. This finding suggests a non-canonical role for SHR, SCR and CYCD6 in determining the orientation of the division plane but not initiation and commitment to division. Thus, the presence or absence of SHR early in the cell cycle determines whether the cell will divide symmetrically or asymmetrically, but other cyclins, and other developmental cues must be present to initiate cell cycle progression (Fig. 4b). A similar mechanism may determine division plane orientation in other systems [59]. In wild-type roots, SHR levels are likely always above the threshold required for division. Thus, the conditions required for asymmetric division of the CEID are met. In the absence of SHR, the CEID cell divides symmetrically, generating the single file of ground tissue cells that we observe in shr mutants.

**Fig. 4.**
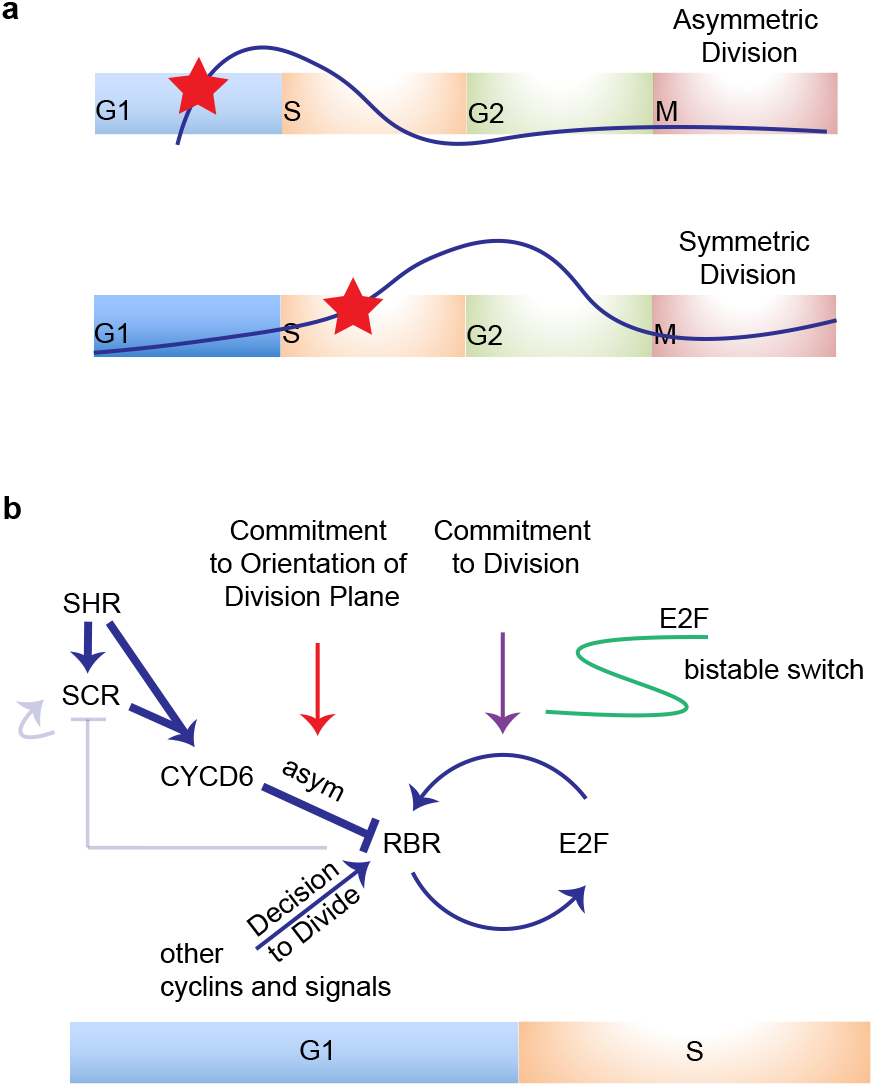
New model for SHR and SCR control of asymmetric division. a) Threshold levels of SHR and SCR (red stars) specify asymmetric division only when present during G1. b) SHR and SCR presence during G1 activates CYCD6 to specify the orientation of the division plane. CYCD6 and other cyclins with their associated kinases phosphorylate RBR, triggering the RBR-E2F bistable switch that commits the cell to asymmetric division. The two positive feedback loops (SCR autoregulatory loop and RBR release of SCR after phosphorylation by CYCD6) play a smaller role during asymmetric division than previously predicted [3] but may be required to generate the high levels of SHR and SCR needed after division.

CYCD6 and other D-type cyclins interface with the cell cycle at the RB-E2F bistable switch that commits the cell to DNA synthesis and irreversible progression through mitosis [29]. Thus, it is most likely that the critical window of sensitivity to SHR occurs prior to the G1-S restriction point (Fig. 4b). This is consistent with previous studies showing the duration of G1 is important for division plane orientation, which suggested that developmental cues specifying asymmetric division are perceived then [59]. The RB-E2F bistable switch acts to integrate the many signals indicating a cell’s readiness to divide [60]. Thus, it is possible that both the timing and orientation of CEID division are determined there. Our findings suggest that rather than triggering asymmetric division through bistable levels themselves, threshold levels of SHR and SCR at the right time alter the division plane and engage an existing cell cycle bistable switch that commits the cell to division.

The dynamic patterns of SHR and SCR expression that we measured are consistent with the presence of positive feedback that could arise from SCR autoregulation (Cui, 2007). Positive feedback can be used to amplify expression [61]. Yet, we observed that the levels of SHR and SCR required for asymmetric division are much lower than the levels at which both proteins are ultimately expressed, and that levels of SHR in the CEID are lower than in the endodermis. These findings are consistent with previous observations that asymmetric division still occurs when SCR levels are significantly reduced [16]. High levels of SHR and SCR are needed in the endodermis to prevent further asymmetric divisions and for SCR to restrict SHR movement into adjacent layers [55]. Thus, it is possible that positive feedback and/or bistability might play a role in generating the high levels of SHR/SCR needed immediately after division to prevent further divisions in the endodermis.

Bistability was proposed to explain both how the decision to divide asymmetrically is made, as well as how it is restricted to the CEID given that SHR and SCR are also expressed elsewhere [3]. The evidence we present here sheds light on how SHR and SCR interface with the cell cycle to specify asymmetric division but suggests an alternate mechanism must exist for spatial and temporal control. The high levels of SHR in the endodermis suppress asymmetric divisions there [62]. Additionally, recent evidence indicates that the Bam/Cle signaling pathway plays a role in restricting asymmetric division to the stem cell niche [63]. It is thus likely that levels of SHR and/or other coregulators are important for restricting asymmetric division in space and time in wild-type plants.

SHR and SCR directly regulate numerous other components of the CYCLIN D/RBR/E2F pathway [18], in addition to CYCD6. This pathway is highly conserved between plants and animals, including humans [64,65], and perturbation of its components is estimated to occur during the development of nearly all cancers. Regulation of CYCLIN D/CDKs is a target of many cancer therapies [66–70]. Furthermore, defects in division plane orientation and asymmetric division have recently been implicated in the genesis of breast and other cancers [7,71]. Most studies of cell cycle control have been in single-cell organisms or cell lines. Future studies of the dynamics of SHR and its cell cycle targets using the system described here could reveal new mechanisms of cell cycle control important during the development of multicellular organisms and suggest opportunities for novel therapeutic interventions in cancer pathogenesis or prevention.

## Materials and Methods

### Construction of Plant Lines

*Arabidopsis thaliana* accession Columbia (Col-0) was used in this study. The SHR:GAL4-GR UAS:SHR-eGFP, SCR:SCR-mKATE2 and UBQ10:H2B-CFP DNA constructs were generated using the Invitrogen Multi-Site Gateway^®^Three-Fragment Vector Construction Kit. GAL4-GR UAS was PCR-amplified from pDONOR 221 GAL4::GR::UAS::GFP::UAS [56] and cloned into the D-TOPO vector (Invitrogen). The SHR CDS was PCR-amplified from a pDONR221 SHR vector, introducing flanking B2 and BamHIXhoI-B3 sequences and cloned into pGEM-T Easy. eGFP [72] was also cloned into pGEM-T Easy, amplified with flanking BamH1 and XhoI sites. A BamH1-XhoI double digest followed by ligation produced pGEM-T Easy SHR-eGFP. SHR-eGFP was inserted into P2R-P3 by linearizing pGEM-T Easy SHR-eGFP with BsaI followed by a BP reaction. D-TOPO GAL4-GR UAS and P2R-P3 SHR-eGFP were combined with pENTR5’pSHR (2.5 kb) [50] into dpGreenBarT [73] in an LR reaction. To make the SCR:SCR-mKATE2 construct, mKate2 was amplified from the pmKATE2-N plasmid from Evrogen introducing flanking B2 and BamHIXhoI-B3 sequences and cloned into pGEM-T Easy. The 19S terminator was amplified, introducing flanking BamHI and XhoI sites and cloned into pGEM-T Easy. mKATE2-19S pGEM-T Easy was generated by a double BamHI XhoI digest of the mKATE2 and 19S pGEM-T Easy vectors. mKATE2-19S was then inserted into P2R-P3 by a BP reaction. This plasmid was combined with pDONORP4-P1R pSCR2.0, and pDONR221 SCR CDS in an LR reaction into pGII0125 [74]. To make UBQ10:H2B-CFP, the H2B CDS from AT5G22880 was amplified and introduced into D-TOPO. This vector, along with pGEM-P2-sCFP-P3 [75] was combined with UBQ10 P4-P1R [76] into dpGreenKanT [77]. The constructs were transformed into Arabidopsis by the floral dip method [78]. To make the triple line, UBQ10:H2B-CFP was transformed into the shr2/+ SHR:GAL4-GR UAS:SHR-eGFP line and the resulting line was crossed to SCR:SCR-mKATE2.

### Live Imaging

#### Confocal microscopy

Live imaging of the 2-color SHR:GAL4-GR UAS:SHR-GFP *shr2* UBQ10:H2B-RFP roots was performed using an inverted Zeiss 510 Meta confocal microscope. We acquired 16-bit images at 512 x 512 resolution with a z-step of 2 μm, 34 slices, and 1x averaging. GFP and RFP were imaged sequentially using the 488 nm and 543 nm excitation lines at 12% and 18% power (15 uW and 7 uW at the sample), respectively, with a Zeiss C-Apochromat 40X 1.2 W Korr water immersion objective (Part #441757-9970). Plants for imaging were grown for 5 days in square Petri dishes containing 1X Murashige and Skoog (MS) 1% sucrose 1% agar media oriented vertically in a Percival in long day growth conditions (16hr/8hr light/dark regime). Prior to imaging, plants were transferred to cell culture/imaging chambers (Thermofisher, Cat. #155360). Roots were covered with a small block of 1% Phytagel containing various concentrations of dex. For hydroxyurea treatments, roots were transferred to MS plates containing 2 μM hydroxyurea (Sigma) for 17 hours prior to transfer to an imaging chamber. The chamber was closed with the provided top to prevent water loss. Immersol was used instead of water as the immersion media to prevent evaporation over the timecourse. We acquired 4 tiled images every 15 minutes for each experiment for up to 24 hours. The tiles were aligned linearly, and we began each experiment with the root tip located at the top of the first tile. The root grew across the four tiles over the course of the 24-hour timecourse.

#### Light sheet microscopy

Light sheet imaging of SHR:GAL4-GR UAS:SHR-eGFP *shr2* SCR:SCR-mKATE2 UBQ10:H2B-CFP and SHR:SHR-GFP *shr2* SCR:SCR-mKATE2 UBQ10:H2B-CFP roots was carried out using a custom-built light sheet microscope (see Light sheet microscope optical setup). Roots were mounted vertically (see Light sheet sample preparation and mounting) and illuminated bidirectionally using scanned light sheets, creating an optical section parallel to the length of the root. The fluorophores eGFP, sCFP3a, and mKATE2 were imaged using 488, 457, and 561 nm excitation lasers at 10, 10, and 30 percent power, respectively. Laser power measured at the output of the optical fiber was 3.0 mW, 3.1 mW, and 7.4 mW +/- 10% for the 488, 457, and 561 nm lasers, respectively, and the optical throughput is approximately 5% from the fiber output to the sample. Multi-color z-stacks of 130 slices were captured at 15-minute intervals for up to 48 hours using a 300-millisecond exposure time and a step size of 1 micron. Images were captured in 16 bits with a pixel size of 0.197 μm. Plants were illuminated from the top with white light (140 μmol/m2/sec), synchronized to the data acquisition to turn off during the camera exposure time.

#### Light sheet microscope optical setup

Continuous-wave 488, 457, and 561 nm visible laser light from an Omicron SOLE 6 Compact Laser Light Engine were combined into a single beam and subsequently split and sent into the left and right sides of the imaging chamber, through two illumination objectives (Nikon Plan Fluorite, 10X, NA = 0.3, water dipping)) to bidirectionally illuminate the sample. Pairs of galvanometers (H6215, Cambridge Technologies) upstream of each of the illumination objectives generated a fast-scanned (500 Hz) sheet of light approximately 2 microns thick at the focal point. Fluorescence emitted from the sample was detected through a water-immersion detection objective (Olympus XLUMPLFLN20XW, 20X, NA = 1.0), filtered through detection filters specific to the appropriate fluorophore (Semrock, GFP: FF03-525/50-25; mKATE2: FF01-624/40-25; CFP: FF01-482/35-25) mounted on a filter wheel (Sutter Instruments Lambda 10-3) and a short-pass 750nm filter, and projected onto a cMOS camera (Andor Zyla 5.5) through a double-achromatic tube lens (focal length = 300 mm; Thorlabs ACT508-300-A-ML), to yield a final magnification of 33X. A piezo translational stage (Physiks Instruments P-622.1CD, with controller E-665) controlled the sample’s fine z-motion for z-stacking, and a rotational stage (Newport PR50CC, with controller ESP300) controlled the angular position. Both stages are mounted onto a 3-dimensional stepper-motor stage stackup (Sutter Instrument MPC-200), which provided coarse positioning. A white LED light source (Thorlabs MCWHF2, driver LEDD1) was used to provide ambient light to the plant sample.

All microscope hardware components (stages, lasers, camera, filter wheel, and white ambient light) were controlled using a custom-written Java application (code available upon request) utilizing the MicroManager core API [79]. Root tips were tracked by shifting the stepper-motor stage positions for the current imaging round by extrapolation using the difference (in x, y, and z) between the centroids of the root tips in the prior two imaging rounds.

#### Light sheet sample preparation and mounting

Seeds were sterilized with chlorine gas, imbibed and stratified for 2 days prior to planting in FEP (Fluorinated Ethylene Propylene) tubes with an inner diameter of 1/32” and outer diameter of 1/16” (Cole-Parmer, Cat #06406-60). Tubes were cut to 30 mm in length, autoclaved, and filled with 1X MS salt mixture, 1% sucrose, 1% Phytagel media. A hollow channel was created by inserting a 150-micron diameter steel rod into the tubing containing molten Phytagel along the edge of the inner wall. Seeds were planted, embryonic root side down, at the opening of the channel. FEP tubes were attached to the inside of square petri dishes sealed with micropore tape that were oriented vertically in a Percival set to 21°C at 5500 lux and programmed for long day conditions. Seeds were grown for 5 days prior to imaging. Immediately prior to imaging, FEP tubes were cut to a length of 15 mm, affixed to a custom-designed, 3D-printed sample holder (Fig. S7b, 3D design files available upon request) and oriented such that the hollow channel containing the root was located closest to the detection objective. The sample holder was lowered into a water-filled chamber, submerging the tip of the FEP tubing, but maintaining the shoot portions of the plant above water. For dex treatments, we used different concentrations of dex, as either a continuous treatment or as pulses. A concentrated dex solution was added directly to the water in the chamber and allowed to diffuse into the bottom of the FEP tubing. For continuous treatments (40 μM or 0.04 μM), the dex solution remained in the chamber for the duration of the timecourse. For pulsed treatments (20 μM and 40 μM dex), we replaced the media after 1 minute and 5 seconds, respectively. We used a 40 μM dex working concentration in the imaging chamber to get maximal induction of asymmetric divisions in the root tip. The higher concentration of dex relative to that used for the confocal experiments is likely due to the need for diffusion through the imaging chamber and capillary tube in the light sheet sample mounting setup.

### Data Analysis

Data were processed and analyzed using Python 3.7, Wolfram Mathematica 13.1, R 3.3.3, Fiji 1.50 and 1.52e, and Imaris 9.5.0.

#### Image pre-processing

Image pre-processing was automated using a series of custom Fiji plugins that utilized native ImageJ functionality. The 16-bit images were first cropped to reduce black space. Next, pixels were binned 2×2 using the ImageJ plugin ‘Binner’ with the ‘average’ method and converted to 8 bits by mapping pixel values in the range min:max to the range 0:255 (8-bit) using the setMinandMax Fiji function. The minimum value (min) was set to the average camera background of 100. The maximum value (max) was set for each channel to minimize saturation in the cells of interest. The maximum values used for the red, green and blue channels were 2500, 2000 and 2300, respectively. Blue channel (nuclei) images were registered over time to each other using the PhaseCorrelation function from the Python ImageLib library, which was embedded into a custom ImageJ plugin. Images from all three channels were shifted according to the computed linear shifts.

#### Cell tracking and gene expression quantification

To obtain the expression trajectories for SHR and SCR, signal intensities for all three channels within the nuclear regions of single cell lineages were obtained through one of two automated methods. For the confocal data, we used a custom segmentation and tracking algorithm written in JAVA as Fiji plugins, utilizing native ImageJ functionality to generate a “track” for each cell consisting of a small, cropped image of the nucleus of only that cell at each timepoint. Each image contained the median z-slice through the nucleus at that timepoint. To do this, we used registered hyperstack images containing all timepoints and z-slices for the nuclei channel as input to the algorithm. A square region of interest (ROI) corresponding to the median section of a user-selected cell at timepoint 1 was added to the ImageJ ROI Manager. Additional ROIs for subsequent timepoints were added to the ROI list automatically by finding the median section of the closest cell at the next timepoint. Segmentation and tracking inaccuracies were corrected manually. Nuclei within a track were segmented using the Otsu method [80]. The segmented nucleus image at each timepoint was then used as a mask to extract the pixel intensities of all three channels.

Signal intensities from the light sheet images were obtaining using Imaris software using the Spot Detection and Tracking algorithms. Spot detection and tracking inaccuracies were corrected manually.

We observed that asymmetric divisions do not occur outside of the meristematic zone. Therefore, we quantified SHR and SCR expression only in cells that remained within the meristematic zone for the duration of the timecourse. This was usually the first five or six cells up from the QC at the start of the timecourse. Only cells from the four or five brightest cell files closest to the detectors (confocal) or camera (light sheet) were quantified.

#### Full trajectories analyses

SHR and SCR gene expression trajectories were created for each cell that span the range from the start of measurement until either the experiment ended without division (max of 24 hours for the confocal data, 48 hours for the light sheet data) or until an asymmetric division occurred. If a symmetric division occurred along the trajectory, then two full trajectories (from start of measurement to end of experiment or asymmetric division) were created, one for each of the daughter cells. The dynamics prior to the symmetric division were duplicated for each daughter cell. The H2B intensity spikes at the time of division which can cause artifacts in the data. Therefore, if a symmetric division occurred, the H2B value for the time that the symmetric division occurred and the previous timepoint were replaced with the average of the previous few timepoints. To remove artifacts, we trimmed up to 4 timepoints from the beginning and ends of the trajectories. Only full trajectories with more than 10 timepoints were retained for analysis.

#### Separated trajectories analyses

To test the hypothesis that SHR levels are evaluated independently for each cell cycle, full trajectories that included one or more symmetric divisions were separated into individual trajectories corresponding to a single cell cycle. Only cells that divided asymmetrically or symmetrically were retained for analysis. Cells that did not divide were not included. To remove artifacts, we trimmed up to 4 timepoints from the beginning and ends of the trajectories.

#### Background Subtraction and Bleedthrough Correction

To correct for autofluorescence and bleedthrough for each channel for the light sheet experiments, we used the following model,

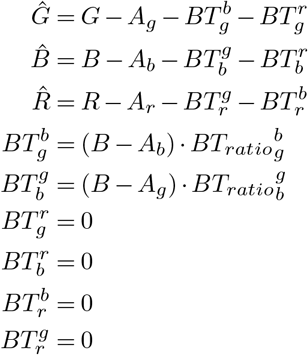

where 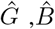 and 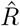 are the corrected signals for the green, blue, and red (corresponding to GFP, CFP, and mKATE2) channels, and G,B,and R are the pixel intensities measured via the Spot Detection algorithm in Imaris for the green, blue, and red (corresponding to GFP, CFP, and mKATE2) channels. *A_x_* is the autofluorescence of the root in the x channel (where x can be g, b, r), 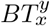 is the bleedthrough of the y channel into the x channel. Bleedthrough involving the red channels were found to be negligible, hence we set the corresponding parameters to 0.

We took advantage of the fact that SHR was not yet induced at the early timepoints and used the average of the first 3 timepoints in the green channel for each cell as the estimate for *A_g_*. The values for *A_r_* were estimated from the image stack at the first timepoint, as the average of the red channel pixel intensities of multiple randomly selected spots within the root tip (but not including the ground tissue). The value for *A_b_* was obtained from roots that did not contain the H2B-CFP marker, as the average of the blue pixel intensities for multiple randomly selected spots within the root tip. The value for 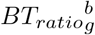 was empirically determined to be 0.017 based on images of roots containing only the UBQ10:H2B-CFP construct and imaged with the same settings we used for all experiments. The value for 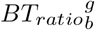 was estimated to be 0.137 based on images of induced SHR:GAL4-GR UAS:SHR-GFP *shr2* plants. For the confocal data, bleedthrough was found to be negligible. Hence, we used the following model to correct for background autofluorescence:

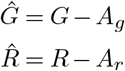

where *A_g_* was estimated as the average of the first 3 timepoints in the green channel for each cell and *A_r_* was estimated from randomly selected regions of roots that did not contain the H2B-RFP marker.

#### Smoothing and Normalization

We smoothed the data for the expression or nuclear size trajectories for each cell using a moving average with a window size of 7. On the edges of the timecourse we used a “reflection” to compensate for missing points in the mean calculation.

Where normalized SHR and SCR trajectories are used, we first divided the SHR-GFP or SCR-mKATE2 fluorescence intensity by the corresponding H2B-RFP or H2B-CFP intensity in that nucleus. Next, for both SHR and SCR trajectories and nuclear size values, we set the minimum value to 0 and the data point at the 90th quantile to 0.9. We used the 90th quantile because it is more robust than the maximal value to noise.

#### Collection of SHR data from the pericycle

We used a custom Python script to extract and quantify the pixel intensities of the pericycle tissue closest to the cell of interest. Using the median z-slice for each cell, we created a localization mask corresponding to two radial sections centered on the center of the cell of interest with a radius of 25 pixels. Each of the two sections was 120 degrees centered on the horizontal axis (see Fig. S5i). We extracted the pixel intensities corresponding to the localization mask. After masking the nucleus using the previously identified nuclear mask, we thresholded the extracted image using the Otsu method (from the OpenCV library (version 4.5.3.56) for Python) and applied the resulting binary mask to the original image to extract the pixel intensities of the pericycle region. Only cell files present along the median longitudinal axis were used in the analysis. To calculate the correlation between pericycle and ground tissue SHR-GFP expression, we used only asymmetrically dividing cells and calculated the correlation up to the timepoint of the first division. SHR-GFP values were normalized to the 90th quantile as described above but were not first divided by the H2B-RFP intensity because those values were not available for the pericycle.

#### Modeling and curve fitting

To reduce the noise for modeling, we derived a single average expression trajectory for SHR and for SCR. We first calculated an average SHR and SCR expression trajectory for each fully induced root for dividing cells. We collected the intensities of the proteins in each cell at each time point and retained only the time points with more than ten measurements to calculate the average for that root. We normalized the dynamics for SHR and SCR for each root as described above. We then manually aligned the different roots to the first inflection point. To obtain one average trajectory for SHR and another for SCR, we took the average expression of all the roots by collecting all the values within a 0.25 hr range.

We tested the fit of our average measured SCR dynamics to the predicted SCR dynamics from three models:

The Michaelis-Menten model:

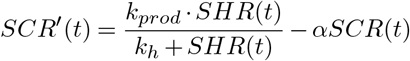

The Hill model:

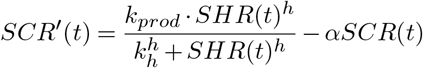

and the Positive Feedback model:

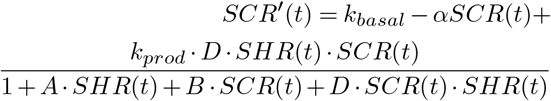

For each model we used the average measured SHR(t) curve as input and identified the parameters corresponding to the best fit between the predicted SCR expression trajectories, SCR(t), and the measured average SCR trajectory. We fit the parameters using a gradient descent method applied to differential equations (https://dpananos.github.io/posts/2019-05-21-odes/).

To determine the model fit, we used the following loss function,

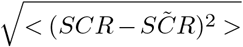

where SCR denotes the measured SCR curve and 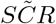 is the estimated SCR curve by the model. When fitting the curves for the Hill model, we used 10 initial condition values for the Hill coefficient, allowing us to reduce the risk of hitting a local minimum in the fitting process and instead reach a global minimum.

We obtained the following best fit parameters:

|  | Model | Parameter | Value |
| --- | --- | --- | --- |
| 1. | Michaelis-Menten | $\alpha$ | 0.144091 |
| | | $k_{Prod}$ | 0.913671 |
| | | $k_h$ | 4.97543 |
| 2. | Hill | $\alpha$ | 1.49001 |
| | | $k_{Prod}$ | 1.7844 |
| | | $k_h$ | 0.746098 |
| | | $h$ | 5.55432 |
| 3. | Positive Feedback | $\alpha$ | 1.4583 |
| | | $k_{Prod}$ | 2.47966 |
| | | $k_{Basal}$ | 0.0282876 |
| | | $A$ | 0.102854 |
| | | $B$ | $1.71131 \cdot 10^{-6}$ |
| | | $D$ | 1.39126 |

To assess the goodness of fit we calculated an adjusted R-squared:

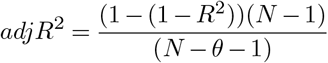

where *R*^2^ is the standard measurement of goodness of fit, N is the number of data points and *θ* is the number of parameters in the model.

#### Calculation of nuclear size

To determine the size of the nucleus for each cell at each timepoint for the confocal data we used a custom Python script to calculate the number of pixels within the circle of the Otsu mask for that nucleus and timepoint. Sometimes, the mask was donut-shaped or was an incomplete circle. To correct for this, we applied a convex hull operation. We first used the OpenCV (version 4.5.3.56) library for Python to calculate the connected components in the mask (two pixels were considered to be touching if they are immediate neighbors horizontally, vertically or diagonally). We filtered the components for those with more than 3 pixels and calculated the convex hull of the new object. This convex hull is considered as the new perimeter of the nucleus. This operation effectively filled in any missing pixels, creating a solid circle. We then calculated the number of white pixels within the circle to determine the size of the nucleus. For the light sheet data, we utilized the ‘Diameter’ statistic for the spot associated with each cell and timepoint.

To calculate normalized nuclear sizes, for trajectories that comprise a complete cell cycle (from symmetric to asymmetric division or symmetric to symmetric division) we set the minimum and the 90th quantile of the nuclear size to 0 and 0.9, respectively. For trajectories that comprise only a part of the cell cycle (from the beginning of the timecourse to the first division), we set the minimum and 90th quantile of the nuclear sizes for the complete family of trajectories (parent and daughter cells) to 0 and 0.9, respectively.

#### Discrimination analysis to identify optimal SHR and SCR thresholds and nuclear size window

To determine the optimal SHR and SCR thresholds for each dataset, we generated a range of thresholds spanning the range of expression values present in the data, from 0 to 0.2 and 0 to 0.6 for the confocal and light sheet data, respectively. We then stepped through each threshold and determined its ability to accurately predict whether a cell divided asymmetrically or did not divide (full trajectories analysis, Fig. 3d) or divided asymmetrically or symmetrically (separated trajectories analysis, Fig 3f). A cell was predicted to divide if the smoothed expression value from any timepoint was equal to or greater than the threshold. For the nuclear size window analysis, we determined the first timepoint where SHR/SCR expression crosses the threshold. If that timepoint fell within the time range corresponding to the nuclear size window (0-0.25, 0.25-0.5, 0.5-0.75 or 0.75-1) it was predicted to divide.

#### Feature analysis

As a preparation step for feature extraction, we aligned the data by their inflection point, defined as a quarter of the way between the 0.1 and 0.9 quantiles. We removed trajectories of non-dividing cells that were shorter than 14 and 38 hours long for the confocal and light sheet data, respectively. We then cropped the remaining trajectories to the aforementioned dimensions. If a cell did not divide by the end of the cropped trajectory, it was labeled as non-dividing even if it divided asymmetrically afterward.

We split the data randomly into train (60%) and test (40%) sets and used the training set to find the threshold and the test set to calculate the accuracy. These sets were used throughout. We defined a set of features to describe various aspects of the aligned trajectory dynamics (see Extended Data Tables 1-4) and found the feature value that maximizes prediction accuracy. For each feature, to determine the value that best separates the dividing and non-dividing cells (full trajectories analysis) or the asymmetrically and symmetrically dividing cells (separated trajectories analysis), we first calculated the mean and standard deviation for each one of the two groups. Next, we defined a range of 3000 steps from the smaller of the two means minus twice its standard deviation, to the larger mean plus twice its standard deviation. For each step in the range, we determined how well the value separated the data and returned the best prediction accuracy out of all values tested. We performed two-tailed Mann-Whitney tests for each feature as a significance measure for the separation of the two groups, and used the Benjamini-Hochberg [81] method for FDR correction.

#### Support vector machine analysis

We used the machine learning algorithm, support vector machine (SVM), to predict the accuracy of separating the data using all the feature values combined. The SVM scores were trained on the training set using the “Classify” function with “SupportVectorMachine” as the method in Wolfram Mathematica. For accuracy prediction we used the ClassifierMeasurement function on the test set.

## Supporting information

Code for trajectory analysis

## Data availability

All imaging data are freely available upon request. All reagents and plant materials are available upon request. Complete trajectories and all metadata tables needed to run the code are included in the Supplementary material.

## Code availability

The trajectory data analysis pipeline code is provided as a supplementary zip file. The microscope controller code and the image processing and quantification pipeline code is available upon request.

## ACKNOWLEDGEMENTS

We would like to thank Stefano DiTalia, Lingchong You, Josh Socolar, Rachel Shahan, Ross Sozzani, Edith Pierre-Jerome, Isaiah Taylor, Trevor Nolan, and Mingyuan Zhu, Sarah Van Dierdonck, and Qianzi Zhou for critical reading of the manuscript and helpful discussions. We thank Dan Holland, Francesco Cutrale, and John Choi at USC for contributing to the light sheet microscope design and construction, and Lisa Cameron and the Light Sheet Microscopy Core Facility at Duke for providing the workstations and support for Imaris image analysis. This work was funded by the US National Institutes of Health (NRSA 5F32GM106690-02 and MIRA 1R35GM131725) to C.W. and P.B., and by the Howard Hughes Medical Institute to P.B. as an Investigator. M.J., S.F., and T.T. were supported by Translational Imaging Center, Bridge Institute, University of Southern California.

## AUTHOR CONTRIBUTIONS

C.W. and P.B. conceived the project and designed the experiments; C.W. and H.B. conducted experiments; C.W. generated transgenic plants; T.T. developed the light sheet imaging platform, with contributions from M.J. and S.F.; C.W., H.B., and R.C. performed image analysis; P.S. and C.W. developed computational tools and performed data analysis of the trajectories, and interpreted the results; C.W. and P.S. wrote the paper with comments from all authors.

---

**Fig. S5.**
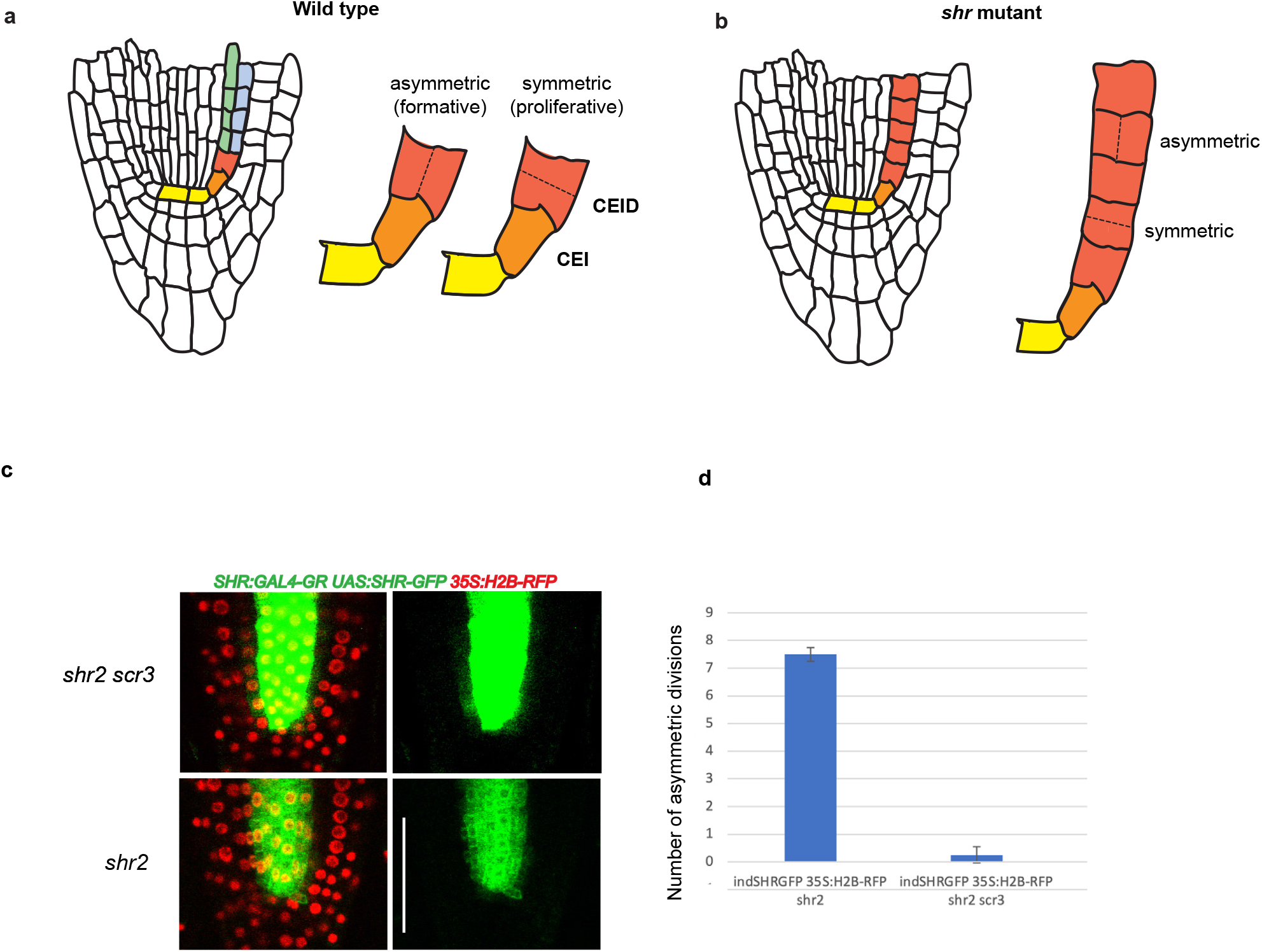
a,b) Diagram of Arabidopsis wild type (a) and *shr* mutant (b) roots showing symmetric and asymmetric division planes. Yellow, QC; Orange, CEI; Red, CEID and shr mutant layer; Blue, cortex; Green, endodermis. c) Confocal images of SHR:GAL4-GR UAS:SHR-GFP 35S:H2B-RFP in a *shr2 scr3* (top) or *shr2* (bottom) background.Roots were fully induced (10 μM dex) for 18 hours. Images show the red and green channels together (left) or the green channel alone (right) (scale bar - 50 μm). d) Number of asymmetric divisions present in the first five cells of 2 cell files in 6-day old inducible SHR-GFP roots in a *shr2* (n = 4 roots from *a* single trial) or *shr2 scr3* (n = 4 roots from a single trial) background after 18 hours of dex. Error bars, s.e.m.

**Fig. S6.**
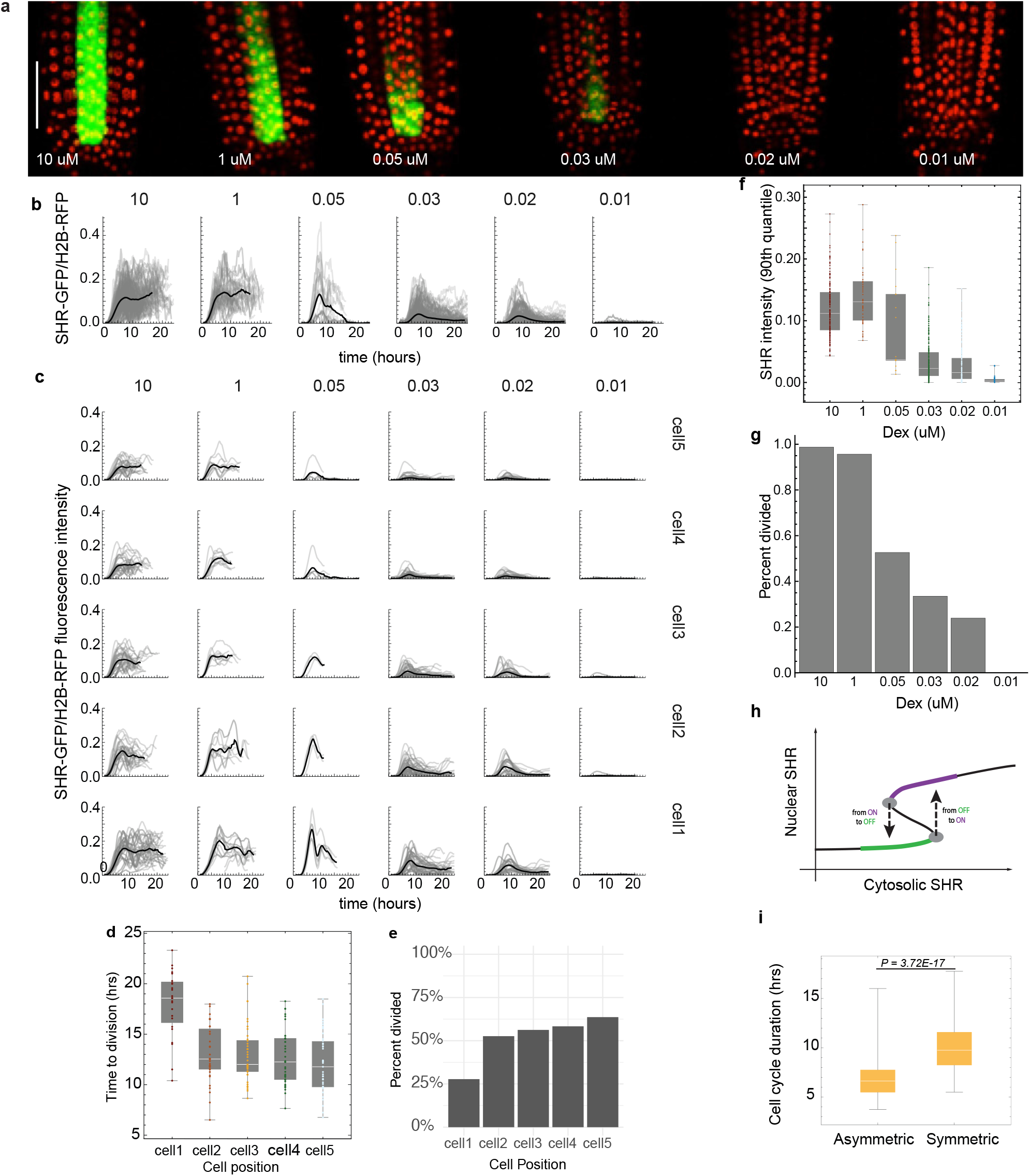
a) Confocal median longitudinal sections of SHR:GAL4-GR UAS:SHR-GFP 35S:H2B-RFP *shr2* roots 18hr after induction with 10, 1,0.05, 0.03, 0.02, and 0.01 μM dex (scale bar - 50 μm). b) SHR trajectories for all cells show a quantitative response to different dex concentrations. Grey line, raw data. Black line, smoothed average. 10 μM, n = 211 cells from 8 roots; 1 μM, n= 63 cells from 2 roots; 0.05 μM, n = 25 cells from 1 root; 0.03 μM; n = 291 cells from 8 roots; 0.02 μM, n = 221 cells from 7 roots; 0.01μM, 124 cells from 3 roots. c) SHR trajectories for all cells broken out by dex concentration and cell position. Grey line, raw data. Black line, smoothed average. For (b) and (c), only timepoints that included at least 30% of the total trajectories for a given dex concentration (b) or dex concentration and cell position (c) were included in the smoothed averages. d) Boxplots of time to division at each cell position for all cells from fully induced roots. cell 1, n = 53 cells; cell 2, n = 38 cells; cell 3, n = 41 cells; cell 4, n = 40 cells; cell 5, n = 39 cells. Cells are from 8 roots. e) Mean percent of cells divided at each cell position for fully induced roots. cell 1, n = 53 cells; cell 2, n = 38 cells; cell 3, n = 41 cells; cell 4, n = 40 cells; cell 5, n = 39 cells. Cells are from 8 roots. f) Boxplots of maximum SHR intensity (90th quantile) for all cells treated with different concentrations of dex. g) Percent of cells that divided asymmetrically by dex concentration. For (f) and (g): 10 μM, n = 158 cells from 8 roots; 1 μM, n = 46 cells from 2 roots.; 0.05 μM, n = 19 cells from 1 root; 0.03 μM, n = 236 cells from 8 roots; 0.02 μM n = 180 cells in 7 roots; 0.01 μM, n = 104 cells from 3 roots. h) Expected hysteresis of nuclear SHR based on the bistable model 3. i) SHR shortens the length of the cell cycle. Average length of the cell cycle for asymmetrically (n = 151 cells from 22 roots) and symmetrically (n = 70 cells from 12 roots) dividing cells. Only complete cell cycles were included (from symmetric to symmetric division, or symmetric to asymmetric division). Unpaired two-sided Mann-Whitney test P is shown. Center line of box plots, median; box limits, 0.25 and 0.75 quartiles; box plot whiskers, full range of data.

**Fig. S7.**
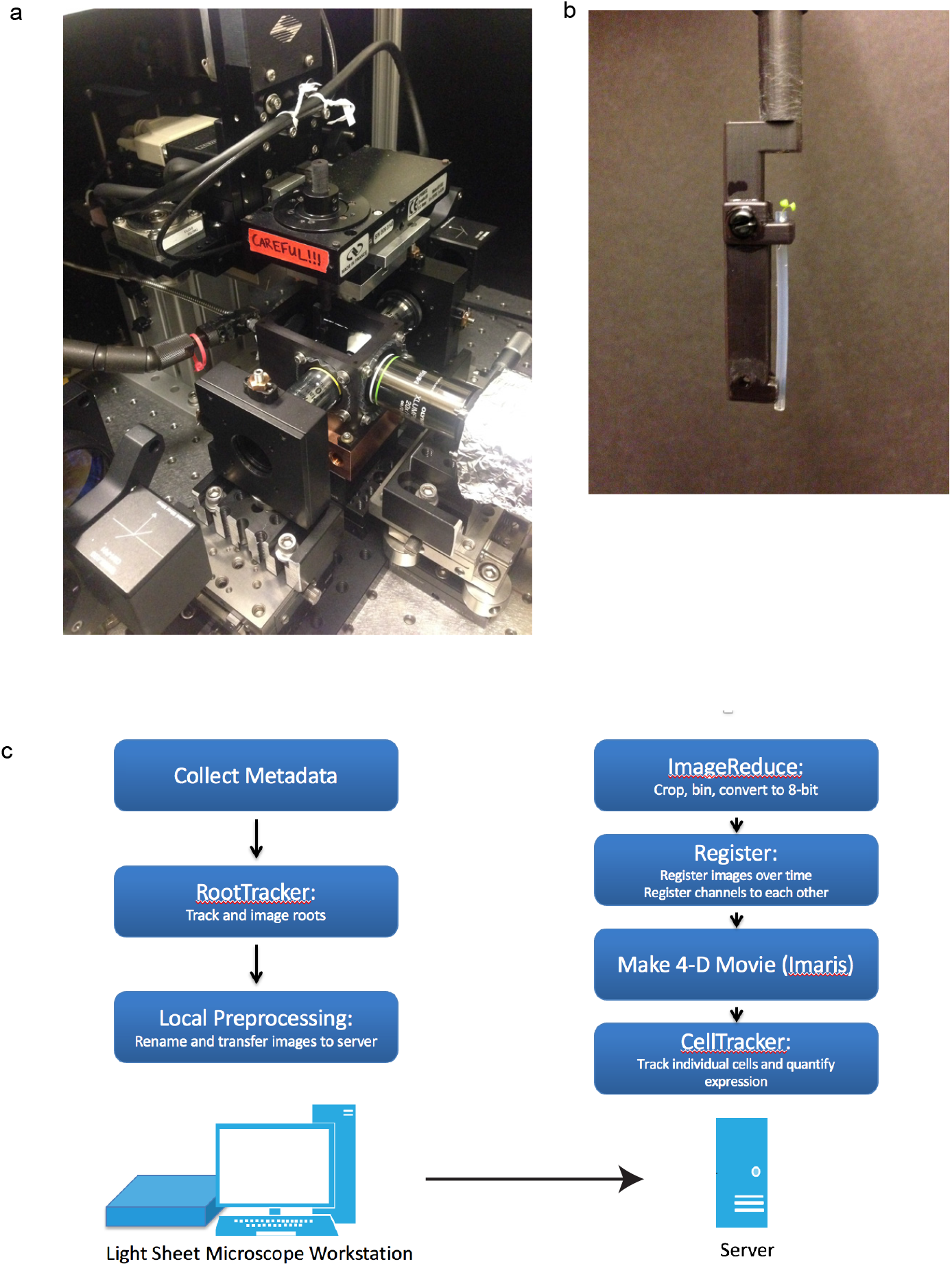
a) Imaging chamber of custom light sheet microscope. b) Capillary tube containing growing root mounted onto custom holder. The holder is lowered into the imaging chamber for imaging. c) Image acquisition and analysis pipeline workflow.

**Fig. S8.**
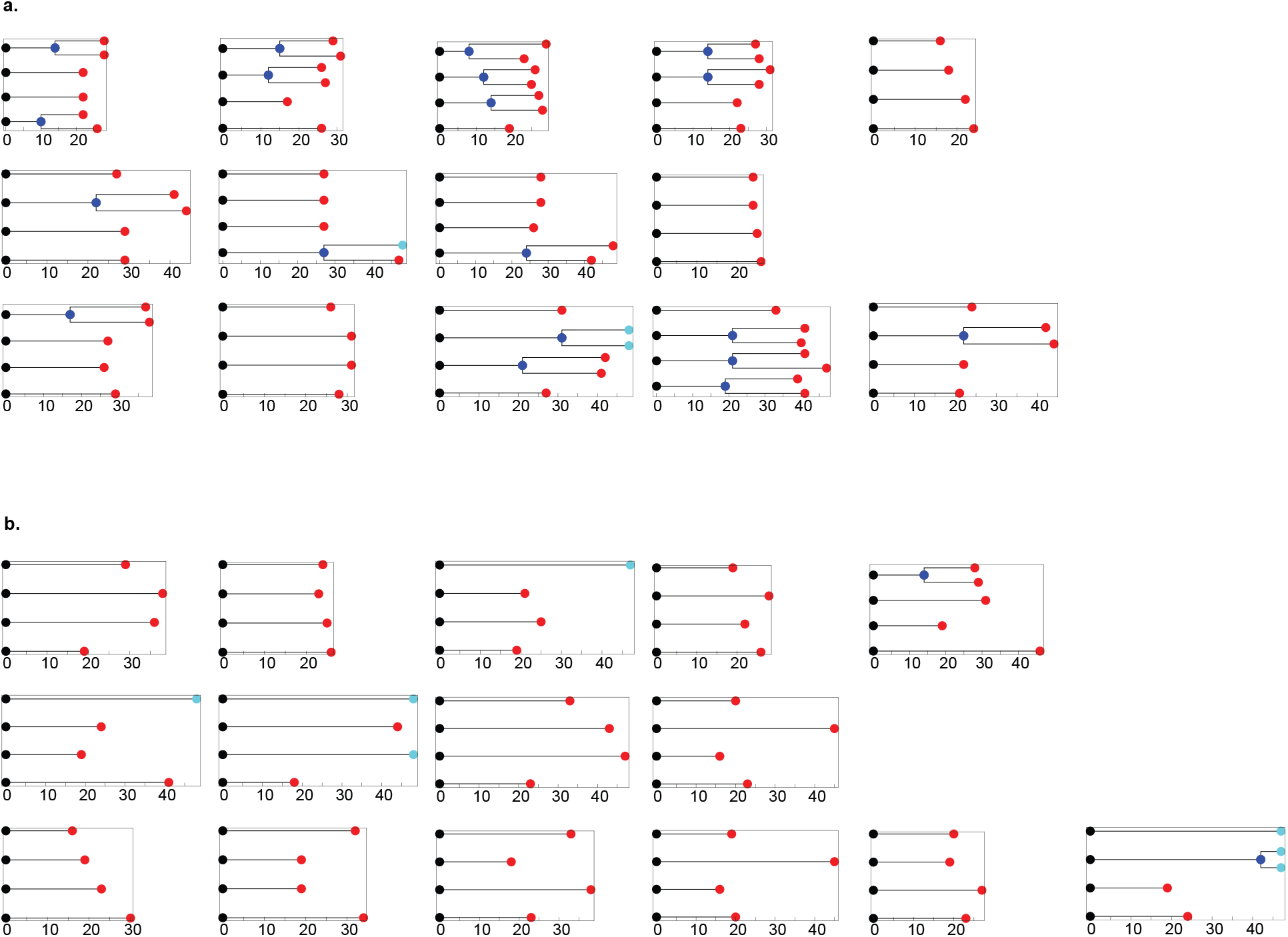
Graphical representation showing the timing of symmetric (blue dots) and asymmetric (red dots) divisions for each cell for roots pre-treated for 17 hours by transfer to plates containing a) mock or b) hydroxyurea followed by transfer to dex for imaging. Each row corresponds to a single root. Each box contains dot plots for cells 2 (bottom) to 5 (top) from a single cell file.

