## Supplementary material for "Patterning and growth are coordinated early in the cell cycle": Code for trajectory analysis: readme.txt

****************************************************************
*Patterning and growth are coordinated early in the cell cycle *
****************************************************************

In this file you will find the information needed to run the code for the trajectory data analysis.

The purpose of this code is to obtain the trajectory data, preprocess it, perform the feature analysis, analyze the thresholds of SHR and SCR for making the decision to asymmetrically divide, fit the different model parameters of SCR given SHR, and export the figures and numbers associated with this paper.    

The code is in the file Script.wls. It is written for Wolfram Mathematica. Specifically it runs successfully in version 13.1.0.0 but is not necessarily limited to this version.

In order to run the code:

a) Install Wolfram Mathematica using:
	https://www.wolfram.com/mathematica/pricing/
	or trial:
	https://www.wolfram.com/mathematica/trial/ (not tested)

    Please note: if you are a part of an institution, there's a chance you can get it through it - please try that option.

b) Open the file using Wolfram Mathematica.
c) The file is divided into cells. In the first cell, you will find two paths: one to the input data folder, and one to the output results folder.
   Please change the paths to correspond to your local directory structure.

To run a cell use shift+return.
To run the entire script, click run all code button on top right.
