## Supplementary material for "Patterning and growth are coordinated early in the cell cycle": Code for trajectory analysis: Script.wls

```
#!/usr/bin/env wolframscript
(* ::Package:: *)

(* ::Section:: *)
(*Definitions and Functions*)


(* ::Subsection:: *)
(*Definitions*)


(* ::Text:: *)
(*Here we define some paths to the data files and results:*)


(*the main folder of the data*)
basePath = "/PATH/TO/DATA/FOLDER/";

(*the location for all the outputs to be stored *)
outputfolder = "/PATH/TO/FINAL/RESULTS/";


(* ::Text:: *)
(*data filenames:*)


(*main confocal data*)
confFileName = "twocolordata_20220909.txt";

(*Figure 3b&c experiment confocal data *)
exp160FileName = "Exp00160_subdata_sort_20220517.txt";

(*hydroxyurea experiment data*)
hydroFileName = "20220221 Divisions in hydroxy vs no hydroxy.xlsx";

(*data for cell size after cell-size correction script - done on masks in a separate python script *)
nsPath = "cellsize_dat_20220310.csv";

(*data for pericycle SHR script  - done on images in a separate python script *)
perPath = "perSHRdat_20220909.csv";

(*main Lightsheet data*)
LSFileName = "2022.07.21_10-34-01_all_data.csv";

(*Experiment metadata Lightsheet data*)
cFileName = "ExperimentMetaData_LightSheet.csv";

(*Experiment metadata Confocal data*)
confmetaFileName = "experiment_info_genotype_points.csv";


(* ::Text:: *)
(*Some Parameter definitions:*)


(*auto-fluorescence estimates for LS and Conf - performed separately manually - see methods*)
AutoFluorescenceEstimatesLS= {4.8,2.1,2.5};
AutoFluorescenceEstimatesConfocal = {1.1, 1.85};


(*bleed-thru matrix - performed separately manually - see methods*)
BTEstimatesConfocal = {{0, 0}, {0, 0}};
BTEstimatesLS = {{0,.017,0},{0.1370,0,0},{0,0,0}};

(*Ranges for threshold calculations*)
rangeLS = Join[Table[th,{th,0,0.01,0.0005}],Table[th,{th,0.01,0.6,0.005}]];
rangeConf = Table[th,{th,0,0.3,0.0025}];

(*Attach current date to folder's name*)
curDate = DateString[{"YearShort", "Month", "Day"}];

(*output location*)
finalFiguresFolder = outputfolder <> "/" <>
     curDate <> "/";

(*sub folders of results for confocal and lightsheet full trajectories and separated trajectories analyses*)
subfoldernames  = {"Conf","ConfSep", "LS","LSSep"};

(*end point of trajectories*)
lptoberemoved =-1;

(*threshold ranges to test for confocal and LS*)
ConfThr = {0.02,0.04,0.06,0.08,0.1,0.2};
LSThr = {0.02,0.05,0.075,0.1,0.3,0.5,0.7,1};


(* ::Text:: *)
(*Some styling definitions:*)


gg= Darker@Green;
mm= Darker@Magenta;
defaultImageSize=Automatic;
defaultColImageSize=Automatic;
colFrameStyle = Automatic;
framestyle = Automatic;
collabel = Automatic;
label = Automatic;
thicknessCol =Automatic;
thicnkess = Automatic;
ticks = Automatic;
ticks = Automatic;
ticks2 = Automatic;
style1d={PlotRange->All,PlotStyle->{ {Opacity[.7],gg}},Ticks-> ticks,LabelStyle->collabel,AxesStyle->colFrameStyle,ImageSize->defaultColImageSize}&/@{1,2,3,4,5};
style1d={PlotRange->All,PlotStyle->{ {Opacity[.7],gg}},Ticks-> ticks,LabelStyle->collabel,AxesStyle->colFrameStyle,ImageSize->defaultColImageSize}&/@{1,2,3,4,5};
style1dLS={PlotRange->All,PlotStyle->{ {Opacity[.7],gg}},Ticks-> (*{2,3}*)ticks,LabelStyle->collabel,AxesStyle->colFrameStyle,ImageSize->defaultColImageSize}&/@{1,2,3,4,5};
style1db={PlotRange->All,PlotStyle-> {{Opacity[.5],Gray}},Ticks-> ticks,LabelStyle->collabel,AxesStyle->colFrameStyle,ImageSize->defaultColImageSize}&/@{1,2,3,4,5};
style1dbLS={PlotRange->All,PlotStyle-> {{Opacity[.5],mm}},Ticks-> (*{2,3}*)ticks,LabelStyle->collabel,AxesStyle->colFrameStyle,ImageSize->defaultColImageSize}&/@{1,2,3,4,5};
style1per={PlotRange->All,LabelStyle->collabel,AxesStyle->colFrameStyle,ImageSize->defaultColImageSize};
style1c={PlotRange->All,PlotStyle-> {{Opacity[.2],Gray}},Ticks-> ticks,LabelStyle->collabel,AxesStyle->colFrameStyle,ImageSize->defaultColImageSize};
style1ca={PlotRange->All,PlotStyle-> {{Opacity[1],Black}},LabelStyle->collabel,FrameTicks-> ticks,Frame->True,FrameStyle->colFrameStyle,FrameLabel-> {"time (h)", "SHR"},AspectRatio->1,ImageSize->defaultImageSize};
style1j={PlotStyle->{{Opacity[0.15],gg}},LabelStyle->label,Ticks-> ticks,AxesStyle->framestyle,FrameLabel->{"Time (h)","SHR Expression"},ImageSize->defaultImageSize, PlotRange-> {All,{0,0.2}}}&/@{1,2};
style3d={PlotStyle->{{Opacity[1]}},LabelStyle->label,Ticks-> ticks,AxesStyle->framestyle,FrameLabel->{"Time (h)","SHR Expression"},ImageSize->defaultImageSize};
style1jO={PlotStyle->{{Opacity[0.5],Green}},LabelStyle->label,Ticks-> ticks2,AxesStyle->framestyle,ImageSize->defaultImageSize, PlotRange-> {All,{0,0.2}}}&/@{1,1};
style1g={PlotRange->All,PlotStyle-> {Opacity[.9],#},LabelStyle->label,AxesStyle->framestyle,Ticks-> ticks,AspectRatio->1,ImageSize->defaultImageSize}&/@{gg,Green};
style2= {FrameLabel-> {"delay (h)","counts"},LabelStyle->label,AxesStyle->framestyle,ChartStyle-> ColorData[1][4],PlotRange->All,ImageSize->defaultImageSize};
cd = ColorData[10];


(* ::Subsection:: *)
(*Functions*)


(* ::Text:: *)
(*Functions that are required during data loading and initial processing *)


(*gets a list of cell names and returns only the cell names that terminate the trajectories. 
Example: Input:{"cell1","cell1a","cell1b","cell1ba","cell1bb","cell2","cell2a","cell2aa","cell2ab","cell2b","cell3","cell3a","cell3b","cell4","cell5","cell5a","cell5b"}
Output: {"cell1a","cell1ba","cell1bb","cell2aa","cell2ab","cell2b","cell3a","cell3b","cell4","cell5a","cell5b"}
*)
isnotContained[l_]:=Block[{list=l},
						 Cases[Table[
							   {item,Or@@(StringContainsQ[#,item]&/@DeleteCases[list,item])}
						  ,{item,list}],{_,False}] [[;;,1]]];


(*smoothes the data using moving average: 
	inputs: dat -> the data
			smoothness -> window size 
	outputs: smoothed data*)
smooth[data_,smoothness_]:=Block[{hlk2,factor,dat},
MovingMap[Mean,data[[;;lptoberemoved]],{smoothness,Center},"Reflected"]];

(*A step in the preparation of duplicated trajectories for each daughter cell that are to share the same mother's trajectory
  Input: gathered data by trajectory
  Output: data with duplicated entries of mother cell but with daughter's name in cell name *)
compTraj[gd_]:= Block[{r,fgd=gd},
r=(Select[#,Length@#==2&]&@Table[Select[#,StringLength@#>=StringLength@dd-1&&StringContainsQ[dd,#]& ],{dd,#}]&@(GatherBy[#,#[[4]]&][[;;,1,4]]))&@gd;
Table[fgd = Join[#,fgd]&@(Flatten[{#[[;;3]],#[[4]]/.Rule@@pair,#[[5;;]]}]&/@Select[fgd,#[[4]]==pair[[1]]&]),{pair,r}];
DeleteDuplicates@fgd
];

(*Function to normalize the data so that the min of the data will be at 0 and the 0.9 quantile will be at .9
input: a vector
output: a normalized vector*)
normalize[v_] :=Block[{v9=Quantile[v,.9],v1=Min[v]}, If[v9!= v1 , 0.9(v-v1)/(v9-v1),0.9(v-v1)/(v9+0.01)]];


(*change the data format and return both normalized and unnormalized versions.
inputs: dat-> the data
        rule -> a lookup table linking cell identifiers to min and max values for normalization
        trajNorm -> a flag to whether normalize the data as joined (True) or separated (False)
outputs: a normalized version of the data in the wn** format
        *)
d2wn[dat_,rule_,trajNorm_] := DeleteCases[Join[#[[;;5]],{SortBy[N@#[[6]],First]},
								{SortBy[Join[{#[[1,1]]},Block[{q=#[[1,2;;-2]]}, normalize[#]&/@q],
								{0.9 (#[[1,-1]]-#[[2,1]])/(#[[2,2]]-#[[2,1]])}]\[Transpose]&@{#[[6]]\[Transpose],{#[[1]],#[[2]],If[trajNorm,StringTake[#[[3]],5],#[[3]]]}/.rule},First]}
								,#[[8;;]]]&/@dat,{__,"N",__}]/.{"A"->1,"S"->0};


(* ::Text:: *)
(*a group of functions that align the data*)


(*align the data to the time it reaches ~25% of the expression range and return cropped trajectory
	inputs: dat -> the data in wn format
			q -> total length in hours of the cropped trajectories
			i -> the time a trajectory must have present before the alignment point; set -i to 0 
	output: aligned data in the same wn format*)
aligningByRise[dat_,q_,i_]:= Block[{midPoints,mdivsC,aligned,sel},
						midPoints=# [[FirstPosition[If[#>0,1,0]&/@(((0.25(Quantile[#,0.9]-Quantile[#,0.1])+Quantile[#,0.1])&@#[[;;,2]]) - #[[;;,2]]),0],1]]&/@dat[[;;,6]];
						mdivsC =  Join[{#[[1,;;,1]]-#[[2]]},#[[1,;;,2;;]]\[Transpose]]\[Transpose]&/@({dat[[;;,6]],midPoints[[;;,1]]}\[Transpose]);
						sel = Select[({dat,mdivsC}\[Transpose]),(Min@#[[2,;;,1]]<= -i && Max@#[[2,;;,1]]>= q-i&& #[[1,8]]==0)||#[[1,8]]==1&];
						aligned =Join[#[[1,;;5]],{Join[{#[[;;,1]]+i},#[[;;,2;;]]\[Transpose]]\[Transpose],Join[{#[[;;,1]]+i},Block[{q2=#[[;;,2;;]]\[Transpose]}, normalize[#]&/@q2]]\[Transpose]}&@Select[
						SortBy[#[[2]],#[[1]]&],#[[1]]>= -i&&#[[1]]<=q-i&],If[#[[2,-1,1]]>=q-i,{0},{1}],#[[1,8;;]]]&/@sel;
						GatherBy[aligned,#[[6]]&][[;;,1]]];

(*shift every trajectory to start at t=0 
	inputs: dat -> the data in wn format
	output: aligned data in the same wn format*)
aligningTostart[dat_]:= Block[{aligned},
						aligned= Join[#[[;;5]],{N@Block[{m=Min@#[[;;,1]]},Join[{#[[;;,1]]-m},#[[;;,2;;]]\[Transpose]]\[Transpose]]&@#[[6]]},{N@Block[{m=Min@#[[;;,1]]},Join[{#[[;;,1]]-m},#[[;;,2;;]]\[Transpose]]\[Transpose]]&@#[[7]]},
						#[[8;;]]]&/@dat;
						GatherBy[aligned,#[[6]]&][[;;,1]]];

(*shift every trajectory to end at t=0 
	inputs: dat -> the data in wn format
	output: aligned data in the same wn format*)
aligningToEnd[dat_]:= Block[{aligned},
						aligned= Join[#[[;;5]],{N@Block[{m=Max@#[[;;,1]]},Join[{#[[;;,1]]-m},#[[;;,2;;]]\[Transpose]]\[Transpose]]&@#[[6]]},{N@Block[{m=Max@#[[;;,1]]},Join[{#[[;;,1]]-m},#[[;;,2;;]]\[Transpose]]\[Transpose]]&@#[[7]]},
						#[[8;;]]]&/@dat;
						GatherBy[aligned,#[[6]]&][[;;,1]]];


(* ::Text:: *)
(*Functions related to feature extraction*)


(*Take a data vector (expression values and nuclear sizes) as input and output the mean, sd and cv for the vector*)
stat[vec_]:={Mean@#,StandardDeviation@#,StandardDeviation@#/Mean@#}&@vec;

(*find the time at max rate and return the corresponding data at that time
inputs: vec -> data in {time,val1,val2,...,valN} format}
		rateDet -> the column number to use for determining the position of max rate
		strech -> the step size (dt in dx/dt), a small step size is more accurate but very sensitive to noise  
outputs: the data point at which max rate is achieved*)
rates[vec_,rateDet_, stretch_]:= {vec[[{Position[#,Max[#]][[1,1]]+(stretch-1)/2}]],{Max@#}}&@
								((vec[[stretch;;,rateDet]]-vec[[;;-stretch,rateDet]])/(vec[[stretch;;,1]]-vec[[;;-stretch,1]]));

(*return the Area Under the Curve of a vector in {time, val} format*)
auc[vec_]:= NIntegrate[Interpolation[vec][x],{x,vec[[1,1]],vec[[-1,1]]}];

(*calculate the value of a vector (expression values and nuclear sizes) for a given set of quantiles
inputs: vec -> data in {time,val1,val2,...,valN} format}
		qdet -> the column number to use for determining the quantiles
		vals -> a set of quantiles
		
outputs: the data points at which the relevant data reaches the quantiles*)
quantiles[vec_,qdet_,vals_]:= Table[{ Quantile[vec[[;;,qdet]],q], SortBy[Select[vec,#[[qdet]]>= Quantile[vec[[;;,qdet]],q]&],First][[1,1]]},{q,vals}];

(*returns the min and max of a vector*)
noNorms[vec_]:= {Min@#,Max@#}&@vec;

(* the feature extraction function
intputs: dat -> the aligned dataset in wn format 
		 sep -> a flag that specifies if the features for the seperated datasets should be calculated
		 scr -> a flag indicating whether scr values exist in the data 
		 thresh -> a list of threshold values for shr  
		 threshSCR -> a list of threshold values for scr  
		 shrThs -> a list of threshold values for nuclear size  
outputs:
		{feature-titles, features}
		 
		 
 *)
calcFeatures[dat_,sep_,scr_,thresh_,threshSCR_,shrThs_]:= Block[{stretch=7,initTitles,initFeatures,rftitlesC,ratesFeaturesConfocal,iftitlesC,IntFeaturesConfocal,cellsizetMAXitles,cellsizetMAXitlesSCR,cellsizetCumuMAXitles,cellsizetCumuMAXitlesSCR,
									  sftitlesC,statFeaturesConfocal,tlftitlesC,timelevelsFeaturesConfocal,csftitlesC,cellsizelevelsFeaturesConfocal,cellsizetAUCitlesSCR,cellsizeAUCFeaturesSCR,cellsizeMAXFeatures,cellsizeMAXFeaturesSCR,
									  cellsizetCumuAUCitlesSCR,cellsizeCumuAUCFeaturesSCR,cellsizeCumuMAXFeatures,cellsizeCumuMAXFeaturesSCR,cellsizetCumuAUCitles,cellsizeCumuAUCFeatures,
									  cellsizetitles,cellsizeFeatures,poftitlesC, posfeaturesConfocal,allTs,allFs, suffixes, prefixes,nsc,cellsizetAUCitles,cellsizeAUCFeatures,
									  cellsizetitlesSCR,cellsizeFeaturesSCR ,ratitles={},ratfeatures={},threshTitles,threshFeatures,threshTitlesSCR={},threshFeaturesSCR={}},
			(* some definitions *)						  
			suffixes = If[scr,{" - shr"," - scr", " - nuclear size"," - norm nuclear size"},{" - shr", " - nuclear size"," - norm nuclear size"}];
			prefixes = If[scr,{"time ", "shr ","scr ","nuclear size "},{"time ", "shr ","nuclear size "}];
			nsc=If[scr,4,3];
			(* build cell identitites and metadata rows*)
			initTitles = If[sep,{"exp","file","cell","dex","ind","div","timetodiv"},{"exp","file","cell","dex","ind","newdiv"(*,"div"*),"timetodiv"}];
			initFeatures = If[sep,Table[Flatten[{a[[;;5]],a[[8;;]]}]&@a,{a,dat}],Table[Flatten[{a[[;;5]],a[[8]],a[[10;;]]}]&@a,{a,dat}]];
			
			(* max rate features*)
			rftitlesC= Table[#<>q&/@Flatten[{{Table[w <> "at max rate",{w,prefixes}]},{"max rate"}}],{q,suffixes}]//Flatten;
			
			ratesFeaturesConfocal=If[scr,Table[Flatten[{rates[a[[6]],2, stretch],rates[a[[6]],3, stretch], rates[a[[6]],4, stretch], rates[a[[7]],4, stretch]}],{a,dat}],
										 Table[Flatten[{rates[a[[6]],2, stretch],rates[a[[6]],3, stretch],rates[a[[7]],3, stretch]}],{a,dat}]];
			
			(* AUC features*)
			iftitlesC= Table[#<>q&/@{"AUC"},{q,suffixes[[;;-3]]}]//Flatten;
			IntFeaturesConfocal=If[scr,Table[Flatten[{auc[a[[6,;;,{1,2}]]],auc[a[[6,;;,{1,3}]]]}],{a,dat}],
									   Table[Flatten[{auc[a[[6,;;,{1,2}]]]}],{a,dat}]];
			
			(*mean,std and cv features*)		   
			sftitlesC = Table[#<>q&/@{"mean","std","cv"},{q,suffixes}]//Flatten;
			statFeaturesConfocal=If[scr, Table[Flatten[{ stat@a[[6,;;,2]],stat@a[[6,;;,3]],stat@a[[6,;;,4]],stat@a[[7,;;,4]]}],{a,dat}],
										 Table[Flatten[{ stat@a[[6,;;,2]],stat@a[[6,;;,3]],stat@a[[7,;;,3]]}],{a,dat}]];

			(*quantile features for shr and scr*) 
			tlftitlesC= Table[#<> " " <>ToString@aa<>q&/@{"quantile","first time at quantile"},{q,suffixes[[;;-3]]},{aa,{0.5,0.95}}]//Flatten;
			timelevelsFeaturesConfocal=If[scr,Table[Flatten[{quantiles[a[[6]],2,{0.5,0.95}],quantiles[a[[6]],3,{0.5,0.95}]}],{a,dat}],
											  Table[Flatten[{quantiles[a[[6]],2,{0.5,0.95}]}],{a,dat}]];
			(*quantile features for nuclear size*)
			csftitlesC= Table[#<> " " <>ToString@aa<>q&/@{"quantile","first time at quantile"},{q,suffixes[[-2;;]]},{aa,{0.3,0.5,0.7,0.95}}]//Flatten;
			cellsizelevelsFeaturesConfocal=Table[Flatten[{quantiles[a[[6]],nsc,{0.3,0.5,0.7,0.95}],quantiles[a[[7]],nsc,{0.3,0.5,0.7,0.95}]}],{a,dat}];
			
			
			(*these feature are calculated only for the separated datasets*) 
			If[sep,
				(*find the nuclear size at first time shr crosses a threshold*) 
				cellsizetitles = {Table["norm nuclear size at first shr level of " <> ToString@sh,{sh,shrThs}]}//Flatten;
				cellsizeFeatures = Table[Flatten[{Table[If[sh>Max@a[[6,;;,2]],0,FirstCase[({a[[6,;;,2]],a[[7,;;,nsc]]}\[Transpose]),_?(#[[1]]>= sh&)][[2]]],{sh,shrThs}]}],{a,dat}];
				
				(*shr AUC in a given nuclear size window*)							  
				cellsizetAUCitles = {Table["AUC shr levels at norm nuclear window of " <> 
																				 ToString@cc,{cc,{0.25,0.5,0.75,1}}]}//Flatten;
				cellsizeAUCFeatures = Table[Flatten[{Table[If[cc>Max@a[[7,;;,nsc]] || cc<Min@a[[7,;;,nsc]],0,
										NIntegrate[Interpolation[a[[6,;;,;;2]]][t],{t,FirstCase[({a[[6,;;,1]],a[[7,;;,nsc]]}\[Transpose]),_?(#[[2]]>= cc -0.25 &)][[1]],FirstCase[({a[[6,;;,1]],a[[7,;;,nsc]]}\[Transpose]),_?(#[[2]]>= cc&)][[1]]}]],{cc,{0.25,0.5,0.75,1}}]}],{a,dat}];
				
				(*Max shr in a given nuclear size window*)						
				cellsizetMAXitles = {Table["Max shr levels at norm nuclear window of " <> 
																				 ToString@cc,{cc,{0.25,0.5,0.75,1}}]}//Flatten;
				cellsizeMAXFeatures = Table[Flatten[{Table[If[cc>Max@a[[7,;;,nsc]] || cc<Min@a[[7,;;,nsc]],0,
										Max[Max[Select[a[[6]],FirstCase[({a[[6,;;,1]],a[[7,;;,nsc]]}\[Transpose]),_?(#[[2]]>= cc -0.25 &)][[1]]<= #[[1]]<=FirstCase[({a[[6,;;,1]],a[[7,;;,nsc]]}\[Transpose]),_?(#[[2]]>= cc&)][[1]]&][[;;,2]]],0]],{cc,{0.25,0.5,0.75,1}}]}],{a,dat}];
			
		
				(*find the nuclear size at first time scr crosses a threshold*)
				If[scr,
						cellsizetitlesSCR = {Table["norm nuclear size at first scr level of " <> ToString@sh,{sh,shrThs}]}//Flatten;
						cellsizeFeaturesSCR = Table[Flatten[{Table[If[sh>Max@a[[6,;;,3]],0,FirstCase[({a[[6,;;,3]],a[[7,;;,nsc]]}\[Transpose]),{_?(#>= sh&),_}][[2]]],{sh,shrThs}]}],{a,dat}];,
												cellsizetitlesSCR={};	
												cellsizeFeaturesSCR=Table[{},{a,dat}];
													  ];
					
				If[scr,
				(*scr AUC in a given nuclear size window*)
							cellsizetAUCitlesSCR = {
												Table["AUC scr levels at norm nuclear window of " <> 
																					 ToString@cc,{cc,{0.25,0.5,0.75,1}}]}//Flatten;
							cellsizeAUCFeaturesSCR = Table[Flatten[{Table[If[cc>Max@a[[7,;;,nsc]] || cc<Min@a[[7,;;,nsc]],0,
												NIntegrate[Interpolation[a[[6,;;,{1,3}]]][t],{t,FirstCase[({a[[6,;;,1]],a[[7,;;,nsc]]}\[Transpose]),_?(#[[2]]>= cc -0.25 &)][[1]],FirstCase[({a[[6,;;,1]],a[[7,;;,nsc]]}\[Transpose]),_?(#[[2]]>= cc&)][[1]]}]],{cc,{0.25,0.5,0.75,1}}]}],{a,dat}];
				(*Max scr in a given nuclear size window*)	
							cellsizetMAXitlesSCR = {Table["Max scr levels at norm nuclear window of " <> 
																				 ToString@cc,{cc,{0.25,0.5,0.75,1}}]}//Flatten;
				
							cellsizeMAXFeaturesSCR = Table[Flatten[{Table[If[cc>Max@a[[7,;;,nsc]] || cc<Min@a[[7,;;,nsc]],0,
														Max[Max[Select[a[[6]],FirstCase[({a[[6,;;,1]],a[[7,;;,nsc]]}\[Transpose]),_?(#[[2]]>= cc -0.25 &)][[1]]<= #[[1]]<=FirstCase[({a[[6,;;,1]],a[[7,;;,nsc]]}\[Transpose]),_?(#[[2]]>= cc&)][[1]]&][[;;,3]]],0]],{cc,{0.25,0.5,0.75,1}}]}],{a,dat}];
				
										
											,
											cellsizetAUCitlesSCR={};	
											cellsizeAUCFeaturesSCR=Table[{},{a,dat}];
											cellsizetMAXitlesSCR={};	
											cellsizeMAXFeaturesSCR=Table[{},{a,dat}];
											cellsizetCumuAUCitlesSCR={};	
											cellsizeCumuAUCFeaturesSCR=Table[{},{a,dat}];
											cellsizetCumuMAXitlesSCR={};	
											cellsizeCumuMAXFeaturesSCR=Table[{},{a,dat}];
											
				
												  ];,
											cellsizetitles={};	
											cellsizeFeatures=Table[{},{a,dat}];	  
											cellsizetAUCitles={};	
											cellsizeAUCFeatures=Table[{},{a,dat}];
											cellsizetMAXitles={};	
											cellsizeMAXFeatures=Table[{},{a,dat}];
											cellsizetCumuAUCitles={};	
											cellsizeCumuAUCFeatures=Table[{},{a,dat}];
											cellsizetCumuMAXitles={};	
											cellsizeCumuMAXFeatures=Table[{},{a,dat}];
											cellsizetitlesSCR={};	
											cellsizeFeaturesSCR=Table[{},{a,dat}];
											cellsizetAUCitlesSCR={};		  
											cellsizeAUCFeaturesSCR=Table[{},{a,dat}];
											cellsizetMAXitlesSCR={};	
											cellsizeMAXFeaturesSCR=Table[{},{a,dat}];
											cellsizetCumuAUCitlesSCR={};	
											cellsizeCumuAUCFeaturesSCR=Table[{},{a,dat}];
											cellsizetCumuMAXitlesSCR={};	
											cellsizeCumuMAXFeaturesSCR=Table[{},{a,dat}];
												  ];

			(* correlation between shr and scr*) 		    
		    ratitles = If[scr,Table[{"cor" <> q},{q,{" shr / scr"}}]//Flatten,{}];
			ratfeatures = If[scr,Table[Flatten[{					
									Correlation[a[[6,;;,2]],a[[6,;;,3]]]
									}]
								  ,{a,dat}]/.Correlation[__,__]->0
										       
							,Table[{},{a,dat}]];
			(* crossing shr threshold*)
			threshTitles = {"norm nuclear size at shr threshold","time from theshold to end"};
			threshFeatures = Table[Flatten@If[thresh>Max@a[[6,;;,2]],{0,0},{#[[1]],a[[6,-1,1]]-#[[2]]}&@FirstCase[({a[[6,;;,2]],a[[7,;;,nsc]],a[[6,;;,1]]}\[Transpose]),_?(#[[1]]>= thresh&)][[{2,3}]]],{a,dat}];
			
			(* crossing scr threshold*)
			If[scr,
				threshTitlesSCR = {"norm nuclear size at scr threshold"};
				threshFeaturesSCR = Table[Flatten@If[threshSCR>Max@a[[6,;;,3]],{0},FirstCase[({a[[6,;;,3]],a[[7,;;,nsc]]}\[Transpose]),_?(#[[1]]>= threshSCR&)][[{2}]]],{a,dat}];,
										threshTitlesSCR={};	
										threshFeaturesSCR=Table[{},{a,dat}];
			];
			
			(*cell position *) 
			poftitlesC = {"cell pos"};
			posfeaturesConfocal = Table[Flatten[{ToExpression@StringTake[a[[3]],{5}]}],{a,dat}];
			
			(* create unified objects for all titles and another for all features*) 
			allTs = Flatten@ {initTitles,rftitlesC,iftitlesC,sftitlesC,tlftitlesC,poftitlesC,csftitlesC,cellsizetitles,cellsizetitlesSCR,ratitles,threshTitles,threshTitlesSCR,cellsizetAUCitles,cellsizetAUCitlesSCR,cellsizetMAXitles,cellsizetMAXitlesSCR};
			allFs = N[Flatten/@({initFeatures,ratesFeaturesConfocal,IntFeaturesConfocal,statFeaturesConfocal,
			timelevelsFeaturesConfocal,posfeaturesConfocal,cellsizelevelsFeaturesConfocal,cellsizeFeatures,cellsizeFeaturesSCR,ratfeatures,threshFeatures,threshFeaturesSCR,cellsizeAUCFeatures,cellsizeAUCFeaturesSCR,cellsizeMAXFeatures,cellsizeMAXFeaturesSCR}\[Transpose])]/.{Indeterminate->0,ComplexInfinity->0};
			{allTs,allFs}];
			
(*a distance measure based on correlation (0 is correlated and 1 is not correlated *)
distCor[x_,y_]:=Chop@(1-Abs@Correlation[x,y])

(*calculate the best threshold that separates the feature data into divided and undivided cells
	inputs: 
			ruleTrain -> rule (lookup table) for training set  (feature -> dividing/non-dividing) 
			val -> threshold resolution 
	output: returns the best threshold
			*)
	
class4oneFeature[ruleTrain_,val_:3000]:= Block[{coms,stds,dat,V=val,c1,c2,tab},
									(*separate data into divided and non-divided *)
									dat=SortBy[GatherBy[ruleTrain, #[[2]]&],#[[1,2]]&][[;;,;;,1]];
									(* calculate Center of Mass and sd for each group *) 
									coms = Mean/@dat;
									stds = StandardDeviation/@dat;
									(* prepare range of values to test the thresholds from the smallest com - 2 sds to the largest com + 2 sds *) 
									If[coms[[2]] > coms[[1]], 
											c1 = coms[[1]]- (2 stds[[1]]); c2 = coms[[2]]+ (2 stds[[2]]);								
											(* calculate the prediction score for every potential theshold and return the one with the highest score*) 
											tab = Table[
														{a,Total[Join[Boole[#==1]&/@(Boole@Map[# <= a&,dat[[1]]]),Boole[#==1]&/@(Boole@Map[# > a&,dat[[2]]])]]},
														{a,c1,c2,(c2 - c1)/V}];			
{											SortBy[tab,#[[2]]&][[-1,1]],1},

											 c1 = coms[[1]]+2 stds[[1]]; c2 = coms[[2]]- 2 stds[[2]];
											 tab= Table[{a,Total[Join[Boole[#==1]&/@(Boole@Thread[dat[[1]] >= a]),Boole[#==1]&/@(Boole@Thread[dat[[2]] < a])]]},
														{a,c2,c1,(c1 - c2)/V}];
													
										 {SortBy[tab,#[[2]]&][[-1,1]],-1}]];

(*calculate accuracy of a threshold for a feature 
	inputs: 
			ruleTest -> rule (lookup table) for testing set  (feature -> dividing/non-dividing)
			thresh -> threshold to test
	output: accuracy score
			*)																					
accuracy[ruleTest_,thresh_]:= Block[{l = Length@ruleTest, m},
									m = Total[Boole[Boole[If[thresh[[2]]>0, #[[1]] > thresh[[1]],#[[1]]<thresh[[1]]]]==#[[2]]]&/@ruleTest];
									N@m/l];


(* ::Text:: *)
(*Functions that search for optimal threshold and plot it*)


(*internal function that runs over a range of potential thresholds and calculates the prediction accuracy for crossing it for each nuclear size bin:
	inputs: dat -> dataset in wn format 
			bins -> # of nuclear size windows to use - in the paper we used 4 
			col -> 2 for SHR 3 for SCR
			range -> the set of thresholds to test
			norm -> a flag - whether to normalize the nuclear size
	output: 
			the percentage of accurate predictions for each threshold for each nuclear size window*)
thresholds[dat_,bins_,col_,range_,norm_]:=Block[{p=N@1/bins,nn=If[norm,normalize,Identity]},
												100. N@Total/@#/Length/@#&/@Table[
																	Table[
																		Boole[Equal@@{Boole[(th2 - p)<= #[[Quiet[FirstPosition[#[[;;,2]],_?(#> th&)][[1]]],-1]]< th2]/.Boole[_]-> 0&@
																		({#[[6,;;,1]],#[[6,;;,col]],nn@#[[7,;;,-1]]}\[Transpose]),#[[-2]]/.{"A" -> 1, "S" -> 0, "N" -> 0}}]&/@dat,{th2,p,1,p}],{th,range}]];
																		
(*get a cell's SHR or SCR trajectory and predict whether it will divide or not given a nuclear size window and a threshold:
	inputs: row -> a cell from a wn formatted dataset
			col -> 2 for SHR 3 for SCR
			th -> threshold to cross
			th2 -> upper limit of nuclear size window
			p  -> nuclear size window length
			norm -> a flag - whether to normalize the nuclear size
	output: 
			a boolean value whether the cell is preticted to divide or not*)																		
predictions[row_,col_,th_,th2_,p_, norm_]:= (Boole[((th2 - p))<=  #[[FirstPosition[#[[;;,2]],_?(#> th&)][[1]],-1]]< th2]/.Boole[_]-> 0)&@({row[[6,;;,1]],row[[6,;;,col]],If[norm,normalize@#,Identity@#]&@N@row[[7,;;,-1]]}\[Transpose])

(*plot the threshold plot (fig 3d in paper):
	inputs: path -> path to save the image
			range -> the x axis values - thresholds
			per -> prediction accuracies
			opAll -> number of nuclear size windows
	output: 
			a file of the plot in path*)																		
plotThreshold[path_,range_,per_,opAll_]:= Block[{p=N@1/opAll},Export[path,ListLinePlot[{range,Flatten[#[[1]]]}\[Transpose]&/@({per\[Transpose],Range[p,1,p]}\[Transpose]),PlotRange->{All,{20,100}},PlotStyle->Thickness[thicnkess],Ticks->{{0.1,0.2,0.3,0.4,.5,.6},{25,50,75,100}},AxesStyle->framestyle,LabelStyle->label,ImageSize->defaultImageSize]]]

(*runs over a range of potential thresholds and calculates the prediction accuracy for crossing it for each nuclear size bin and exports the data files associated with it:
	inputs: dat -> dataset in wn format 
			opAll -> number of nuclear size windows
			col -> 2 for SHR 3 for SCR
			range -> the set of thresholds to test
			rule -> the set of rules that link cells with min and 0.9 quantile values to use for normalization
			path -> path to save the images
			prefix -> initial string for file names
			sep -> a flag that specied if the data is separated or not 
			export -> whether to export files
	output: 
			the percentage of accurate predictions for each threshold for each nuclear size window as well as 3 csv files of the individual cell prediction, stats and the percentages. Figures for paper (3d) *)
plotThresholdCurves[dat_,opAll_,col_,range_,rule_,path_,prefix_,sep_,export_:True]:= Block[{precentagesTh2WinF={},precentagesTh2WinFN={},
					dataForPercentages,titles,ftitles,dfIn,dfIn2,dfStats, titlesS,p=N@1/opAll,dataForIncorrect,ret},
					(* titles for tables to be exported later*)
																   											   
					titles = Flatten[{"Exp","File","cell","Full_min_cell_size","Full_p9Q_cell_size","Sep_min_cell_size","Sep_max_cell_size",
									  "div prediction for NS window "<> #&@ToString@#&/@Range[p,1,p],"div"}];
																									  
					titlesS = Flatten[{"Exp","File","cell","Full_min_cell_size","Full_p9Q_cell_size","Sep_min_cell_size","Sep_max_cell_size","div prediction for NS window with Full range NS"<> #&@ToString@#&/@Range[p,1,p],
					   				 "div prediction for NS window with Sep range NS"<> #&@ToString@#&/@Range[p,1,p],"div"}];
																					
					(* the percentage of accurate predictions for each threshold for each nuclear size window *)						
					precentagesTh2WinF=thresholds[dat,opAll,col,range,False];
										
					If[sep, precentagesTh2WinFN= thresholds[dat,opAll,col,range,True]; ,{}];
					
					(* runs over each cell and calculates the prediciton of that cell for each bin -> to export later in csv format *)														
					dataForPercentages=Block[{vec,
											 vecT=range[[Position[#,Max@#][[-1,1]]]]&/@(precentagesTh2WinF\[Transpose]),
											 vecNT},
											If[sep, vecNT=range[[Position[#,Max@#][[-1,1]]]]&/@(precentagesTh2WinFN\[Transpose]);,{}];
											
											vec =Table[  th2,{th2,p,1,p}];
											Print@vecT;
											Flatten@{#[[;;3]],{#[[1]],#[[2]], If[sep,#[[3]],StringTake[#[[3]],5]]}/.rule, {Min@#,Quantile[#,0.9]}&@#[[6,;;,-1]],
													Table[ predictions[#,col,vecT[[th2]],vec[[th2]],p ,False] ,{th2,1,opAll}],
													If[sep, Table[ predictions[#,col,vecNT[[th2]],vec[[th2]],p ,True] ,{th2,1,opAll}] ,{}], #[[8]]}&/@dat];
					
					(* calculate stats*)																	
					dfIn =Table[Block[{lenn,sdat,nd,d,wrper},
							sdat = Select[dataForPercentages[[;;,{1,2,3,c,-1}]],#[[4]]!= #[[5]]&];
							lenn= Length@sdat;
							{nd,d} = {Cases[sdat,{___,0}],Cases[sdat,{___,1}]}[[;;,;;,;;3]];
							wrper=(100.0*{Length@nd,Length@d})/lenn;
							{nd,d,wrper,100.0*lenn/Length@dataForPercentages}],{c,8,Length@dataForPercentages[[1]]-1}];
					
											
																												
					dfStats = {#[[2]],#[[1,1]],#[[1,2]]}&/@dfIn[[;;,3;;]];
					
					(*choose titles*)																											
					ftitles = If[sep,titlesS,titles];
					
					(* export tables*) 
					If[export,
					Export[path<>"/data_thresholds_"<>prefix<>ToString@opAll<>"_bins "<>DateString[]<>".csv",Prepend[dataForPercentages, ftitles]];		
					Export[path<>"/data_stats_"<>prefix<>ToString@opAll<>"_bins "<>DateString[]<>".csv",Prepend[Prepend[#[[1]],#[[2]]]&/@({dfStats, ftitles[[8;;-2]]}\[Transpose]),{"col","%badlyPredicted","%not_div but pred to div","%div but pred not to div"}]];
				
					
					ret={precentagesTh2WinF,precentagesTh2WinFN},
					ret= {precentagesTh2WinF,precentagesTh2WinFN,Prepend[dataForPercentages, ftitles]} ;
					];
					ret
					]

	(*calculate the threshold for each bin that bests predict the cell fates
	inputs: dat -> dataset in wn format 
			opAll -> number of nuclear size windows
			col -> 2 for SHR 3 for SCR
			range -> the set of thresholds to test
			sep -> a flag that specied if the data is separated or not 
	output: 
			the threshold that yield the higher accuarcy for each window *)								
bestThresholds[dat_,opAll_,col_,range_,sep_]:= Block[{precentagesTh2WinF,precentagesTh2WinFN,optimal,vecT,vecNT={},p=N@1/opAll,dataForIncorrect},
									precentagesTh2WinF=thresholds[dat,opAll,col,range,False];
									If[sep,precentagesTh2WinFN= thresholds[dat,opAll,col,range,True];  ,{}];
									vecT=range[[Position[#,Max@#][[-1,1]]]]&/@(precentagesTh2WinF\[Transpose]);  
									If[sep,
										 vecNT=range[[Position[#,Max@#][[-1,1]]]]&/@(precentagesTh2WinFN\[Transpose]); ,{}];
									{vecT,vecNT}

																						
																				];


(* ::Text:: *)
(*ODE parameter fitting related functions*)


(*This function is an adaptation of the algorithm for parameter fitting of gradient descent that appears in the following blog: 
 https://dpananos.github.io/posts/2019-05-21-odes/
 We use as input a given measurement and try to fit the response to it 
 inputs:
        eqs -> ODEs to best fit parameters with respect to variables
        varsLoss -> variables that are used to calculate lossses (SCR in our case)
        inputName ->  variable that needs to get value from external sources (SHR in our case)
        params -> a list of parameter names to fit 
        dat -> data of shr and scr of a cell
        init-> initial condition equations for ODE
        R -> scaling factor for regularization of parameter magnitudes
        \[Lambda]In -> initial step size of gradient descent
        paramsI-> initial guess for parameters - must be the same length as parameters
        ind -> index of cell (usefull if running this in a table or cluster)
        qqq -> max number of iterations to run the algorithm
        SCRM -> a factor to scale the data to the output of the model
  output:
        the following structures for the best loss reached: 
        {ind,iteration,best-fit parameters values ,best loss reached ,step size at this point}
        
 *)
gradODEs[eqs_,varsLoss_,inputName_,params_,dat_,init_,R_,\[Lambda]In_,paramsI_,ind_,qqq_,SCRM_:1.0]:= Module[{rhs,lhs,J,dfdp,sensi,eqsSensi,convergence={},pp,vars},
				(* calcualte Jacobian - the gradient matrix of the ODEs with respect to the variables used to calcualte the losses*)
				J= Outer[D[#1[[2]],#2]&,eqs[[{1}]],varsLoss]//FullSimplify;
				(* calcualte the gradient matrix of the ODEs with respect to the parameters*)
				dfdp = Outer[D[#1[[2]],#2]&,eqs[[{1}]],params]//FullSimplify;
				(*build new variable to calculate gradients with original ODEs*)			    
				sensi= Table[ToExpression[i<>ToString@#<>"sens[t]"]&/@params,{i,StringDelete[#,"[t]"]&/@(ToString/@varsLoss)}];
				(*build new ODEs to run with original ODEs*)			    
				eqsSensi = Flatten[Thread/@Thread@(D[sensi,t]== J . sensi + dfdp)]; 
				vars = Flatten[Join[varsLoss,sensi]];
				
				Block[{cS,cC,Flag2,\[Epsilon]=10^-6,timeVals,\[Lambda]=\[Lambda]In,curr={dat[[6]],ind}},
						(*shr *)
						cS = curr[[1,;;,{1,2}]]+\[Epsilon];
						(*scr *)
						cC = {curr[[1,;;,1]],curr[[1,;;,3]]/SCRM}\[Transpose]+\[Epsilon];
						(*time *)
						timeVals= curr[[1,;;,1]]+\[Epsilon];
						
						(*initialization of results table*) 
						pp =paramsI;
						convergence={};
						Flag2=False;
						
						Table[Block[{ss,hh,grad,loss,tem,ll,gradall={},calcC,lossall={},lossMall={},e,sq,fac=1.4,gg,positions,bssf,flag = True,ppn,equations2Solve,bestIC},
							If[Not@Flag2,
								gradall={};
								lossall={};
								(*find initial codition as steady state when there is no input*) 
								bestIC = #1==Re@#2&@@@(Select[Evaluate[N[((init)/.Thread[params->pp]/.t->Min@timeVals)]],
										 And@@Thread[#[[;;,2]]>=0]&][[1]]);
								
								(*prepare and solve ODEs and check for convergance*)	 	 
								equations2Solve = Flatten@Join[
										Join[#1==Re@#2&@@@(eqsSensi/.Thread[params->pp]),{#1==Re@#2&@@@eqs,
												inputName==Evaluate[Interpolation[cS][t]]}/.Thread[params->pp]],{bestIC,
												#1==Re@#2&@@@DeleteDuplicates@Thread[Flatten[(sensi/.t->Min@timeVals)]==0]/.Thread[params->pp]}];

								ss= Check[NDSolve[equations2Solve,vars,{t,Min@timeVals,Max@timeVals}],-98];
								If[ss==-98,Flag2=True];
								
								(* calculate scr predictions according to model*) 
						        calcC = (varsLoss/.#1/.t->timeVals[[2;;]])[[1]]&@@ss;
						        (* calculate differences between prediction and data withotu first time point *) 
						        hh=calcC- cC[[2;;,2]];
								(* calculate gradients and loss in parameter space*)
						        gg= (Table[sensi/.#,{t,timeVals[[2;;]]}][[;;,1]]\[Transpose])&@@(ss);
						        gradall= # . hh&/@gg +2R (Total[Abs@pp]);
							    (* calculate loss as RMS with regularization*)
							    lossall= Sqrt[Mean[((varsLoss/.#1/.t->timeVals[[2;;]])[[1,1]]-#2[[2;;,2]])^2]]+R (Total[pp^2])&@@({ss,cC});
							(* adapt step size according to loss gain *) 									
							If[(convergence//Length )>1,
								If[convergence[[-1,-2]]<lossall,flag=False; \[Lambda]/=fac^8,\[Lambda]*=fac]
							,""];
							
							(* update parameter values and cap at 10^-9 if loss got better*) 
							bssf = If[(convergence//Length )>1,SortBy[convergence,#[[4]]&][[1]],{0,0,0}];
							If[flag,ppn =pp- Re[\[Lambda] gradall],ppn=bssf[[3]]];
							ppn = Max[#,10^-9]&/@Abs@ppn;
							
							(*add result to results table*)
							AppendTo[convergence,{ind,q,pp,lossall,\[Lambda]}];
							pp =ppn;
							(*cap stepsize*)
							If[\[Lambda]>1,\[Lambda]=1 ,If[\[Lambda]<10^-11,\[Lambda]=10^-11]];
							
							(*finish running if loss convereged*)			
							If[Length[convergence]>70,
								If[And@@Thread[Abs@Differences@convergence[[-10;;,-2]]<7 10^-6],Flag2=True],""];
							];
							If[Mod[q,500]==0,Print@q];
								],{q,qqq}]
							];
							Print[StringRiffle@{"time is:", LocalTime[],"curve is:",ind}];
							(* output best score*) 
							SortBy[convergence,#[[-2]]&][[1]]
						
							];
							
(*Calculates the predicted curve (scr) given eqs parameters and input (shr)
 inputs:
        eqs -> ODEs to best fit parameters with respect to variables
        varsI -> variables that are used as outputs (SCR in our case)
        inputName ->  variable that needs to get value from external sources (SHR in our case)
        params -> a list of parameter names to fit 
        dat -> data of shr and scr of a cell
        init-> initial condition equations for ODE
        R -> scaling factor for regularization of parameter magnitudes
        \[Lambda]In -> initial step size of gradient descent
        paramsI->  parameter values - must be the same length as parameters
        ind -> index of cell (usefull if running this in a table or cluster)
        factor -> a factor to scale the input data
        SCRM -> a factor to scale the data to the output of the model
  output:
        a list of pairs: {time, prediction}
        
 *)						
CalcCurveODEs[eqs_,varsI_,inputName_,params_,dat_,init_,paramsI_,ind_,factor_,SCRM_]:= 
				Module[{cS,cC,Flag2,\[Epsilon]=10^-6,timeVals,curr={dat[[6]],ind}},
						cS = {curr[[1,;;,1]],factor curr[[1,;;,2]]}\[Transpose]+\[Epsilon];
						timeVals= curr[[1,;;,1]]+\[Epsilon];
						Flag2=False;
						Block[{ss,hh,grad,loss,tem,ll,gradall={},lossall={},lossMall={},e,sq,fac=1.2,gg,positions,bssf,flag = True,ppn,equations2Solve,bestIC},
								
							bestIC = #1==Re@#2&@@@(Select[Evaluate[N[((init)/.Thread[params->paramsI]/.t->Min@timeVals)]],
										 And@@Thread[#[[;;,2]]>=0]&][[1]]);
									
						
							equations2Solve = Flatten@Join[
										Join[#1==Re@#2&@@@(eqs/.Thread[params->paramsI]),{
												inputName==Evaluate[Interpolation[cS][t]]}/.Thread[params->paramsI]],
												bestIC];
							
							ss= Check[NDSolve[equations2Solve,varsI,{t,Min@timeVals,Max@timeVals}],-98];
							{timeVals,(varsI/.#1/.t->timeVals)[[1]]&@@ss}\[Transpose]
							
								]
							]
(* calculates \[Chi]^2 of observed (o) and expected (e) curves*)
chisq[o_,e_] := Total[(o-e)^2/(e+0.001)]
(* calculates R^2 of observed (o) and expected (e) curves*)
Rsq[o_,e_] :=((N@((Length[o] o . e - Total[o] Total[e])/(Sqrt[Length[o] o . o - Total[o]^2]Sqrt[Length[e] e . e - Total[e]^2]+10^-9)))^2)  


(* ::Subsection:: *)
(*create directories to store the outputs*)


CreateDirectory[finalFiguresFolder];
subpaths  =finalFiguresFolder <> "/" <> #&/@subfoldernames;
CreateDirectory/@subpaths;


(* ::Section:: *)
(*Load and preprocess data*)


(* ::Subsection:: *)
(*Confocal data*)


(*Load and remove data from after asymmetric division *)
ConfocalRawData = Import[basePath <> confFileName, "CSV"];
ConfocalRawData = Select[ConfocalRawData,StringPosition[#[[5]],"endo"|"cor"][[;;,1]]=={}&];

(*Load nuclear size corrected data*)
nsNew = Import[basePath <> nsPath];
(* Load and extract the time where dex was introduced with respect to experiment inital measurements*)
dexStartConfocal = {#1, QuantityMagnitude @ UnitConvert[#, "h"]& @ (DateObject[
    ToExpression /@ StringSplit[#2, "-"]] - DateObject[ToExpression /@ StringSplit[
    #3, "-"]])}& @@@ (DeleteCases[Import[basePath <> confmetaFileName,
     "CSV"], {""..}][[2 ;; , {2, 6, 7}]] /. "2018-17-10-15-24-43" -> "2018-07-10-15-24-43"
    );

(* separate titles from data*)
{ConfocalRawDataTitles, ConfocalRawData} = {ConfocalRawData[[1]], ConfocalRawData
    [[2 ;; ]]};

(* filter table to relevant columns *)
ConfocalDex = ConfocalRawData[[;;,{2, 3, 4, 5, 6, 7, 8, 9, 10, 11, 12, 13, 16, -3}]];
ConfocalDexTitles = Append[ConfocalRawDataTitles[[{2, 3, 4, 5, 6, 7, 
    8, 9, 10, 11, 12, 13, 16}]], "Dex"];
(*add nuclear size to data  *)
nsNewRule = Dispatch[#[[ ;; 5]] -> {Max[#[[-2;;]]]}& /@ nsNew[[]]];
ConfocalDexNs = (Join[#, #[[{1, 2, 3, 4, 8}]] /. nsNewRule]& /@ ConfocalDex);
ConfocalDexNs = Select[ConfocalDexNs, Length@#==15&];
(*background subtraction of SHR-GFP using mean of first three points*)
backGreenRuleConf= #[[1,{1,2,3}]]-> Mean@#[[;;3,5]]&/@(SortBy[#,#[[8]]&]&/@GatherBy[ConfocalDexNs,#[[1]]||#[[2]]||#[[3]]&]);
(*background and bleedtrhough correction *)
InterimBackSubtractedConfocalDex= {Max[#[[1]],0],Max[#[[2]],0],Max[#[[1]]/#[[2]],0]}&@((IdentityMatrix[2]-BTEstimatesConfocal) . ({#5,#6}-{({#1,#2,#3}/.backGreenRuleConf),
								  AutoFluorescenceEstimatesConfocal[[2]]}))&@@@ConfocalDexNs;
BackSubtractedConfocalDex=Join[#[[1,;;4]],#[[2]],#[[1,8;;]]]&/@({ConfocalDexNs,InterimBackSubtractedConfocalDex}\[Transpose]);

(* rule for end of time course for each cell*)
endroundrule=DeleteDuplicates[({#[[1]],#[[2]],#[[4]]}-> #[[10]]/4+(#[[1]]/.Rule@@@dexStartConfocal))&/@BackSubtractedConfocalDex]/._+("NA")/4->"NA";

(*rule for asymmetric divisions has cell identifiers and a flag that is true if cells ends up dividing asymmetrically *)
asymConfocal = Flatten/@({#[[1,{1,2,4}]],Or@@(#!= "NA"&/@#[[;;,5]])}&/@GatherBy[DeleteDuplicates@ConfocalDex[[;;,{1,2,3,4,-5,-4,-3}]],#[[1]]&&#[[2]]&&#[[4]]&]);

(* collects the data into trajectories and change the last H2B values (1 or 2) to be like the average of the H2B values just before that and recalculate the h2b normalization*)
gatheredDataConfocal = Table[Flatten[
							If[#[[1,12]]== "NA" && Length@#>5,
								Join[#[[;;6]],{#[[5]]/#[[6]]},#[[8;;]]]&/@
									Join[#[[;;-3]],{Join[#[[-2,;;5]],{Mean[#[[-6;;-4,6]]]},#[[-2,7;;]]],Join[#[[-1,;;5]],{Mean[#[[-5;;-3,6]]]},#[[-1,7;;]]]}],
								If[#[[1,12]]== "NA" && Length@#>1,
									Join[#[[;;6]],{#[[5]]/#[[6]]},#[[8;;]]]&/@
										Join[#[[;;-2]],{Join[#[[-1,;;5]],{#[[-2,6]]},#[[-1,7;;]]]}],
									#]
							]&/@GatherBy[d,#[[4]]&],1],{d,GatherBy[BackSubtractedConfocalDex,#[[1]]||#[[2]]||#[[3]]&]}];

(* change trajectory's cell position into last generation and filter out trajectories with less than 11 timepoints*)
daughtercellsConfocal= Table[Select[gd,MemberQ[isnotContained@Flatten[DeleteDuplicates@gd[[;;,4]]],#[[4]]]&],{gd,compTraj/@gatheredDataConfocal}];
longTimesConfocal = MapIndexed[{SortBy[#1,#[[8]]&],#2[[1]]}&,Select[GatherBy[Flatten[daughtercellsConfocal,1],#[[1]]||#[[2]]||#[[4]]&],Length@#>10&]];


(*organize into cells table with the first 2 and last 2 timepoints removed. the data appears in smoothed and unsmoothed versions*)
organizedDataConfocal=Parallelize[{#1,#2,#3,#4,#5,{(#6[[1]]+#1/.Rule@@@dexStartConfocal),smooth[#6[[2]],7],smooth[#6[[3]],7]}\[Transpose],
								 {(#6[[1]]+#1/.Rule@@@dexStartConfocal),#6[[2]],#6[[3]]}\[Transpose]}&@@{#[[1,1,1]],#[[1,1,2]],#[[1,1,4]],#[[1,1,-2]],#[[2]],#[[1,3;;-3,{9,7,15}]]\[Transpose]}&/@longTimesConfocal];

(* remove any trajectory with information about what happens after asymmetric division  *)	
dataC= Select[organizedDataConfocal,StringPosition[#[[3]],"endo"|"cor"][[;;,1]]=={}&];

(* add cell fate explicitly into the data &  select cells from newest batch of experiments  *)
dConf =Select[Flatten[#,1]&/@({#,#[[;;3]]/.Thread[{#1,#2,#3}&@@@asymConfocal[[;;,;;3]]-> asymConfocal[[;;,4]]],#[[;;3]]/.endroundrule
		}&/@dataC)/.{False->0,True->1},ToExpression@StringTake[#[[1]],-5]>9050&];

(* prepare the same construct as before but without cell 1 *)		
dConfnc1=Select[Flatten[#,1]&/@({#,#[[;;3]]/.Thread[{#1,#2,#3}&@@@asymConfocal[[;;,;;3]]-> asymConfocal[[;;,4]]],#[[;;3]]/.endroundrule
		}&/@Select[dataC,ToExpression@StringTake[#[[3]],{5}]>1&])/.{False->0,True->1},ToExpression@StringTake[#[[1]],-5]>9050&];

(* prepare list of symmetric divisions *)				
syms = Quiet@Table[Block[{rd =Cases[ConfocalDex,{cell[[1]],cell[[2]],__}],ns=StringDrop[cell[[3]],-1] ,times={}},{While[StringLength[ns]>4,AppendTo[times,SortBy[Select[rd,#[[4]]==ns&
				],#[[9]]&][[-1,9]]];ns=StringDrop[ns,-1]]};cell[[;;3]]->Reverse@times + (cell[[1]]/.Rule@@@dexStartConfocal)],{cell,Select[dConf,StringLength[#[[3]]]>5&][[;;]]}];

(*prepare data for division diagram -- not used in paper*)
predivhydConAll={#1,#3,StringTake[#4,5],#4,#2,{Red,#5},{Blue,#6},{Cyan,#7}}&@@@DeleteDuplicates[ConfocalRawData[[;;,{2,-3,3,5,11,12,13}]]];
divOcAll=DeleteCases[predivhydConAll,{_,"NA"},\[Infinity]][[;;,{1,2,3,4,5,6}]];

(* choose only fully induced roots*)
d10=Select[dConf,#[[4]]==10&];


(* ::Text:: *)
(*prepare data for darker curve in fig1f*)


(*choose cell *)
cell4Fig1=Cases[ConfocalDex,{"Exp09025","lowerleft","cell2",__}];

(* GFP channel background subtraction*)
cell4Fig1bck = #[[1,{1,2,3}]]-> Mean@#[[;;3,5]]&/@(SortBy[#,#[[8]]&]&/@GatherBy[cell4Fig1,#[[1]]||#[[2]]||#[[3]]&]);

(*background subtraction*)
InterimBackSubtractedcf1= {Max[#[[1]],0],Max[#[[2]],0],Max[#[[1]]/#[[2]],0]}&@((IdentityMatrix[2]-BTEstimatesConfocal) . ({#5,#6}-{({#1,#2,#3}/.cell4Fig1bck),
								  AutoFluorescenceEstimatesConfocal[[2]]}))&@@@cell4Fig1;
BackSubtractedcell4Fig1=Join[#[[1,;;4]],#[[2]],#[[1,8;;]]]&/@({cell4Fig1,InterimBackSubtractedcf1}\[Transpose]);

(*set rule for asymmetric divisions*)	
asymfig1 = Flatten/@({#[[1,{1,2,4}]],Or@@(#!= "NA"&/@#[[;;,5]])}&/@GatherBy[DeleteDuplicates@cell4Fig1[[;;,{1,2,3,4,-5,-4,-3}]],#[[1]]&&#[[2]]&&#[[4]]&]);
(* collects the data into trajectories and change the last H2B values (1 or 2) to be like the average of the SHR values just before that and recalculate the h2b normalization term*)
gatheredDatafig1 = Table[Flatten[If[#[[1,12]]== "NA" && Length@#>5,Join[#[[;;6]],{#[[5]]/#[[6]]},#[[8;;]]]&/@Join[#[[;;-3]],
							{Join[#[[-2,;;5]],{Mean[#[[-6;;-4,6]]]},#[[-2,7;;]]],Join[#[[-1,;;5]],{Mean[#[[-5;;-3,6]]]},#[[-1,7;;]]]}],
							If[#[[1,12]]== "NA" && Length@#>1,Join[#[[;;6]],{#[[5]]/#[[6]]},#[[8;;]]]&/@Join[#[[;;-2]],{Join[#[[-1,;;5]],{#[[-2,6]]},#[[-1,7;;]]]}],#]
							]&/@GatherBy[d,#[[4]]&],1],{d,GatherBy[BackSubtractedcell4Fig1,#[[1]]||#[[2]]||#[[3]]&]}];

(* change trajectory's cell position into last generation and filter out trajectories with less than 11 timepoints*)
daughtercellsfig1= Table[Select[gd,MemberQ[isnotContained@Flatten[DeleteDuplicates@gd[[;;,4]]],#[[4]]]&],{gd,compTraj/@gatheredDatafig1}];
longTimesfig1 = MapIndexed[{SortBy[#1,#[[8]]&],#2[[1]]}&,Select[GatherBy[Flatten[daughtercellsfig1,1],#[[1]]||#[[2]]||#[[4]]&],Length@#>10&]];

(*organize into cells table with the first 2 and last 2 timepoints removed. the data appears in smoothed and unsmoothed versions*)
organizedDatafig1={#1,#2,#3,#4,#5,{(#6[[1]]+#1/.Rule@@@dexStartConfocal),smooth[#6[[2]],7]}\[Transpose],
								 {(#6[[1]]+#1/.Rule@@@dexStartConfocal),#6[[2]]}\[Transpose]}&@@{#[[1,1,1]],#[[1,1,2]],#[[1,1,4]],#[[1,1,-2]],#[[2]],#[[1,3;;-3,{9,7}]]\[Transpose]}&/@longTimesfig1;


(* ::Subsection:: *)
(*Separated confocal dataset*)


(* collects the data into trajectories and keep all points except last two points. Remove first two points for cells at the beginning of measurement, and three points for cells after symmetric division. 
Keep only trajecotries that are at least 11 time-points long. Recalculate the h2b normalization *)
gatheredDataConfocalSep= Table[Join[#[[;;6]],{#[[5]]/#[[6]]},#[[8;;]]]&/@If[StringLength@d[[1,4]]> 5,d[[4;;-3]],d[[3;;-3]] ],{d,Select[GatherBy[BackSubtractedConfocalDex,#[[;;4]]&],Length@#>= 8+3&]}];

(*organize into cells table. the data appears in smoothed and unsmoothed versions*)
organizedDataConfocalSep=Parallelize[MapIndexed[{#1,#2,#3,#4,#5,{(#6[[1]]+#1/.Rule@@@dexStartConfocal)[[;;]],smooth[#6[[2]],7],smooth[#6[[3]],7]}\[Transpose],{(#6[[1]]+#1/.Rule@@@dexStartConfocal),#6[[2]],#6[[3]]}\[Transpose]}&@@{#1[[1,1]],#1[[1,2]],#1[[1,4]],#1[[1,-2]],#2[[1]],#1[[;;,{9,7,15}]]\[Transpose]}&,gatheredDataConfocalSep]];

(*rule for type of divisions "N" -> not divided, "S" -> symmetrically divided and "A"-> asymmetric divisions *)
asymConfocalSep = Flatten/@({#[[{1,2,4}]],Which[#[[7]]!= "NA","N",#[[6]]!= "NA", "S", #[[5]]!= "NA","A"]}&/@GatherBy[DeleteDuplicates@ConfocalDex[[;;,{1,2,3,4,-5,-4,-3}]],#[[;;4]]&][[;;,1]])/.Null-> "N";

(* remove any trajectory with information about what happens after asymmetric division  *)
dataCSep= Select[organizedDataConfocalSep,StringPosition[#[[3]],"endo"|"cor"][[;;,1]]=={}&];

(* add cell fate explicitly into the data &  select cells from newest batch of experiments  *)
dConfSep=Select[Flatten[#,1]&/@({#,#[[;;3]]/.Thread[{#1,#2,#3}&@@@asymConfocalSep[[;;,;;3]]-> asymConfocalSep[[;;,4]]],#[[;;3]]/.endroundrule}&/@dataCSep)/.{False->0,True->1},ToExpression@StringTake[#[[1]],-5]>9050&];

(* prepare the same construct as before but without cell 1 *)
dConfnc1Sep=Select[Flatten[#,1]&/@({#,#[[;;3]]/.Thread[{#1,#2,#3}&@@@asymConfocalSep[[;;,;;3]]-> asymConfocalSep[[;;,4]]],#[[;;3]]/.endroundrule}&/@Select[dataCSep,ToExpression@StringTake[#[[3]],{5}]>1&])/.{False->0,True->1},ToExpression@StringTake[#[[1]],-5]>9050&];


(* ::Text:: *)
(*Prepare rules for normalizing cell size *)


(*choose only cells from beginning of experiment and their daughters. Gather them by origin (mother and daughters), and for each group  we take the min and 0.9 quantile.     *)
ruleforNSnorm1={#[[1,1]],#[[1,2]],StringTake[#[[1,3]],5]}->N@( {Min[#],Quantile[#,0.9]}&@N@Flatten[#[[;;,6,;;,3]]])&/@GatherBy[Select[dConfnc1Sep,StringLength@#[[3]]<7&],#[[;;2]] && StringTake[#[[3]],5]&];
(*For each cell we take the min and 0.9 quantile.     *)
ruleforNSnorm2={#[[1,1]],#[[1,2]],#[[1,3]]}->N@( {Min[#],Quantile[#,0.9]}&@N@Flatten[#[[;;,6,;;,3]]])&/@GatherBy[dConfnc1Sep,#[[;;2]] && #[[3]]&];
(* for cells at the start of experiment we use rule 1 and for all the daugther cells we use rule 2 *)
ruleforNSnorm = Join[Select[ruleforNSnorm1,StringLength[#[[1,3]]]==5&],Select[ruleforNSnorm2,StringLength[#[[1,3]]] > 5&]];


(* ::Subsection:: *)
(*Data for pericycle and nuclear SHR comparison (fig 1g &h) *)


(*load data*)
per=Import[basePath <> perPath,"CSV" ];

(*prepare rule that links cell identity and round to levels of SHR*)
mrule=Dispatch[Thread[per[[;;,{1,2,3,4,5}]]-> per[[;;,9;;]]]];

(*select from confocal data only cells with pericycle SHR levels and merge *)
combper=Parallelize[Join[#[[1]],#[[2]]]&/@Select[({ConfocalDex,DeleteCases[Evaluate[ConfocalDex[[;;,{1,2,3,4,8}]]/.mrule],"",\[Infinity]]}\[Transpose]),Length@#[[2]]==3&]];

(* collects the data into trajectories and change the last H2B values (1 or 2) to be like the average of the H2B values just before that and recalculate the h2b normalization*)
gatheredDataConfocalPer = Table[Flatten[If[#[[1,12]]== "NA" && Length@#>5,Join[#[[;;6]],{#[[5]]/#[[6]]},#[[8;;]]]&/@Join[#[[;;-3]],{Join[#[[-2,;;5]],{Mean[#[[-6;;-4,6]]]},#[[-2,7;;]]],Join[#[[-1,;;5]],{Mean[#[[-5;;-3,6]]]},#[[-1,7;;]]]}],If[#[[1,12]]== "NA" && Length@#>1,Join[#[[;;6]],{#[[5]]/#[[6]]},#[[8;;]]]&/@Join[#[[;;-2]],{Join[#[[-1,;;5]],{#[[-2,6]]},#[[-1,7;;]]]}],#]]&/@GatherBy[d,#[[4]]&],1],{d,GatherBy[combper,#[[1]]||#[[2]]||#[[3]]&]}];

(* change trajectory's cell position into last generation and filter out trajectories with less than 11 timepoints*)
daughtercellsConfocalper= Table[Select[gd,MemberQ[isnotContained@Flatten[DeleteDuplicates@gd[[;;,4]]],#[[4]]]&],{gd,compTraj/@gatheredDataConfocalPer}];
longTimesConfocalper= MapIndexed[{SortBy[#1,#[[8]]&],#2[[1]]}&,Select[GatherBy[Flatten[daughtercellsConfocalper,1],#[[1]]||#[[2]]||#[[4]]&],Length@#>10&]];

(*organize into cells table with the last 2 timepoints and first 2 timepoints removed. the data appears in smoothed and unsmoothed versions*)
organizedDataPerConfocal=Parallelize[{#1,#2,#3,#4,#5,{(#6[[1]]+#1/.Rule@@@dexStartConfocal)[[;;lptoberemoved]],smooth[#6[[2]],7],smooth[#6[[3]],7],smooth[#6[[4]],7],smooth[#6[[5]],7]}\[Transpose],{(#6[[1]]+#1/.Rule@@@dexStartConfocal),#6[[2]],#6[[3]],#6[[4]],#6[[5]]}\[Transpose]}&@@{#[[1,1,1]],#[[1,1,2]],#[[1,1,4]],#[[1,1,-4]],#[[2]],#[[1,(*3*);;-5,{9,5,15,16,17}]]\[Transpose]}&/@longTimesConfocalper];

(* remove any trajectory with information about what happens after asymmetric division  *)
dataC= Select[organizedDataPerConfocal,StringPosition[#[[3]],"endo"|"cor"][[;;,1]]=={}&];

(* add cell fate explicitly into the data &  select cells from newest batch of experiments  *)
dConfPer =Select[Flatten[#,1]&/@({#,#[[;;3]]/.Thread[{#1,#2,#3}&@@@asymConfocal[[;;,;;3]]-> asymConfocal[[;;,4]]],#[[;;3]]/.endroundrule}&/@dataC)/.{False->0,True->1},ToExpression@StringTake[#[[1]],-5]>9050&];

(* prepare the same construct as before but without cell 1 *)
dConfPernc1 =Select[Flatten[#,1]&/@({#,#[[;;3]]/.Thread[{#1,#2,#3}&@@@asymConfocal[[;;,;;3]]-> asymConfocal[[;;,4]]],#[[;;3]]/.endroundrule}&/@Select[dataC,ToExpression@StringTake[#[[3]],{5}]>1&])/.{False->0,True->1},ToExpression@StringTake[#[[1]],-5]>9050&];

(* collects the data into trajectories and keep all points except first 2 and last 2 points. Remove first three points for cells after symmetric division. 
Keep only trajecotries that are at least 11 time-points long. Recalculate the h2b normalization *)
gatheredDataConfocalPerSep= Table[Join[#[[;;6]],{#[[5]]/#[[6]]},#[[8;;]]]&/@If[StringLength@d[[1,4]]> 5,d[[4;;-5]],d[[(*3*);;-5]] ],{d,Select[GatherBy[combper,#[[;;4]]&],Length@#>= 8+3&]}];

(*organize into cells table. the data appears in smoothed and unsmoothed versions*)
organizedDataConfocalPerSep=Parallelize[MapIndexed[{#1,#2,#3,#4,#5,{(#6[[1]]+#1/.Rule@@@dexStartConfocal)[[;;]],smooth[#6[[2]],7],smooth[#6[[3]],7],smooth[#6[[4]],7],smooth[#6[[5]],7]}\[Transpose],{(#6[[1]]+#1/.Rule@@@dexStartConfocal),#6[[2]],#6[[3]],#6[[4]],#6[[5]]}\[Transpose]}&@@{#1[[1,1]],#1[[1,2]],#1[[1,4]],#1[[1,-4]],#2[[1]],#1[[;;,{9,5,15,16,17}]]\[Transpose]}&,gatheredDataConfocalPerSep]];

(* remove any trajectory with information about what happens after asymmetric division  *)
dataCPerSep= Select[organizedDataConfocalPerSep,StringPosition[#[[3]],"endo"|"cor"][[;;,1]]=={}&];

(* add cell fate explicitly into the data &  select cells from newest batch of experiments  *)
dConfPerSep=Select[Flatten[#,1]&/@({#,#[[;;3]]/.Thread[{#1,#2,#3}&@@@asymConfocalSep[[;;,;;3]]-> asymConfocalSep[[;;,4]]],#[[;;3]]/.endroundrule}&/@dataCPerSep)/.{False->0,True->1},ToExpression@StringTake[#[[1]],-5]>9050&];

(* prepare the same construct as before but without cell 1 *)
dConfnc1PerSep=Select[Flatten[#,1]&/@({#,#[[;;3]]/.Thread[{#1,#2,#3}&@@@asymConfocalSep[[;;,;;3]]-> asymConfocalSep[[;;,4]]],#[[;;3]]/.endroundrule}&/@Select[dataCPerSep,ToExpression@StringTake[#[[3]],{5}]>1&])/.{False->0,True->1},ToExpression@StringTake[#[[1]],-5]>9050&];

(* take only cells up to the first division *)
dConfnc1PerSepFirstGen = Select[dConfnc1PerSep,StringLength[#[[3]]]==5&];


(* ::Subsection:: *)
(*Hydroxy-urea data*)


(* ::Text:: *)
(*Load, separate titles and data, delete empty lines and remove cell1. *)


hydroRawData = Import[basePath <> hydroFileName, "XLSX"][[1]] /.""->"NA"; 
{hydroRawDataTitles, hydroRawData} = {hydroRawData[[1,;;7]], hydroRawData[[2 ;; ,;;7]]};
hydroRawData = DeleteCases[hydroRawData,{""..}];
hydroDatanc1=Select[hydroRawData,ToExpression@StringTake[#[[4]],{5}]>1&];


(* ::Text:: *)
(*prepare data for division diagram (figure 3j - bottom)*)


predivhydCon={#1,#3,StringTake[#4,5],#4,#2,{Red,#5},{Blue,#6},{Cyan,#7}}&@@@hydroDatanc1;
divOc=DeleteCases[predivhydCon,{_,"NA"},\[Infinity]][[;;,{1,2,3,4,5,6}]];


(* ::Subsection:: *)
(*LightSheet data*)


(*load data*)
LSRawData =Import[basePath <> LSFileName,"CSV"];

(*LSRawData = DeleteCases[LSRawData,{_,"Exp00455",__}];*)
(*load dex conentrations*)
colors =Import[basePath <> cFileName,"csv"];

(* separate titles from data*)
{LSRawDataTitles,LSRawData}  = {LSRawData[[1]],LSRawData[[2;;]]};

(* Load and extract the time where dex was introduced with respect to experiment inital measurements*)
{colorsTitles,colors} = {colors[[1]],colors[[2;;]]};

dexStartLS = Quiet@DeleteCases[{#1,QuantityMagnitude@UnitConvert[#,"h"]&@(DateObject[ToExpression/@StringSplit[#2,_?(#=="-"||#=="_"&)]]-DateObject[ToExpression/@StringSplit[#3,_?(#=="-"||#=="_"&)]])}&@@@
			(Map[ StringReplace[ToString@#," *"-> ""]&,colors[[;;,{1,5,6}]],{2}])
			,{___,DateObject[{-1},"Year","Gregorian",-4.`],___}];

rule = "Exp00"<>ToString@#[[1]]->#[[2]]&/@colors[[;;,{1,3}]];
 
 (* change experiment names to lowercase *)
tbrule = Select[#-> ToLowerCase@#&/@DeleteDuplicates@LSRawData[[;;,2]],#[[1]]!= #[[2]]&];

(* filter table to relevant columns *)
LSDex = Append[#,#[[1]]/.rule]&/@LSRawData[[;;,{2,3,4,5,7,9,11,8,12,10,13,14,15,16,6,17}]]/.tbrule;
LSDex =DeleteCases[ {#1,#2,#3,#4,#5,#6,#7,#8,#9,#10,#11+0.25,#12,#13,#14,#15,#16,#17}&@@@LSDex,{_,_,(*"cell1"|*)"cell6"|"cell7"|"cell8"|"cell9",__}];
LSDexTitles = Append[LSRawDataTitles[[{2,3,4,5,7,9,11,8,12,10,13,14,15,16,6,17}]],"Dex"]; 

(*background subtraction of SHR-GFP using mean of first three points*)
backGreenRuleLS= #[[1,{1,2,3}]]-> Mean[#[[;;3,5]]]&/@(SortBy[#,#[[10]]&]&/@GatherBy[LSDex,#[[1]]||#[[2]]||#[[3]]&]);

(*background and bleedtrhough correction *)
InterimBackSubtractedLSDex= {Max[#[[1]],0],Max[#[[2]],0],Max[#[[1]]/#[[2]],0],Max[#[[3]],0],Max[#[[3]]/#[[2]],0]}&@((IdentityMatrix[3]-BTEstimatesLS) . ({#5,#6,#8}-{({#1,#2,#3}/.backGreenRuleLS),AutoFluorescenceEstimatesLS[[2]],AutoFluorescenceEstimatesLS[[3]]}))&@@@LSDex;
BackSubtractedLSDex=Join[#[[1,;;4]],#[[2]],#[[1,10;;]]]&/@({LSDex,InterimBackSubtractedLSDex}\[Transpose]);

(* collects the data into trajectories and change the last H2B values (1 or 2) to be like the average of the SHR values just before that and recalculate the h2b normalization*)
gatheredDataLS = Table[Flatten[
						If[#[[1,14]]== "NA" && Length@#>5,
							Join[#[[;;6]],{#[[5]]/#[[6]],#[[8]],#[[8]]/#[[6]]},#[[10;;]]]&/@
								Join[#[[;;-3]],{Join[#[[-2,;;5]],{Mean[#[[-6;;-4,6]]]},#[[-2,7;;]]],Join[#[[-1,;;5]],{Mean[#[[-5;;-3,6]]]},#[[-1,7;;]]]}],
							If[#[[1,14]]== "NA" && Length@#>1,
								Join[#[[;;6]],{#[[5]]/#[[6]],#[[8]],#[[8]]/#[[6]]},#[[10;;]]]&/@
									Join[#[[;;-2]],{Join[#[[-1,;;5]],{#[[-2,6]]},#[[-1,7;;]]]}],
								#]]&/@GatherBy[d,#[[4]]&],1],{d,GatherBy[BackSubtractedLSDex,#[[1]]||#[[2]]||#[[3]]&][[;;]]}];

(* change trajectory's cell position into last generation and filter out trajectories with less than 11 timepoints*)
daughtercellsLS=
Table[
Select[gd,MemberQ[isnotContained@Flatten[DeleteDuplicates@gd[[;;,4]]],#[[4]]]&],{gd,compTraj/@gatheredDataLS}]; 
longTimesLS = MapIndexed[{SortBy[#1,#[[10]]&],#2[[1]]}&,Select[GatherBy[Flatten[daughtercellsLS,1],#[[1]]||#[[2]]||#[[4]]&],Length@#>10&]]; 

(*organize into cells table. the data appears in smoothed and unsmoothed versions and remove last 4 points*)
organizedDataLS=Parallelize[Quiet@{#1,#2,#3,#4,#5,{(#6[[1]]+((StringTake[#1,-3])/.Rule@@@dexStartLS))[[;;lptoberemoved]],smooth[#6[[2]],7],smooth[#6[[3]],7],smooth[#6[[4]],7]}\[Transpose],{(#6[[1]]+((StringTake[#1,-3])/.Rule@@@dexStartLS)),#6[[2]],#6[[3]],#6[[4]]}\[Transpose]}&@@{#[[1,1,1]],#[[1,1,2]],#[[1,1,4]],#[[1,1,-1]],#[[2]],#[[1,;;-5,{11,7,9,16}]]\[Transpose]}&/@longTimesLS];

(*rule for asymmetric divisions has cell identifiers and a flag that is true if cells ends up dividing asymmetrically *)
asymLS = Flatten/@({#[[1,{1,2,4}]],Or@@(#!= "NA"&/@#[[;;,5]])}&/@GatherBy[DeleteDuplicates@LSDex[[;;,{1,2,3,4,12,13,14}]],#[[1]]&&#[[2]]&&#[[4]]&]);

(* rule for end of time course for each cell*)
endroundruleLS = DeleteDuplicates[({#[[1]],#[[2]],#[[4]]}-> #[[12]]/4+(StringTake[#[[1]],-3]/.Rule@@@dexStartLS))&/@LSDex]/._+("NA")/4->"NA";

(* add cell fate explicitly into the data &  select cells from newest batch of experiments  *)
dLS =Flatten[#,1]&/@({#,#[[;;3]]/.Thread[{#1,#2,#3}&@@@asymLS[[;;,;;3]]-> asymLS[[;;,4]]],#[[;;3]]/.endroundruleLS}&/@organizedDataLS)/.{False->0,True->1};

(* prepare the same construct as before but without cell 1 *)
dLSnc1 = Flatten[#,1]&/@({#,#[[;;3]]/.Thread[{#1,#2,#3}&@@@asymLS[[;;,;;3]]-> asymLS[[;;,4]]],#[[;;3]]/.endroundruleLS}&/@Select[organizedDataLS,ToExpression@StringTake[#[[3]],{5}]>1&])/.{False->0,True->1};

(* prepare list of symmetric divisions *)
symsLS = Table[Block[{rd =Cases[LSDex,{cell[[1]],cell[[2]],__}],ns=StringDrop[cell[[3]],-1] ,times={}},{While[StringLength[ns]>4,AppendTo[times,SortBy[Select[rd,#[[4]]==ns&],#[[11]]&][[-1,11]]];ns=StringDrop[ns,-1]]};cell[[;;3]]->Reverse@times + (((StringTake[cell[[1]],-3])/.Rule@@@dexStartLS))],{cell,Select[dLS,StringLength[#[[3]]]>5&][[;;]]}];


(* ::Subsection:: *)
(*Separated LS dataset*)


(* collects the data into trajectories and keep all points except last four points. Remove first three points for cells after symmetric division. 
Keep only trajecotries that are at least 11 time-points long. Recalculate the h2b normalization *)
gatheredDataLSSep= Table[Join[#[[;;6]],{#[[5]]/#[[6]]},#[[8;;]]]&/@If[StringLength@d[[1,4]]> 5,d[[4;;-5]],d[[;;-5]] ],{d,Select[GatherBy[BackSubtractedLSDex,#[[;;4]]&],Length@#>= 8+3&]}];

(*organize into cells table. the data appears in smoothed and unsmoothed versions*)
organizedDataLSSep=Parallelize[MapIndexed[{#1,#2,#3,#4,#5,{(#6[[1]]+((StringTake[#1,-3])/.Rule@@@dexStartLS))[[;;lptoberemoved]],smooth[#6[[2]],7],smooth[#6[[3]],7],smooth[#6[[4]],7]}\[Transpose],{(#6[[1]]+((StringTake[#1,-3])/.Rule@@@dexStartLS)),#6[[2]],#6[[3]],#6[[4]]}\[Transpose]}&@@{#1[[1,1]],#1[[1,2]],#1[[1,4]],#1[[1,-1]],#2[[1]],#1[[;;,{11,7,9,16}]]\[Transpose]}&,gatheredDataLSSep]];

(*rule for type of divisions "N" -> not divided, "S" -> symmetrically divided and "A"-> asymmetric divisions *)
asymLSSep = Flatten/@({#[[{1,2,4}]],Which[#[[7]]!= "NA","N",#[[6]]!= "NA", "S", #[[5]]!= "NA","A"]}&/@GatherBy[DeleteDuplicates@LSDex[[;;,{1,2,3,4,-6,-5,-4}]],#[[;;4]]&][[;;,1]])/.Null-> "N";

(* remove any trajectory with information about what happens after asymmetric division  *)
dLSSep =Flatten[#,1]&/@({#,#[[;;3]]/.Thread[{#1,#2,#3}&@@@asymLSSep[[;;,;;3]]-> asymLSSep[[;;,4]]],#[[;;3]]/.endroundruleLS}&/@organizedDataLSSep);

(* add cell fate explicitly into the data &  select cells from newest batch of experiments  *)
dLSnc1Sep=Flatten[#,1]&/@({#,#[[;;3]]/.Thread[{#1,#2,#3}&@@@asymLSSep[[;;,;;3]]-> asymLSSep[[;;,4]]],#[[;;3]]/.endroundruleLS}&/@Select[organizedDataLSSep,ToExpression@StringTake[#[[3]],{5}]>1&])/.{False->0,True->1};

(* Remove trajectories of length 1 from the separated LS dataset*)
dLSnc1Sep = Delete[dLSnc1Sep,Position[Length[DeleteDuplicates[#]]&/@dLSnc1Sep[[;;,6,;;,4]],1]];


(* ::Text:: *)
(*Prepare rules for normalizing cell size *)


(*choose only cells from beginning of experiment and their daughters. Gather them by origin (mother and daughters), and for each group  we take the min and 0.9 quantile.     *)
ruleforlsnorm1={#[[1,1]],#[[1,2]],StringTake[#[[1,3]],5]}->N@( {Min[#],Quantile[#,0.9]}&@N@Flatten[#[[;;,6,;;,4]]])&/@GatherBy[Select[dLSnc1Sep,StringLength@#[[3]]<7&],#[[;;2]] && StringTake[#[[3]],5]&];
(*For each cell we take the min and 0.9 quantile.     *)
ruleforlsnorm2={#[[1,1]],#[[1,2]],#[[1,3]]}->N@( {Min[#],Quantile[#,0.9]}&@N@Flatten[#[[;;,6,;;,4]]])&/@GatherBy[dLSnc1Sep,#[[;;2]] && #[[3]]&];
(* for cells at the start of experiment we use rule 1 and for all the daugther cells we use rule 2 *)
ruleforlsnorm = Join[Select[ruleforlsnorm1,StringLength[#[[1,3]]]==5&],Select[ruleforlsnorm2,StringLength[#[[1,3]]] > 5&]];


(* ::Subsection:: *)
(*Data for quantification of figs3a-c *)


(*Load and reorder data  and separate titles from data*)
exp160sraw = Import[basePath <> exp160FileName, "CSV"];
exp160raw = Join[#[[;;5]],{0},{#[[6]]},#[[9;;10]],{#[[12]]},{#[[8]]},{#[[11]]},#[[13;;]]]&/@exp160sraw;
{exp160rawTitles, exp160rawData} = {exp160raw[[1]], exp160raw[[2 ;; ]]};

(* change experiment names to lowercase *)
tbruleexp160 = Select[#-> ToLowerCase@#&/@DeleteDuplicates@exp160rawData[[;;,2]],#[[1]]!= #[[2]]&];

(* filter table to relevant columns and rows *)
exp160Dex = exp160rawData[[;;,{2,3,4,5,7,9,11,8,12,10,13,14,15,16,6}]];
exp160Dex =DeleteCases[ {#1,#2,#3,#4,#5,#6,#7,#8,#9,#10,#11+0.25,#12,#13,#14,#15}&@@@exp160Dex,{_,_,"cell6"|"cell7"|"cell8"|"cell9",__}];
exp160DexTitles = Append[exp160rawTitles[[{2,3,4,5,7,9,11,8,12,10,13,14,15,16,6,17}]],"Dex"]; 

(*background subtraction of SHR-GFP using mean of first three points*)
backGreenRuleexp160= #[[1,{1,2,3}]]-> Mean@#[[;;3,5]]&/@(SortBy[#,#[[10]]&]&/@GatherBy[exp160Dex,#[[1]]||#[[2]]||#[[3]]&]);

(*background and bleedtrhough correction *)
InterimbackGreenRuleexp160= {Max[#[[1]],0],Max[#[[2]],0],Max[#[[1]]/#[[2]],0],Max[#[[3]],0],Max[#[[3]]/#[[2]],0]}&@((IdentityMatrix[3]-BTEstimatesLS) . ({#5,#6,#8}-{({#1,#2,#3}/.backGreenRuleexp160),AutoFluorescenceEstimatesLS[[2]],AutoFluorescenceEstimatesLS[[3]]}))&@@@exp160Dex;
BackSubtractedexp160Dex=Join[#[[1,;;4]],#[[2]],#[[1,10;;]]]&/@({exp160Dex,InterimbackGreenRuleexp160}\[Transpose]);

(* collects the data into trajectories and change the last H2B values (1 or 2) to be like the average of the SHR values just before that and recalculate the h2b normalization*)
gatheredDataexp160 = Table[Flatten[If[#[[1,14]]== "NA" && Length@#>5,Join[#[[;;6]],{#[[5]]/#[[6]],#[[8]],#[[8]]/#[[6]]},#[[10;;]]]&/@Join[#[[;;-3]],{Join[#[[-2,;;5]],{Mean[#[[-6;;-4,6]]]},#[[-2,7;;]]],Join[#[[-1,;;5]],{Mean[#[[-5;;-3,6]]]},#[[-1,7;;]]]}],If[#[[1,14]]== "NA" && Length@#>1,Join[#[[;;6]],{#[[5]]/#[[6]],#[[8]],#[[8]]/#[[6]]},#[[10;;]]]&/@Join[#[[;;-2]],{Join[#[[-1,;;5]],{#[[-2,6]]},#[[-1,7;;]]]}],#]]&/@GatherBy[d,#[[4]]&],1],{d,GatherBy[BackSubtractedexp160Dex,#[[1]]||#[[2]]||#[[3]]&][[;;]]}];

(* change trajectory's cell position into last generation and filter out trajectories with less than 11 timepoints*)
daughtercellsexp160=
Table[
Select[gd,MemberQ[isnotContained@Flatten[DeleteDuplicates@gd[[;;,4]]],#[[4]]]&],{gd,compTraj/@gatheredDataexp160}]; 
longTimesexp160 = MapIndexed[{SortBy[#1,#[[10]]&],#2[[1]]}&,Select[GatherBy[Flatten[daughtercellsexp160,1],#[[1]]||#[[2]]||#[[4]]&],Length@#>10&]]; 

(*organize into cells table. the data appears in smoothed and unsmoothed versions and remove last 4 points*)
organizedDataexp160=Parallelize[Quiet@{#1,#2,#3,#4,#5,{#6[[1]][[;;lptoberemoved]],smooth[#6[[2]],7],smooth[#6[[3]],7]}\[Transpose],{(#6[[1]]),#6[[2]],#6[[3]]}\[Transpose]}&@@{#[[1,1,1]],#[[1,1,2]],#[[1,1,4]],#[[1,1,-1]],#[[2]],#[[1,;;-5,{11,7,9}]]\[Transpose]}&/@longTimesexp160];

(*rule for asymmetric divisions has cell identifiers and a flag that is true if cells ends up dividing asymmetrically *)
asymexp160 = Flatten/@({#[[1,{1,2,4}]],Or@@(#!= "NA"&/@#[[;;,5]])}&/@GatherBy[DeleteDuplicates@exp160Dex[[;;,{1,2,3,4,12,13,14}]],#[[1]]&&#[[2]]&&#[[4]]&]);

(* rule for end of time course for each cell*)
endroundruleexp160 = DeleteDuplicates[({#[[1]],#[[2]],#[[4]]}-> #[[12]]/4)&/@exp160Dex]/._+("NA")/4->"NA";

(* add cell fate explicitly into the data &  select cells from newest batch of experiments  *)
dexp160 =Flatten[#,1]&/@({#,#[[;;3]]/.Thread[{#1,#2,#3}&@@@asymexp160[[;;,;;3]]-> asymexp160[[;;,4]]],#[[;;3]]/.endroundruleexp160}&/@organizedDataexp160)/.{False->0,True->1};


(* ::Section:: *)
(*Alignment and feature analysis*)


(* ::Subsection:: *)
(*Prepare for feature extraction*)


(* ::Text:: *)
(*Move from data formatted as d into wn constructs, in the wn format the non-normalized and the normalized versions of the data appears both of them smoothed and consolidate data into single variable.  *)


wnLSSep  = d2wn[dLSnc1Sep,ruleforlsnorm,False];
wnLS  = d2wn[dLSnc1,ruleforlsnorm,True];
wnConfSep  = d2wn[dConfnc1Sep,ruleforNSnorm,False];
wnConf  = d2wn[dConfnc1,ruleforNSnorm,True];
wnDat = {wnConf,wnConfSep, wnLS,wnLSSep};


(* ::Text:: *)
(*align data: the separated trajectories are aligned to their beginnings so that they all start at 0. The other datasets are aligned to their inflection points (see methods)*)
(*group under one variable*)


alignedConf = aligningByRise[wnConf,14,0];
alignedLS = aligningByRise[wnLS,38,4.75];
alignedConfSep = aligningTostart[wnConfSep];
alignedLSSep = aligningTostart[wnLSSep];
aligneddat = {alignedConf,alignedConfSep, alignedLS,alignedLSSep};


(* ::Text:: *)
(*calculate best threshold for separating data either for full trajectory and for separated - get the best threshold for first nuclear size window*)


(* LS SHR*)
{thHLS,x} =  bestThresholds[wnLS,1,2,rangeLS,False];
(* Confocal SHR*)
{thHConf,x} =bestThresholds[wnConf,1,2,rangeConf,False];
(* LS SCR*)
{thCLS,x} =  bestThresholds[wnLS,1,3,rangeLS,False];
(* LS separated SHR*)
{thLSSep,x} =  bestThresholds[wnLSSep,4,2,rangeLS,True];
(* LS separated SCR*)
{thLSSepSCR,x} =  bestThresholds[wnLSSep,4,3,rangeLS,True];
(* Confocal separated SHR*)
{thConfSep,x} =  bestThresholds[wnConfSep,4,2,rangeConf,True];
(* take first nuclear size window*)
thHLS = thHLS[[1]];
thHConf = thHConf[[1]];
thCLS = thCLS[[1]];
thLSSep=thLSSep[[1]];
thLSSepSCR=thLSSepSCR[[1]];
thConfSep=thConfSep[[1]];


(* ::Subsection:: *)
(*Extract features and save them under the same variable (ts for titles and features for the features)*)


{TLS,featuresLS     }  = calcFeatures[alignedLS, False, True,thHLS,thCLS,LSThr];
{TLSsep,featuresLSSep  } = calcFeatures[alignedLSSep, True, True,thLSSep,thLSSepSCR,LSThr];
{TCon,featuresConf   } = calcFeatures[alignedConf, False, False,thHConf,thCLS,ConfThr];
{TConSep,featuresConfSep} = calcFeatures[alignedConfSep, True, False,thConfSep,thCLS,ConfThr];

ts = {TCon, TConSep, TLS, TLSsep};
features  = {featuresConf, featuresConfSep, featuresLS, featuresLSSep};


(* ::Subsection:: *)
(*Feature analysis and SVM *)


(* ::Text:: *)
(*Prepare training and testing sets*)


sets = Table[
SeedRandom[5434219];
trpca=RandomSample[Range[1,f],Round[0.6*f]];
tepca= Complement[Range[1,f],trpca];
{trpca,tepca}
,{f,Length/@features}];


(* ::Text:: *)
(*calculate the prediction score of the best threshold for each feature *)


(* run over all datasets*)
onebyoneScores = Parallelize@Table[Block[{feats},
					(* over all fetures (the first 8 are cell identifiers)*)
					feats=Range[8,Length@f[[1,1]]];
					Block[{trset,teset,dd,rf,clas,ba,res},
					Table[
						(*prepare training and testing rules*) 
						trset =Thread[f[[1]][[f[[2,1]],rf]]->f[[1]][[f[[2,1]],6]]];
						teset =Thread[f[[1]][[f[[2,2]],rf]]->f[[1]][[f[[2,2]],6]]];
						(* find brst threshold*) 
						clas=class4oneFeature[trset];
						(*index,feature name, accuracy on test set, threshld, feature direction *) 
						{rf,f[[3,rf]],accuracy[teset,clas],clas[[1]],clas[[2]]}
						,{rf,feats}]]]
						,{f,({features,sets,ts}\[Transpose])}];


(* ::Text:: *)
(*for each non-separated dataset we remove feature that are highly correlated with the length of the trajectory - as this is a technical cue that let's the ML algorithm to easily identify non-divided cells. *)


corLengths  = Table[Block[{ll=Length/@f[[2,;;,6]]},{#,f[[3,#]]}&/@Select[(Abs@Table[{s,Correlation[f[[1,;;,s]],ll]},{s,8,f[[1,1]]//Length}]),#[[2]]>= .7&][[;;,1]]],{f,({features,aligneddat,ts}\[Transpose])}];

remCors = Table[Delete[#,({#}&/@f[[2,;;,1]])]&/@f[[1]],{f,({features,corLengths}\[Transpose])}];
remCorsT= Table[Delete[f[[1]],({#}&/@f[[2,;;,1]])],{f,({ts,corLengths}\[Transpose])}];

features2= {remCors[[1]],features[[2]],remCors[[3]],features[[4]]};
ts2= {remCorsT[[1]],ts[[2]],remCorsT[[3]],ts[[4]]};


(* ::Text:: *)
(*Calculate feature p-values using Mann-Whitney test for all features for all datasets*)


pvals=Table[MannWhitneyTest[#](*}*)&@(GatherBy[f[[;;,{6,#}]],First][[;;,;;,2]])&/@Range[8,Length@f[[1]]],{f,features}];


(* ::Text:: *)
(*Calculate FDR using the BH method*)


FDRs=Table[fdr=pval;(fdr[[#[[;;,1]]]] =#[[;;,2]] )&@MapIndexed[{#1, ( pval[[#1]] Length@pval)/#2[[1]]}&,Ordering[pval]];fdr,{pval,pvals}];


(* ::Text:: *)
(*Calculate and export a table with the SVM score for each dataset*)


Export[finalFiguresFolder<> "MLResults "<>DateString[]<>".csv",Prepend[(Prepend[Parallelize@Table[ 
	Table[Block[{trset,teset,dd,rf,clas,ba,res,feats=s},
			trset =Thread[f[[1]][[f[[2,1]],feats]]->f[[1]][[f[[2,1]],6]]];
			teset =Thread[f[[1]][[f[[2,2]],feats]]->f[[1]][[f[[2,2]],6]]];
			clas=Classify[trset,ClassPriors-><|0.->0.5,1.->0.5|>,Method->"SupportVectorMachine",TrainingProgressReporting->None,RandomSeeding->054562];
			ClassifierMeasurements[clas,teset,"Accuracy"]
			]
,{s,Prepend[f[[3;;]],Range[8,Length@f[[1,1]]]]}],

{f,{features,sets}\[Transpose]}],{"All"}]\[Transpose]),{"","Conf","ConfSep","LS","LSSep"}]\[Transpose]];


(* ::Section:: *)
(*Threshold plots*)


(* ::Subsection:: *)
(*Calculate and export threshold plot and associated data*)


(* ::Text:: *)
(*LS, LSsep and  SHR and SCR*)


windowsLS = Parallelize@Table[
	Block[{w,wsep},
		w=  plotThresholdCurves[wnLS,4,difs[[1]],rangeLS,ruleforlsnorm,subpaths[[3]],difs[[2]],False];
		wsep = plotThresholdCurves[wnLSSep,4,difs[[1]],rangeLS,ruleforlsnorm,subpaths[[4]],difs[[2]],True];
		{w,wsep}]	
	,{difs,{{2,"SHR_"},{3,"SCR_"}}}];


(* ::Text:: *)
(*Conf and Confsep*)


WindowsCon = {plotThresholdCurves[wnConf,4,2,rangeConf,ruleforNSnorm,subpaths[[1]],"",False],
			  plotThresholdCurves[wnConfSep,4,2,rangeConf,ruleforNSnorm,subpaths[[2]],"",True]};
	


(* ::Text:: *)
(*prepare data for fig 3f*)


dataforBars=Prepend[Map[Max,Transpose/@windowsLS[[;;,2,1]],{2}],Max/@(WindowsCon[[2,1]]\[Transpose])];


(* ::Section:: *)
(*Model Fitting*)


(* ::Subsection:: *)
(*prepare data - end up with one trajectory for SHR and one for SCR*)


(* ::Text:: *)
(*Take the mean of each root - align the roots normalize them and the the mean for all roots.*)


(* take the mean of each root we take deata points with at least 11 root in them*)
rootdat=(Join[#[[1,{1,2,3,4,5}]],{Mean/@TakeWhile[(GatherBy[Flatten[#[[;;,6]],1],#[[1]]&]),Length@#>10&]},{Mean/@TakeWhile[(GatherBy[Flatten[#[[;;,7]],1],#[[1]]&]),Length@#>10&]},#[[1,{8}]]]&/@GatherBy[wnLS,#[[1]]&&#[[8]]&]);
rootdatwork = Select[rootdat,Length@#[[6]]>0&];
(* select only fully induced roots*)
forAlignments = SortBy[Select[rootdatwork,#[[4]]==40.&],#[[1]]&];
(* Following code is a small GUI for alignment of the roots*)
(*Manipulate[Show[Table[ListLinePlot[Legended[{#1-{t1,t2,t3,t4,t5,t6,t7,t8,t9}[[i]],#2}&@@@#[[7,;;,;;2]],#[[1]]],PlotStyle->cd[i],PlotRange->All]&@forAlignments[[i]],{i,9}],PlotRange->All],##]&@@Evaluate[{{#,0},-7,7}&/@ToExpression["t" <> ToString@#&/@Range@9]]*)
(*{t1,t2,t3,t4,t5,t6,t7,t8,t9}->{4.48,0.4,0.66,3.5,0.14,4.7,0.08,2.26,0.18}*)

(* aligned data*)
aligned=Join[#[[1,;;5]],{Block[{t=#[[2]]},{#1-t,#2,#3,#4}&@@@#[[1,7]]]}]&/@({forAlignments,{4.48,0.4,0.66,3.5,0.14,4.7,0.08,2.26,0.18}}\[Transpose]);
(* Take mean of data*)
dataLSdata=Select[{#[[;;,1]],normalize@#[[;;,2]],normalize@#[[;;,3]]}\[Transpose]&@(Mean/@TakeWhile[(GatherBy[Flatten[#[[;;,6]],1],Round[#[[1]],0.25]&])&@aligned,Length@#>1&]),#[[1]]>=0&];

(* Add some dummy fields at beginning of data to fit in the fitting function*)
dff=Join[ConstantArray[1,5],{dataLSdata}];


(* ::Subsection:: *)
(*prepare models for fitting*)


(*make sure some variables are cleared*)
c=.;
A=.;
x=.;

(*define equation models*)
hillModel           = {Derivative[1][c][t] == (-\[Alpha])*c[t] + kProd*(s[t]^h/(kh^h + s[t]^h))};
hillFsimplerModel   = {Derivative[1][c][t] == kBasal - \[Alpha]*c[t] + (kProd*tot*s[t]*c[t])/(1 + A*s[t] + B*c[t] + tot*s[t]*c[t])};
MMModel             = {Derivative[1][c][t]==-\[Alpha] c[t]+(kProd s[t])/(kh+s[t])};

(*define model's parameters *)				 				 
hillParams           = {\[Alpha],kProd,kh,h};
hillFsimplerParams   = {\[Alpha],kProd,kBasal,A,B,tot};
MMParams             = {\[Alpha] ,kProd,kh};

(*define initial conditions for each model *)				 				 
hillIC               = {{c[t]== 0}};
hillFsimplerIC       = {{c[t]==kBasal/\[Alpha]}};
MMIC                 =  {{c[t]== 0}};

(*some constant paramteres for fitting function, c[t] in all is scr s[t] is shr and no scaling is needed (prefactor1/2 = 1) *)				 				 				  				  				  
vars = {c[t]} ;
ins = s[t];
prefactor = 1;
prefactor2 = 1;

(* name of models *)
modelnamesAll={"hill","Michaelis-Menten","FB"};


(* ::Subsection:: *)
(*Fitting the models *)


(* ::Text:: *)
(*Fitting results data file - If running fitting is too long - one can load this file with our results - change path if needed and load results.*)


{resHills,simpleFeedbackHills2,resMM} =<<  (basePath <> "fit_params_20220908b.fits");


(* ::Text:: *)
(*Running fitting for the Michaelis-Menten model - can take a couple hours to run thus commented out*)


(*resMM = gradODEs[MMModel,vars,ins,MMParams,dff,MMIC,0,0.01,{0.1,0.1,5},1,5000];*)


(* ::Text:: *)
(*Running fitting for the feedback model - can take a few hours to run*)


(*simpleFeedbackHills2 = {gradODEs[hillFsimplerModel,vars,ins,hillFsimplerParams,dff,hillFsimplerIC,0,0.01,{1,1,1,0.1,0.1,1},1,5000,1]};*)


(* ::Text:: *)
(*Running fitting for the hill models. It runs 10 times with different initial parameter guess for the hill coefficient (from 1 to 10)- can take a few hours to run *)


(*resHills = ParallelTable[gradODEs[hillModel,vars,ins,hillParams,dff,hillIC,0,0.01,{0.1,2,2,q},1,5000,1],{q,1,10}];*)


(* ::Text:: *)
(*Save Fitting results*)


(*{resHills,simpleFeedbackHills2,resMM} >> "fit_params_20220908b.fits";*)


(* ::Text:: *)
(*Get best fitted curve by solving ODE equations wit best parameters*)


Hillfit      =Quiet@Evaluate@NDSolveValue[Flatten[{hillModel/.Flatten[{Thread[hillParams->#],s[t]->Interpolation[dataLSdata[[;;,;;2]]][t]}],hillIC/.x_[t]->x[0.1]}],c[t],{t,0.1,35}]&/@resHills[[;;,3]];
sfeedhillfit =Quiet@Evaluate@NDSolveValue[Flatten[{hillFsimplerModel,hillFsimplerIC/.x_[t]->x[0.1]}/.Flatten[{Thread[hillFsimplerParams->#],s[t]->Interpolation[dataLSdata[[;;,;;2]]][t]}]],c[t],{t,0.1,35}]&/@simpleFeedbackHills2[[;;,3]];
MMfit        =Quiet@Evaluate@NDSolveValue[Flatten[{MMModel/.Flatten[{Thread[MMParams->resMM[[3]]],s[t]->Interpolation[dataLSdata[[;;,;;2]]][t]}],MMIC/.x_[t]->x[0.1]}],c[t],{t,0.1,35}];


(* ::Subsection:: *)
(*Obtain (\[Chi]^2) , R^2  and adjusted R^2 values for the fits*)


(* ::Text:: *)
(*We chose the best fit hill as the 6th (see methods)*)


(* ::Text:: *)
(*R^2*)


r2M = Quiet@Chop@Rsq[MMfit/.t->dataLSdata[[;;,1]],dataLSdata[[;;,3]]];
r2H = Quiet@Chop@Rsq[Hillfit[[6]]/.t->dataLSdata[[;;,1]],dataLSdata[[;;,3]]];
r2F = Quiet@Chop@Rsq[sfeedhillfit[[1]]/.t->dataLSdata[[;;,1]],dataLSdata[[;;,3]]];


(* ::Text:: *)
(*adjusted R^2*)


r2AM = Quiet@Chop@(1-(1-(Rsq[MMfit/.t->dataLSdata[[;;,1]],dataLSdata[[;;,3]]]))(Length[dataLSdata[[;;,3]]]-1)/(Length[dataLSdata[[;;,3]]]-Length[hillParams]-1));
r2AH = Quiet@Chop@(1-(1-(Rsq[Hillfit[[6]]/.t->dataLSdata[[;;,1]],dataLSdata[[;;,3]]]))(Length[dataLSdata[[;;,3]]]-1)/(Length[dataLSdata[[;;,3]]]-Length[hillParams]-1));
r2AF = Quiet@Chop@(1-(1-(Rsq[sfeedhillfit[[1]]/.t->dataLSdata[[;;,1]],dataLSdata[[;;,3]]]))(Length[dataLSdata[[;;,3]]]-1)/(Length[dataLSdata[[;;,3]]]-Length[hillFsimplerParams]-1));


(* ::Text:: *)
(*\[Chi]^2*)


chiM = Quiet@Chop@chisq[dataLSdata[[;;,3]],MMfit/.t->dataLSdata[[;;,1]]];
chiH = Quiet@Chop@chisq[dataLSdata[[;;,3]],Hillfit[[6]]/.t->dataLSdata[[;;,1]]];
chiF = Quiet@Chop@chisq[dataLSdata[[;;,3]],sfeedhillfit[[1]]/.t->dataLSdata[[;;,1]]];


(* ::Section:: *)
(*Relevant Figures for Paper*)


(* ::Subsection:: *)
(*Figure 1*)


(* ::Text:: *)
(*Figure 1c - prepare all confocal cell files column plots in a folder *)


(* create directory*)
dir1c = finalFiguresFolder <> "/fig1c-smoothed cell file" <>DateString[];
CreateDirectory[dir1c];
(*create plots*)
(*Parallel*)Table[Block[{synf1d = Reverse@SortBy[GatherBy[{StringTake[#[[3]],{5}],#[[-1]]/."A"-> Orange/."S"->Black/."N"-> White, #[[6]]}&@Join[#[[;;5]],#[[6,-1]],{#[[8]]}]&/@Cases[dConfSep,{cf[[1]],cf[[2]],__}],First],#[[1,1]]&],
cfile = (Reverse@SortBy[#,ToExpression@StringTake[#[[1,3]],{5}]&]&@GatherBy[Cases[dConf,{cf[[1]],cf[[2]],__}],
		ToExpression@StringTake[#[[3]],{5}]&]),maxx,maxy},
maxx= Flatten[cfile[[;;,;;,6,;;,1]]]//Max;
maxy= Flatten[cfile[[;;,;;,6,;;,2]]]//Max;
Export[dir1c<>"/"<>StringRiffle@cf<>".pdf",#]&@GraphicsColumn[#,ImageSize->defaultImageSize,Spacings->{0,0},ImagePadding->10]&@
	Table[Show[{ListLinePlot[q[[1,;;,6,;;,;;2]],##]&@@q[[2]],
				ListLinePlot[q[[1,;;,7,;;,;;2]],##,ImageSize->defaultImageSize]&@@q[[3]]},PlotRange->{{0,maxx},{0,1.05maxy}},Epilog-> Evaluate[{Dashed,Thick,#[[2]],Line[{{#[[3]],0},{#[[3]],1}}]}&/@q[[4]]]],{q,{cfile,style1d,style1db,synf1d}\[Transpose]}]],{cf,DeleteDuplicates[dConfSep[[;;,;;2]]]}];


(* ::Text:: *)
(*Figure 1d - quantification of cell on left*)


Export[finalFiguresFolder<>"fig1d_"<>#[[2]]<>".pdf",ListLinePlot[#[[1]],PlotRange->All]]&/@({{{#1,#2}\[Transpose],{#1,#3}\[Transpose],{#1,#4}\[Transpose]}&@@(Cases[ConfocalDex,{"Exp09065","upperleft","cell2",__}][[;;,{9,5,6,7}]]\[Transpose]),{"SHR","H2B","SHR_over_H2B"}}\[Transpose]);


(* ::Text:: *)
(*Figure 1f - peak less than 0.08 *)


Export[finalFiguresFolder<>"fig1f-maxExpression_5peaks_not_sym_0p08_"<>DateString[]<>".pdf",#]&@
Show[{ListLinePlot[SortBy[Select[aligningToEnd[wnConf],#[[8]]==1&& Max@#[[6,;;,2]]<0.08&& StringLength[#[[3]]]==5&&(#[[6,-1,1]]-SortBy[#[[6]],#[[2]]&][[-1,1]] > 5 ) &],
#[[6,-1,1]]-SortBy[#[[6]],#[[2]]&][[-1,1]]&][[-5;;,6,;;,;;2]],AspectRatio->1,AxesOrigin->{-15,0},PlotRange-> All,##]&@@style1jO[[2]],
	ListLinePlot[aligningToEnd[Cases[organizedDatafig1,{"Exp09025","lowerleft","cell2",__}]][[1,6,;;,;;2]],PlotStyle->{{Opacity[.5],gg}}]},PlotRange->All];


(* ::Text:: *)
(*Figure 1g - cell files  of pericycle shr and nuclear shr*)


(* Make directory*)
perResPath2 = finalFiguresFolder <> "/fig1g-per " <> DateString[]<>"/";
CreateDirectory[perResPath2];
(*create plots*)
Identity[Table[
			Export[perResPath2 <> StringRiffle[cf,"_"]<>".pdf",
			GraphicsColumn[Table[ListLinePlot[{{r[[6,;;,1]],normalize@r[[6,;;,2]]}\[Transpose],{r[[6,;;,1]],normalize@r[[6,;;,5]]}\[Transpose]},
			PlotStyle->Evaluate[{Opacity[.5],#}&/@{gg,Black}],PlotRange->{{0,Max[Flatten@Cases[dConfnc1PerSepFirstGen[[;;]],{cf[[1]],cf[[2]],__}][[;;,6,;;,1]]]},{0,1.3}},Ticks->{Range[0,25,5],{0,0.25,0.5,0.75,1,1.25}},##]&@@style1per,{r,
				Reverse@SortBy[Cases[dConfnc1PerSepFirstGen[[;;]],{cf[[1]],cf[[2]],__}],#[[3]]&]}],Spacings->{0,-95},ImagePadding->20],ImageSize->defaultImageSize],
				{cf,DeleteDuplicates@dConfnc1PerSepFirstGen[[;;,;;2]]}]];


(* ::Text:: *)
(*Figure 1h - correlation histogram for nuclear and pericycle correlations*)


Export[finalFiguresFolder <> "/fig1h_correlation_per "<>DateString[]<>".pdf",
Histogram[Correlation[normalize@#[[6,;;,2]],normalize@#[[6,;;,5]]]&/@Select[dConfnc1PerSepFirstGen, #[[8]]!= "N"&],30,##]&@@style2];


(* ::Subsection:: *)
(*Figure 2*)


(* ::Text:: *)
(*Figure 2e - example of trajectory of LS*)


(*Make directory *)
equidir = finalFiguresFolder <> "/fig2e-LS cell files FT - normalized" <>DateString[];
CreateDirectory[equidir];

(* Plot and save*)
Table[Block[{synf1d = Reverse@SortBy[GatherBy[{StringTake[#[[3]],{5}],#[[-1]]/."A"-> Orange/."S"->Black/."N"-> White, #[[6]]}&@Join[#[[;;5]],#[[6,-1]],{#[[8]]}]&/@Cases[dLSSep,{cf[[1]],cf[[2]],__}],First],#[[1,1]]&],
cfile = (Reverse@SortBy[#,ToExpression@StringTake[#[[1,3]],{5}]&]&@GatherBy[Cases[dLS,{cf[[1]],cf[[2]],__}],
		ToExpression@StringTake[#[[3]],{5}]&]),maxx,maxy},
maxx= Flatten[cfile[[;;,;;,6,;;,1]]]//Max;
maxy= 1.1;
Export[equidir<>"/"<>StringRiffle@cf<>".pdf",#]&@GraphicsColumn[#,ImageSize->defaultImageSize,Spacings->{0,-70},ImagePadding->10]&@
	Table[Show[{ListLinePlot[{#[[1]],normalize@#[[2]]}\[Transpose]&@(#\[Transpose])&/@(q[[1,;;,6,;;,;;2]]),##]&@@q[[2]],
				ListLinePlot[{#[[1]],normalize@#[[2]]}\[Transpose]&@(#\[Transpose])&/@(q[[1,;;,6,;;,{1,3}]]),##,ImageSize->defaultImageSize]&@@q[[3]]},PlotRange->{{0,maxx},{0,1.05maxy}}],{q,{cfile,style1dLS[[;;Length[synf1d]]],style1dbLS[[;;Length[synf1d]]],synf1d}\[Transpose]}]],{cf,Cases[DeleteDuplicates[dLSnc1[[;;,;;2]]],{"exp00481"|"exp00483","Upperright"|"upperleft"}]}];


(* ::Text:: *)
(*Figure 2f - Michalis Menten fit*)


Export[finalFiguresFolder <> "fig2f-MMFit.pdf",Show[{ListLinePlot[dataLSdata[[;;,{1,2}]],PlotStyle->gg],ListLinePlot[dataLSdata[[;;,{1,3}]],PlotStyle->mm],
Plot[MMfit,{t,0.1,35},PlotStyle->Gray]}]];


(* ::Text:: *)
(*Figure 2g- Hill fit*)


Export[finalFiguresFolder <> "fig2g-HillFit"<>ToString@#[[1]]<>".pdf",Show[{ListLinePlot[dataLSdata[[;;,{1,2}]],PlotStyle->gg],ListLinePlot[dataLSdata[[;;,{1,3}]],PlotStyle->mm],
Plot[#[[2]],{t,0.1,35},PlotStyle->Gray]}]]&/@MapIndexed[{#2[[1]],#1}&,Hillfit];


(* ::Text:: *)
(*Figure 2h - Feedback fit*)


Export[finalFiguresFolder <> "fig2h-linearFeedback.pdf",Show[{ListLinePlot[dataLSdata[[;;,{1,2}]],PlotStyle->gg],ListLinePlot[dataLSdata[[;;,{1,3}]],PlotStyle->mm],
Plot[sfeedhillfit,{t,0.1,35},PlotStyle->Gray]}]];


(* ::Subsection:: *)
(*Figure 3*)


(* ::Text:: *)
(*Figure 3b and 3c - trajectories of cells marked down in 3a*)


Table[Export[finalFiguresFolder<>"fig3b "<>q[[3]]<>DateString[]<>".pdf",ListLinePlot[{q[[6,;;,{1,2}]],q[[6,;;,{1,3}]]},PlotRange->{All,{0,3}},PlotStyle->{{gg},{mm}},Ticks->{{5,10,15,20},{0,0.5,1,1.5,2,2.5,3}},##]&@@style3d];,{q,dexp160}];


(* ::Text:: *)
(*Figure 3d - threshold of non-separated trajectories*)


(*calculate ratio between mean confocal -SHR and mean LS SHR of fully induced roots *)
factor = Mean@Flatten[Select[wnConf,#[[4]]==10&][[;;,6,;;,2]]]/Mean@Flatten[Select[wnLS,#[[4]]==40.&][[;;,6,;;,2]]];
(* Convert LS to have comparable units*)
wnLS3d={#1,#2,#3,#4,#5,{#1,factor #2,factor #3,#4}&@@@#6,#7,#8,#9}&@@@wnLS;
(* Prepare data for figure*)
dat4fig3d=(Flatten@thresholds[#[[1]],1,#[[2]],rangeLS,False]&/@{{wnConf,2},{wnLS3d,2},{wnLS3d,3}});
(* plot figure *)
Export[finalFiguresFolder<>"fig3d "<>DateString[]<>".pdf",ListLinePlot[{rangeLS,#}\[Transpose]&/@dat4fig3d,
			PlotRange->All,PlotStyle->{{Lighter@gg},{gg},{mm}},PlotLegends->{"Confocal SHR","Lightsheet SHR","Lightsheet SCR"},##]&@@style3d];


(* ::Text:: *)
(*Figure 3f - bar plot of threshold for separated trajectories*)


Export[finalFiguresFolder<>"/fig3f-barplot_"<>DateString[]<>".pdf",BarChart[dataforBars\[Transpose],ChartStyle->{Lighter@gg,gg,mm},ChartLegends->{"Confocal SHR","Lightsheet SHR","Lightsheet SCR"},ChartLabels->{{0.25,0.5,0.75,1},{"","",""}},LabelStyle->Large,BarSpacing->Automatic,##]&@@style2];


(* ::Text:: *)
(*Figure 3g and 3h - examples of for asymmetric and symmetric divisions and their cell cycle window shr scr threshold reaching*)


dat4choosing = Select[wnLSSep,#[[8]]==0 && Or@@Thread[#[[7,;;,-1]] < 0.2] && Or@@Thread[#[[7,;;,2]] > 0]&& Or@@Thread[#[[7,;;,3]] > 0]& ];


(*Choose cells *)
c1=Cases[wnLSSep,{"exp00488","Upperright","cell5b",__}][[1]];
c2=Cases[wnLSSep,{"exp00454","left","cell4",__}][[1]];
(*figure 3g*)
Export[finalFiguresFolder<>"/fig3g_divExample_"<>DateString[]<>".pdf",	
			ListLinePlot[{c1[[7,;;,{1,2}]],c1[[7,;;,{1,3}]],c1[[7,;;,{1,-1}]]},PlotStyle->({#}&/@{gg,mm,Black}),PlotRange->All,
			Prolog->{{Gray,Opacity[0.2],EdgeForm[],Rectangle[{0,0},{#,2}]&@Evaluate@FirstCase[c1[[7,;;,{1,-1}]],{_,_?(#>0.25&)}][[1]]}, 
													 {Purple,Opacity[1],Line[{{#,0},{#,2}}]&@Evaluate@FirstCase[c1[[6,;;,{1,2}]],{_,_?(#>thLSSep&)}][[1]]},
													 {Orange,Opacity[1],Line[{{#,0},{#,2}}]&@Evaluate@FirstCase[c1[[6,;;,{1,3}]],{_,_?(#>thLSSepSCR&)}][[1]]}},ImageSize->defaultImageSize,AxesStyle->framestyle]
	
	];

(*figure 3h*)
Export[finalFiguresFolder<>"/fig3h_nondivExample_"<>DateString[]<>".pdf",	
			ListLinePlot[{c2[[7,;;,{1,2}]],c2[[7,;;,{1,3}]],c2[[7,;;,{1,-1}]]},PlotStyle->({#}&/@{gg,mm,Black}),PlotRange->All,
			Prolog->{{Gray,Opacity[0.2],EdgeForm[],Rectangle[{0,0},{#,2}]&@Evaluate@FirstCase[c2[[7,;;,{1,-1}]],{_,_?(#>0.25&)}][[1]]}, 
													 {Purple,Opacity[1],Line[{{#,0},{#,2}}]&@Evaluate@FirstCase[c2[[6,;;,{1,2}]],{_,_?(#>thLSSep&)}][[1]]},
													 {Orange,Opacity[1],Line[{{#,0},{#,2}}]&@Evaluate@FirstCase[c2[[6,;;,{1,3}]],{_,_?(#>thLSSepSCR&)}][[1]]}},ImageSize->defaultImageSize,AxesStyle->framestyle]
	
	];


(* ::Text:: *)
(*Figure 3i -difference in sizes at first timepoints divides by fate*)


InitPlotLS   = Block[{mi=#[[1,1]],ma=#[[1,2]]},(#[[2]]-mi)/(ma-mi) 0.9]&@{#[[;;3]]/.ruleforlsnorm,#[[6,1,4]]}&/@#&/@(SortBy[#,#[[1,8]]&]&@GatherBy[Select[dLSnc1Sep  ,StringLength[#[[3]]]==5 && #[[8]]!= "N"&],#[[8]]&])[[{2,1}]];
InitPlotConf = Block[{mi=#[[1,1]],ma=#[[1,2]]},(#[[2]]-mi)/(ma-mi) 0.9]&@{#[[;;3]]/.ruleforNSnorm,#[[6,1,3]]}&/@#&/@(SortBy[#,#[[1,8]]&]&@GatherBy[Select[dConfnc1Sep,StringLength[#[[3]]]==5 && #[[8]]!= "N"&],#[[8]]&])[[{2,1}]];
Export[finalFiguresFolder<>"fig3i-difference in sizes "<>DateString[]<>".pdf",BoxWhiskerChart[{InitPlotLS,InitPlotConf},Frame->True,FrameStyle->Directive[Thickness[0.01`],GrayLevel[0],30],ChartLabels -> {{"LightSheet","Confocal"},{"","",""}},AspectRatio->1]];


(* ::Text:: *)
(*Figure 3j -bottom - chart of cell division (genealogy)*)


(*prepare all charts*)
graRess=Table[Block[{tit= h[[1]],dat=h[[2]],tt},
tt= {ToExpression@StringReplace[#[[1,1]],"cell"->""],{(ToExpression@StringReplace[StringReplace[#2,{#1->"0","aa"->"-3","ab"-> "-1","ba"->"+1","bb"-> "+3"}],{"a"-> "-2","b"->"+2"}]),#4[[2]],#4[[1]]}&@@@#}&/@GatherBy[dat,#[[1]]&];

tt= {#1,Prepend[#2,{0,0,Black}]}&@@@tt;

tt=Table[{t[[1]]*7,{t[[2,;;,2]],t[[2,;;,1]],t[[2,;;,3]]}\[Transpose]},{t,tt}];

tt={#[[1]],Cases[Select[Subsets[#,{2}],(#[[1,1]]!=0 || (#[[1,1]]==0 && #[[1,2]]==#[[2,2]]))&],{{_,0,_},{_,-2,_}}|{{_,0,_},{_,2,_}}|{{_,2,_},{_,1,_}}|{{_,2,_},{_,3,_}}|{{_,-2,_},{_,-1,_}}|{{_,-2,_},{_,-3,_}}|{{0,0,_},{_,0,_}}]&@#[[2]]}&/@tt;

tt=Map[{Black,Line[{#1,{#1[[1]],#2[[2]]},{#2[[1]],#2[[2]]},#2}&@@#[[;;,;;2]]],#[[1,3]],PointSize[Large],Point@{#[[1,;;2]]},#[[2,3]],PointSize[Large],Point@{#[[2,;;2]]}}&,Table[Block[{t7=t[[1]],rt=t[[2]]},Transpose/@Transpose[{rt[[;;,;;,1]],rt[[;;,;;,2]]+t7,rt[[;;,;;,3]]}]],{t,tt}],{2}];

Graphics[tt,Frame->True,PlotLabel->Style[StringRiffle@tit,8],ImageSize->defaultImageSize,FrameTicks->{{None,None},{{0,5,10,15,20,25,30,35,40,45},None}}]
],{h,({#[[1,;;2]],#[[;;,3;;-1]]}&/@GatherBy[divOc,#[[1]]&&#[[2]]&])}];

(*Export Figure*)
Export[finalFiguresFolder<>"/fig3j-hydroxyCharts "<>DateString[]<>".pdf",GraphicsRow[graRess[[{8,2}]],ImageSize->defaultImageSize]];


(* ::Subsection:: *)
(*Figure S2*)


(* ::Text:: *)
(*Figure s2b - all cell + mean trajectory split by dex concentration*)


Export[finalFiguresFolder<>"/figs2b-mean behavior by dex "<>DateString[]<>".pdf",#]&@GraphicsRow[#,ImageSize->defaultImageSize,ImagePadding->10]&@Table[
			Show[{ListLinePlot[q[[;;,7,;;,;;2]],AspectRatio->1.5,PlotRange->{{0,Max@Flatten@dConf[[;;,7,;;,1]]},{0,1.05Max@Flatten@dConf[[;;,7,;;,2]]}},##]&@@style1c,
					ListLinePlot[TakeWhile[{#[[1,1]],Mean@#[[;;,2]],Length@#,Length@q}&/@GatherBy[SortBy[Flatten[q[[;;,6,;;,;;2]],1],#[[1]]&],
	   Round[#[[1]],0.25]&],#[[3]]>0.3 #[[4]]&][[;;,;;2]],##]&@@style1ca}],{q,Reverse@SortBy[#,#[[1,4]]&]&@GatherBy[dConf,#[[4]]&]}];


(* ::Text:: *)
(*Figure s2c - Matrices of plots split by dex conentration and division and cell position*)


Export[finalFiguresFolder<> "figS2c -  matrix  cells" <> " "<> DateString[]<>".pdf",(GraphicsGrid[(*Show/@#&/@*)Table[Block[
	{minidat = Select[dConf,ToExpression@StringTake[#[[3]],{5}]==pos && #[[4]]==dex &],
	color= Gray ,(*If[div==1,Gray,Pink]*),
	colormean=Black (*If[div==1,Black,Red]*)},
	Show[{ListLinePlot[minidat[[;;,6,;;,;;2]],PlotRange->{{0,24},{0,0.4}},AspectRatio->1,PlotStyle->{{color,Opacity[0.3]}}],
		  ListLinePlot[SortBy[Mean/@Select[GatherBy[Flatten[minidat[[;;,6,;;,;;2]],1],Round[#[[1]],0.25]&],Length@#>0.3 Length@minidat&],#[[1]]&],PlotRange->All,PlotStyle->colormean]}]
],
{pos,{5,4,3,2,1}},{dex,Reverse@{0.01,.02,0.03,0.05,1,10}}]])];


(* ::Text:: *)
(*Figure s2d - Time to division by cell position *)


Export[finalFiguresFolder<> "figS2d - time to division by cell position" <> " "<> DateString[]<>".pdf",
Show[{BoxWhiskerChart[#,ChartLabels->{"cell1","cell2","cell3","cell4","cell5"},ChartStyle->Gray,PlotRange->All,AspectRatio->1],
	  ListPlot[#,PlotStyle->(ColorData[16]/@{1,2,3,4,5}),PlotRange->All,AspectRatio->1]&@MapIndexed[Block[{num=#2[[1]]},{num,#}&/@#1]&,#]}]&@Table[Block[
	{minidat = Select[d10,#[[8]]==1 && ToExpression@StringTake[#[[3]],{5}]==pos  &]},
	(Sort/@minidat[[;;,7,;;,1]])[[;;,-1]]
],
{pos,5}]];


(* ::Text:: *)
(*Figure s2e- percent of division by cell position *)


Export[finalFiguresFolder<> "figS2e - percentage of division by cell position" <> " "<> DateString[]<>".pdf",
BarChart[#,ChartLabels->{"cell1","cell2","cell3","cell4","cell5"},ChartStyle->Gray,PlotRange->All,AspectRatio->1]&@Table[Block[
	{minidat = Select[d10,ToExpression@StringTake[#[[3]],{5}]==pos  &]},
	Count[#,1]/Length@#&@minidat[[;;,8]]
],
{pos,5}]];


(* ::Text:: *)
(*Figure s2f - SHR levels by dex *)


Export[finalFiguresFolder<> "figS2f - SHR by dex" <> " "<> DateString[]<>".pdf",
Show[{BoxWhiskerChart[#,ChartLabels->Reverse@{"0.01","0.02","0.03","0.05","1","10"},ChartStyle->Gray,PlotRange->All,AspectRatio->1],
	  ListPlot[#,PlotStyle->(ColorData[16]/@{1,2,3,4,5,6}),PlotRange->All]&@MapIndexed[Block[{num=#2[[1]]},{num,#}&/@#1]&,#]}]&@Table[Block[
	{minidat = Select[dConfnc1, #[[4]]==dex  &]},
	(Quantile[#,0.9]&/@minidat[[;;,6,;;,2]])
],
{dex,Reverse@{0.01,0.02,0.03,0.05,1,10}}]];


(* ::Text:: *)
(*Figure s2g - percent of division by dex *)


Export[finalFiguresFolder<> "figS2g - percent of division by dex" <> " "<> DateString[]<>".pdf",
BarChart[#,ChartLabels->Reverse@{"0.01","0.02","0.03","0.05","1","10"},ChartStyle->Gray,PlotRange->All,AspectRatio->1]&@Table[Block[
	{minidat = Select[dConfnc1, #[[4]]==dex  &]},
	Count[#,1]/Length@#&@minidat[[;;,8]]
],
{dex,Reverse@{0.01,0.02,0.03,0.05,1,10}}]];


(* ::Text:: *)
(*Figure s2i - cell duration difference between asymmetric and symmetric divisions*)


Export[finalFiguresFolder<> "figS2i_duration_cell_cycle.pdf",BoxWhiskerChart[#[[;;,1]],ChartLabels->#[[;;,2]]]&@({#[[;;,6,-1,1]]-#[[;;,6,1,1]],#[[1,8]]}&/@GatherBy[Select[dConfnc1Sep,StringLength@#[[3]]>5&],#[[8]]&])[[{1,3}]]];


(* ::Subsection:: *)
(*Figure S4*)


Export[finalFiguresFolder<>"/figS4 - hydroxy "<>DateString[]<>".pdf",GraphicsGrid[GatherBy[graRess,StringSplit[#[[3,2,1]]][[1]]&][[{2,4,5,1,3,6}]],ImageSize->1000,Alignment->Left]];


(* ::Section:: *)
(*Numbers from paper*)


number4Paper={};


AppendTo[number4Paper, {"number of lineages in Confocal",Length@dConf}];
AppendTo[number4Paper, {"number of roots in Confocal",dConf[[;;,1]]//Union//Length}];
AppendTo[number4Paper, {"Percentage asym div in 1mM Dex",100. Total[#]/Length@#&@(Select[dConfnc1,#[[4]]==1&][[;;,-2]])}];
AppendTo[number4Paper, {"Percentage asym div in 10mM Dex",100. Total[#]/Length@#&@(Select[dConfnc1,#[[4]]==10&][[;;,-2]])}];
AppendTo[number4Paper,{"Mean Correlation between per and shr at the beginning of TC - a-div",Correlation[normalize@#[[6,;;,2]],normalize@#[[6,;;,5]]]&/@Select[dConfnc1PerSep,StringLength@#[[3]] == 5 && #[[8]]== "A"&]//Mean}];
AppendTo[number4Paper, {"First asym div in LightSheet",Min[Select[Select[dLSnc1,#[[4]]==40.&],#[[8]]==1&][[;;,-1]]]}];
AppendTo[number4Paper, {"Percentage asym div in before 48 LightSheet",Count[Select[Select[dLSnc1,#[[4]]==40.&],#[[8]]==1&][[;;,-1]],_?(#<48&)]/Length@Select[dLSnc1,#[[4]]==40.&]*100//N}];
AppendTo[number4Paper,{"adj R2 MM model",r2AM}];
AppendTo[number4Paper,{"adj R2 Hill model",r2AH}];
AppendTo[number4Paper,{"adj R2 FB model",r2AF}];
AppendTo[number4Paper,{"Max threshold curve LS SHR",Max[dat4fig3d[[2]]]}];
AppendTo[number4Paper,{"Max threshold curve LS SCR",Max[dat4fig3d[[3]]]}];
AppendTo[number4Paper,{"Max threshold curve Confocal",Max[dat4fig3d[[1]]]}];
AppendTo[number4Paper, {"number of features Confocal",Dimensions[featuresConf][[2]] - 7}];
AppendTo[number4Paper, {"number of features LightSheet",Dimensions[featuresLS][[2]] - 7}];
AppendTo[number4Paper,{#[[1]],#[[2]]}]&/@({Flatten[Table["Sep Max threshold curve for " <>b <>" in window size:" <> a,{b,{"Conf","LS-SHR","LS-SCR"}},{a,{"0.25"}}]],dataforBars[[;;,1]]}\[Transpose]);
AppendTo[number4Paper,{"Pval: Mann-Whitney test for LS Initial sizes symmetric vs. asym.",MannWhitneyTest[InitPlotLS[[;;2]]]}];
AppendTo[number4Paper,{"Pval: Mann-Whitney test for Confocal Initial sizes symmetric vs. asym.",MannWhitneyTest[InitPlotConf[[;;2]]]}];
AppendTo[number4Paper,{"MannWhitney cell cycle duration diff.",MannWhitneyTest[({#[[;;,6,-1,1]]-#[[;;,6,1,1]],#[[1,8]]}&/@GatherBy[Select[dConfnc1Sep,StringLength@#[[3]]>5&],#[[8]]&])[[{1,3},1]]]}];


(* ::Section:: *)
(*Export Tables *)


(* ::Text:: *)
(*Export number tables*)


Export[finalFiguresFolder<>"numbers_table "<>DateString[]<>".csv",number4Paper];


(* ::Text:: *)
(*Export features*)


Export[#[[1]]<>"/features "<>DateString[]<>".csv",Prepend[#[[3]],#[[2]]]]&/@({subpaths,ts,features}\[Transpose]);


(* ::Text:: *)
(*Export normalized and non-normalized data*)


Prepend[Flatten[Table[Table[Flatten@Join[l[[1,;;5]],N@Flatten@v,l[[1,8;;]]],{v,{l[[1,{6,7}]]\[Transpose],l[[2,7,;;,-1]],N@normalize@l[[2,7,;;,-1]]}\[Transpose]}],{l,{DeleteCases[dConfnc1,{___,"N",___}],wnConf}\[Transpose]}],1],      {"Exp","file","cell","dex","ind","time","smoothed shr","smoothed nuclear size","time", "shr","nuclear size","norm NS FT","norm NS sep","div","time to div"}]//Export[subpaths[[1]]<>"/Data "<> DateString[]<> ".csv",#]&;
Prepend[Flatten[Table[Table[Flatten@Join[l[[1,;;5]],N@Flatten@v,l[[1,8;;]]],{v,{l[[1,{6,7}]]\[Transpose],l[[2,7,;;,-1]],N@normalize@l[[2,7,;;,-1]]}\[Transpose]}],{l,{DeleteCases[dConfnc1Sep,{___,"N",___}],wnConfSep}\[Transpose]}],1],{"Exp","file","cell","dex","ind","time","smoothed shr","smoothed nuclear size","time", "shr","nuclear size","norm NS FT","norm NS sep","div","time to div"}]//Export[subpaths[[2]]<>"/Data "<> DateString[]<> ".csv",#]&;									
Prepend[Flatten[Table[Table[Flatten@Join[l[[1,;;5]],N@Flatten@v,l[[1,8;;]]],{v,{l[[1,{6,7}]]\[Transpose],l[[2,7,;;,-1]],N@normalize@l[[2,7,;;,-1]]}\[Transpose]}],{l,{DeleteCases[dLSnc1,{___,"N",___}],wnLS}\[Transpose]}],1],          {"Exp","file","cell","dex","ind","time","smoothed shr","smoothed scr","smoothed nuclear size","time", "shr","scr","nuclear size","norm NS FT","norm NS sep","div","time to div"}]//Export[subpaths[[3]]<>"/Data "<> DateString[]<> ".csv",#]&;
Prepend[Flatten[Table[Table[Flatten@Join[l[[1,;;5]],N@Flatten@v,l[[1,8;;]]],{v,{l[[1,{6,7}]]\[Transpose],l[[2,7,;;,-1]],N@normalize@l[[2,7,;;,-1]]}\[Transpose]}],{l,{DeleteCases[dLSnc1Sep,{___,"N",___}],wnLSSep}\[Transpose]}],1],    {"Exp","file","cell","dex","ind","time","smoothed shr","smoothed scr","smoothed nuclear size","time", "shr","scr","nuclear size","norm NS FT","norm NS sep","div","time to div"}]//Export[subpaths[[4]]<>"/Data "<> DateString[]<> ".csv",#]&;


(* ::Text:: *)
(*Export feature scores*)


featureTable=Table[ Block[{temp},
temp = (Reverse@SortBy[
	   (If[MemberQ[f[[2,;;,1]],#1[[1]]],Append[Join[#1[[;;4]],{#2,#3}],"high lentgh correlation"],Append[Join[#1[[;;4]],{#2,#3}],""]]&@@@({f[[3]],f[[4]],f[[5]]}\[Transpose])),#[[3]]&]);

Export[f[[1]]<>"/oneByoneFeatures "<>DateString[]<>".csv",temp];
temp],
	   
{f,{subpaths,corLengths,onebyoneScores,pvals,FDRs}\[Transpose][[;;]]}];


Table[
Export[f[[1]]<>"/oneByoneFeatures short"<>DateString[]<>".csv",
	  Select[f[[2]],#[[-1]]==""&& #[[3]]>=0.75  && #[[6]]<0.05&] ],
	   
{f,{subpaths,featureTable}\[Transpose][[{1,3}]]}];

Table[
Export[f[[1]]<>"/oneByoneFeatures short"<>DateString[]<>".csv",
	  Select[f[[2]], #[[3]]>=0.75  && #[[6]]<0.05&] ],
	   
{f,{subpaths,featureTable[[;;,;;,;;-2]]}\[Transpose][[{2,4}]]}];


Export[finalFiguresFolder<>"feature names.csv",DeleteDuplicates[Flatten[ts]]];
```
