## Extended Data Tables for "Patterning and growth are coordinated early in the cell cycle"

**Extended Data Table 1.** Predictive features from the light sheet data utilizing full trajectories (see Methods).

| **Feature** | **Predictive Accuracy*** | **p-value (Mann-Whitney)** | **FDR** |
| --- | --- | --- | --- |
| std - shr | 85.0% | 2.07E-11 | 1.86E-10 |
| quantile 0.95 - shr | 84.4% | 1.80E-08 | 8.09E-08 |
| quantile 0.5 - shr | 84.4% | 9.89E-08 | 3.90E-07 |
| first time at quantile 0.5 - shr | 83.8% | 2.61E-04 | 5.87E-04 |
| mean - shr | 83.8% | 2.24E-07 | 7.85E-07 |
| cor shr / scr | 82.5% | 5.45E-15 | 5.73E-14 |
| shr at max rate - scr | 81.3% | 1.21E-09 | 6.93E-09 |
| max rate - shr | 81.3% | 7.88E-07 | 2.48E-06 |
| mean - nuclear size | 80.0% | 5.09E-18 | 1.60E-16 |
| norm nuclear size at shr threshold | 79.4% | 1.01E-09 | 7.05E-09 |
| quantile 0.7 - nuclear size | 79.4% | 4.59E-16 | 5.78E-15 |
| scr at max rate - norm nuclear size | 79.4% | 7.26E-05 | 1.83E-04 |
| quantile 0.5 - nuclear size | 78.8% | 1.95E-16 | 4.10E-15 |
| quantile 0.3 - nuclear size | 78.8% | 2.57E-16 | 4.05E-15 |
| time at max rate - norm nuclear size | 78.8% | 2.19E-09 | 1.15E-08 |
| time at max rate - nuclear size | 78.8% | 2.26E-09 | 1.10E-08 |
| first time at quantile 0.95 - norm nuclear size | 77.5% | 4.33E-04 | 9.10E-04 |
| first time at quantile 0.95 - nuclear size | 77.5% | 4.33E-04 | 9.41E-04 |
| nuclear size at max rate - norm nuclear size | 77.5% | 7.92E-03 | 1.47E-02 |
| quantile 0.3 - norm nuclear size | 76.9% | 4.38E-03 | 8.36E-03 |
| quantile 0.95 - scr | 76.9% | 9.19E-05 | 2.15E-04 |
| nuclear size at max rate - scr | 76.9% | 1.18E-09 | 7.42E-09 |
| first time at quantile 0.5 - scr | 76.3% | 2.61E-08 | 1.10E-07 |
| std - scr | 76.3% | 1.40E-06 | 4.02E-06 |
| AUC - scr | 76.3% | 7.56E-05 | 1.83E-04 |
| shr at max rate - nuclear size | 76.3% | 2.67E-05 | 7.01E-05 |
| max rate - scr | 76.3% | 1.14E-03 | 2.24E-03 |
| nuclear size at max rate - shr | 76.3% | 1.91E-07 | 7.08E-07 |
| quantile 0.5 - scr | 75.6% | 2.64E-02 | 4.38E-02 |
| mean - norm nuclear size | 75.0% | 9.38E-03 | 1.64E-02 |

* Predictive accuracy reflects the ability of a discrimination model to separate the data into asymmetrically divided vs. undivided cells. Only significant (FDR < 0.001) features with a predictive accuracy > 75% are shown.

**Extended Data Table 2.** Predictive features from the confocal data utilizing full trajectories.

| **Features** | **Predictive Accuracy** | **p-value (Mann-Whitney)** | **FDR** |
| --- | --- | --- | --- |
| quantile 0.5 - shr | 90.8% | 6.69E-90 | 3.01E-88 |
| shr at max rate - nuclear size | 87.0% | 5.63E-85 | 8.45E-84 |
| mean - shr | 85.4% | 2.47E-88 | 5.56E-87 |
| shr at max rate - shr | 85.4% | 2.24E-80 | 2.02E-79 |
| time from theshold to end | 85.1% | 1.84E-58 | 9.19E-58 |
| quantile 0.95 - shr | 85.1% | 2.18E-80 | 2.46E-79 |
| max rate - shr | 84.3% | 7.86E-78 | 5.89E-77 |
| AUC - shr | 81.6% | 8.31E-75 | 5.34E-74 |
| norm nuclear size at shr threshold | 79.7% | 1.99E-52 | 8.13E-52 |
| cv - shr | 79.7% | 1.87E-53 | 8.43E-53 |
| first time at quantile 0.5 - shr | 79.3% | 7.28E-38 | 2.52E-37 |
| std - shr | 77.4% | 7.91E-69 | 4.45E-68 |

* Predictive accuracy reflects the ability of a discrimination model to separate the data into asymmetrically divided and undivided cells. Only significant (FDR < 0.001) features with a predictive accuracy > 75% are shown.

**Extended Data Table 3.** Predictive features from the light sheet data utilizing separated trajectories (see Methods).

| **Feature** | **Predictive Accuracy*** | **p-value (Mann-Whitney)** | **FDR** |
| --- | --- | --- | --- |
| Max shr levels at norm nuclear window of 0.25 | 93.5% | 1.88E-64 | 1.79E-62 |
| AUC shr levels at norm nuclear window of 0.25 | 93.5% | 7.44E-63 | 2.36E-61 |
| Max shr levels at norm nuclear window of 0.5 | 91.0% | 2.44E-64 | 1.16E-62 |
| AUC shr levels at norm nuclear window of 0.5 | 91.0% | 1.07E-58 | 2.55E-57 |
| norm nuclear size at shr threshold | 90.0% | 1.57E-51 | 1.87E-50 |
| AUC scr levels at norm nuclear window of 0.25 | 89.5% | 9.00E-56 | 1.42E-54 |
| Max scr levels at norm nuclear window of 0.25 | 86.5% | 1.99E-57 | 3.78E-56 |
| Max scr levels at norm nuclear window of 0.75 | 86.0% | 1.45E-47 | 1.37E-46 |
| norm nuclear size at scr threshold | 85.5% | 6.00E-41 | 3.80E-40 |
| std - norm nuclear size | 84.0% | 8.04E-36 | 3.64E-35 |
| Max shr levels at norm nuclear window of 0.75 | 82.5% | 2.12E-48 | 2.24E-47 |
| AUC shr levels at norm nuclear window of 0.75 | 82.0% | 5.29E-35 | 2.18E-34 |
| norm nuclear size at first shr level of 0.02 | 81.0% | 1.01E-28 | 3.32E-28 |
| cv - norm nuclear size | 81.0% | 2.78E-45 | 2.20E-44 |
| quantile 0.95 - scr | 80.5% | 1.24E-35 | 5.34E-35 |
| quantile 0.5 - scr | 80.5% | 6.79E-45 | 4.61E-44 |
| AUC scr levels at norm nuclear window of 0.75 | 79.5% | 6.27E-35 | 2.48E-34 |
| AUC - scr | 79.5% | 1.84E-39 | 1.09E-38 |
| AUC scr levels at norm nuclear window of 0.5 | 79.0% | 2.13E-47 | 1.84E-46 |
| norm nuclear size at first shr level of 0.05 | 79.0% | 3.88E-19 | 9.45E-19 |
| mean - scr | 79.0% | 5.11E-45 | 3.74E-44 |
| max rate - norm nuclear size | 78.5% | 5.81E-38 | 3.07E-37 |
| Max scr levels at norm nuclear window of 0.5 | 78.0% | 1.42E-52 | 1.93E-51 |
| scr at max rate - scr | 78.0% | 1.12E-38 | 6.26E-38 |
| mean - norm nuclear size | 77.5% | 1.10E-33 | 4.17E-33 |
| std - scr | 77.0% | 2.32E-22 | 6.48E-22 |
| shr at max rate - norm nuclear size | 77.0% | 8.83E-10 | 1.86E-09 |
| scr at max rate - nuclear size | 77.0% | 9.72E-31 | 3.55E-30 |
| norm nuclear size at first shr level of 0.075 | 76.5% | 6.38E-15 | 1.41E-14 |
| nuclear size at max rate - norm nuclear size | 76.5% | 2.20E-24 | 6.34E-24 |
| shr at max rate - nuclear size | 76.5% | 3.30E-29 | 1.12E-28 |
| scr at max rate - shr | 76.5% | 1.14E-27 | 3.48E-27 |
| quantile 0.5 - shr | 75.0% | 8.34E-38 | 4.17E-37 |
| AUC - shr | 75.0% | 6.71E-30 | 2.36E-29 |
| max rate - scr | 75.0% | 5.65E-20 | 1.41E-19 |

*Predictive accuracy reflects the ability of a discrimination model to separate the data into asymmetrically and symmetrically dividing cells. Only significant (FDR < 0.001) features with a predictive accuracy > 75% are shown.

**Extended Data Table 4.** Predictive features from the confocal data utilizing separated trajectories.

| **Features** | **Predictive Accuracy*** | **p-value (t-test)** | **p-value (Mann Whitney)** |
| --- | --- | --- | --- |
| Max shr levels at norm nuclear window of 0.25 | 88.5% | 6.92E-85 | 4.08E-83 |
| AUC shr levels at norm nuclear window of 0.25 | 86.2% | 4.35E-80 | 8.56E-79 |
| AUC shr levels at norm nuclear window of 0.5 | 85.4% | 3.07E-73 | 4.53E-72 |
| Max shr levels at norm nuclear window of 0.5 | 85.0% | 2.42E-82 | 7.13E-81 |
| time from theshold to end | 83.0% | 1.51E-55 | 7.42E-55 |
| Max shr levels at norm nuclear window of 0.75 | 81.8% | 1.48E-67 | 1.09E-66 |
| mean - shr | 81.4% | 3.97E-70 | 3.91E-69 |
| AUC - shr | 79.8% | 4.55E-68 | 3.83E-67 |
| quantile 0.5 - shr | 79.4% | 6.40E-72 | 7.55E-71 |
| shr at max rate - shr | 79.4% | 1.92E-62 | 1.26E-61 |
| quantile 0.95 - shr | 78.7% | 7.70E-60 | 4.13E-59 |
| shr at max rate - nuclear size | 78.7% | 3.19E-61 | 1.88E-60 |
| AUC shr levels at norm nuclear window of 0.75 | 78.3% | 3.65E-53 | 1.66E-52 |
| cv - norm nuclear size | 77.5% | 4.31E-53 | 1.82E-52 |
| std - norm nuclear size | 77.5% | 1.95E-33 | 4.99E-33 |
| mean - norm nuclear size | 77.1% | 5.30E-49 | 2.09E-48 |
| quantile 0.3 - norm nuclear size | 75.9% | 1.22E-46 | 4.50E-46 |
| std - shr | 75.9% | 2.34E-38 | 6.92E-38 |
| Max shr levels at norm nuclear window of 0.25 | 88.5% | 6.92E-85 | 4.08E-83 |
| AUC shr levels at norm nuclear window of 0.25 | 86.2% | 4.35E-80 | 8.56E-79 |
| AUC shr levels at norm nuclear window of 0.5 | 85.4% | 3.07E-73 | 4.53E-72 |
| Max shr levels at norm nuclear window of 0.5 | 85.0% | 2.42E-82 | 7.13E-81 |
| time from theshold to end | 83.0% | 1.51E-55 | 7.42E-55 |
| Max shr levels at norm nuclear window of 0.75 | 81.8% | 1.48E-67 | 1.09E-66 |
| mean - shr | 81.4% | 3.97E-70 | 3.91E-69 |
| AUC - shr | 79.8% | 4.55E-68 | 3.83E-67 |
| quantile 0.5 - shr | 79.4% | 6.40E-72 | 7.55E-71 |
| shr at max rate - shr | 79.4% | 1.92E-62 | 1.26E-61 |
| quantile 0.95 - shr | 78.7% | 7.70E-60 | 4.13E-59 |
| shr at max rate - nuclear size | 78.7% | 3.19E-61 | 1.88E-60 |
| AUC shr levels at norm nuclear window of 0.75 | 78.3% | 3.65E-53 | 1.66E-52 |
| cv - norm nuclear size | 77.5% | 4.31E-53 | 1.82E-52 |
| std - norm nuclear size | 77.5% | 1.95E-33 | 4.99E-33 |
| mean - norm nuclear size | 77.1% | 5.30E-49 | 2.09E-48 |
| quantile 0.3 - norm nuclear size | 75.9% | 1.22E-46 | 4.50E-46 |
| std - shr | 75.9% | 2.34E-38 | 6.92E-38 |

* Predictive accuracy reflects the ability of a discrimination model to separate the data into asymmetrically and symmetrically dividing cells. Only significant (FDR < 0.001) features with a predictive accuracy > 75% are shown.
